# Multi-Omics Profiling Reveals Gene Signatures and Therapeutic Targets in HER2- Guided Gastric Cardia Adenocarcinoma Patients

**DOI:** 10.1101/2024.03.02.583089

**Authors:** Li-Dong Wang, Pengwei Xing, Xue-Ke Zhao, Xin Song, Meng-Xia Wei, Duo You, Ling-Ling Lei, Rui-Hua Xu, Ran Wang, Lei Ma, Pei-Nan Chen, Xinmin Li, Minglu Xie, Miao Zhao, He Zhang, Hui-Fang Lv, Ai-Li Li, Xian-Zeng Wang, She-Gan Gao, Xingqi Chen

## Abstract

Therapeutic targets for Gastric Cardia Adenocarcinoma (GCA), particularly for HER2- negative patients, are lacking. Here, we conducted multi-omics profiling on 128 GCA patients using mass spectrometry, whole-exome sequencing, RNA-Seq, and metabolomics. We found HER2 to be a favorable prognostic marker for GCA. Employing molecular counting, we categorized patients into HER2-high, -low, and -negative groups. We uncovered an enrichment of DNA repair features in the HER2-high group, while HER2-low and -negative groups exhibited strong inflammation. We found that tumor mutation burden may not be the distinguishing factor among these three groups. We revealed that the HER2-negative and -low groups have a tumor-suppressive immune microenvironment, and HER2 expression is associated with fatty acid metabolic profiles and inflammation in the blood of patients. Our study revealed anti-inflammatory and immune checkpoint inhibition, targeting PD-L2 and the CD47/SIRPA pair, as therapeutic strategies for HER2-negative GCA patients. These findings highlight promising avenues for personalized treatment for GCA.

## Introduction

Gastric cardia adenocarcinoma(GCA) is a type II adenocarcinoma at the gastroesophageal junction ^1^. GCA exhibits distinct features compared to esophageal and gastric cancers ^2–5^, yet therapeutic approaches often overlap. Consequently, specific therapeutic targets for GCA are lacking. Human Epidermal Growth Factor Receptor 2 (HER2), the protein product of the Erb- B2 Receptor Tyrosine Kinase 2 (*ERBB2*) gene, plays a crucial role in regulating cell growth and division^6^. In certain cancers, such as breast cancer ^7,8^ and gastroesophageal cancer ^9^, HER2 can become overexpressed or amplified, leading to uncontrolled cell growth ^10^. Targeted therapies, such as trastuzumab (Herceptin) and others, have been developed to specifically inhibit HER2 signaling and are effective in treating HER2-positive breast ^11^ and gastroesophageal ^12^ cancers. Currently, HER2 targeted therapy has become a vital treatment option for certain types of gastroesophageal cancers, including esophageal Adenocarcinoma (EAC), Esophageal squamous-cell carcinoma (ESCC), GCA, and stomach adenocarcinoma (STAD), particularly in advanced or metastatic cases ^9,13–15^. Anti-HER2 targeted therapies have improved survival outcomes in advanced gastroesophageal cancer patients with HER2 overexpression ^13,14,16^. However, the relatively low prevalence (less than 30%) of HER2- positive cases poses challenges ^17–19^, particularly for HER2-negative patients. The first-line treatment for advanced HER2-negative gastric cancer and esophagogastric junction adenocarcinoma mainly involves immunotherapy combined with chemotherapy, without specific targets^9^. In solid malignancies, the overexpression or amplification of HER2 is associated with an adverse prognosis ^20,21^, paradoxically, independent studies including our recent study, have shown that *ERBB2* gene focal amplification or HER2 overexpression might confer a favorable prognostic impact in patients with esophagogastric junction adenocarcinoma, EAC ^22^ and GCA ^23^. Hence, the role of HER2 as either a therapeutic target or a prognostic marker in GCA remains uncertain. Consequently, conducting a comprehensive multi-omics molecular profiling of GCA patients, stratified by the degree of HER2 expression, is imperative for elucidating this paradox and gaining a more profound understanding that can inform targeted therapy strategies for these malignancies, particularly the HER2-negative patients.

In this investigation, we assembled a cohort comprising 128 patients diagnosed with GCA. We conducted an extensive array of analyses on our GCA cohort, including RNA-Seq for transcriptome profiling, whole exome sequencing (WES) for examining genetic alterations, mass spectrometry for the identification and quantification of proteins, single-cell RNA-Seq (scRNA-Seq), and spatial transcriptomics for dissecting the tumor microenvironment. With a highly sensitive molecular counting approach, we stratified GCA patients into HER2-high, - low, and -negative groups. This method enabled us to undertake a comprehensive and detailed characterization of molecular features for each patient group, including the exploration of gene expression patterns, profiling of proteins, and examination of genetic alterations. We found HER2 as a favorite prognostic marker in GCA and EAC, but not in gastric cancers. Furthermore, our investigations unveiled a substantial enrichment of DNA repair pathways in the HER2 high-expression group, while the HER2 low-expression and negative group prominently exhibited features indicative of inflammation. We found that tumor mutation burden (TMB) may not be the primary distinguishing factor among these three groups, but the HER2 high group showed substantially higher genome instability compared with the HER2 negative group. We also found *ARID1A* mutations were significant for prognosis, especially in the HER2-negative group. Notably, scRNA-Seq and spatial transcritons revealed the HER2- high group displayed stronger growth factor signaling, whereas the HER2-low and HER2- negative groups showed pronounced inflammation and higher M2 macrophage levels, and variations in immune inhibitory checkpoints across the distinct HER2 expression groups. With validation from multiple angles, our study suggests that anti-inflammatory therapy and immune checkpoint inhibition targeting the CD47/SIRPA axis may be potential therapeutic strategies for HER2-negative GCA patients.

## Results

### HER2/*ERBB2* is a favorable prognostic marker in GCA and EAC

Previously, we and others found that *ERBB2* gene focal amplification or HER2 overexpression might confer a favorable prognostic impact on patients with esophageal adenocarcinoma (EAC) and GCA ^22,23^. This conclusion is derived from immunohistochemistry (IHC) staining of HER2 or gene amplification detection using whole-genome sequencing. These measurements involve positive and negative HER2 staining or amplification assessments of patients. The HER2 scoring system typically employs immunohistochemistry (IHC) and/or fluorescence in situ hybridization (FISH) techniques to evaluate the quantity and distribution of HER2 protein and gene amplification in cancer tissue samples ^24^. In the whole-genome sequencing method, HER2 amplification is examined within amplification regions^24^. It is well known there is a big heterogeneous expression of HER2 protein in different regions of the tumor from the same patient ^17^. RNA-Seq is a molecular counting method used to calculate the number of transcripts. To better understand the relationship between HER2 expression and patient prognosis in GCA, we performed RNA-seq and mass spectrometry on both tumors and NATs from 54 GCA patients (**Supplementary Figure 1a-c**). We observed a strong correlation between *ERBB2* RNA expression levels and HER2 protein levels in tumors (Pearson correlation = 0.7559), but amost no correlation in NATs (Pearson correlation = -0.03) (**Figure 1a**). Based on the median expression level of *ERBB2* RNA, we stratified GCA patients into high and low groups (**Figure 1b**, **Supplementary Figure 2)**. We also analyzed RNA-Seq data from TCGA for EAC, ESCC, and STAD (Stomach adenocarcinoma), stratifying patients into HER2-high and HER2-low groups based on the median *ERBB2* RNA expression levels. Significant differences in survival probabilities were observed among GCA (*p* = 0.011) and EAC (*p* = 0.0096) patients (**Figure 1c**). Specifically, the *ERBB2* high group demonstrated significantly better survival probabilities compared to the *ERBB2* low group. No significant survival difference was found in ESCC (*p* = 0.59) or STAD (*p* = 0.57). Further analysis within STAD subtypes revealed no difference in diffuse or intestinal carcinoma types (**Figure 1c**). Although not statistically significant, there was a trend for better survival in the *ERBB2* high group for STAD tubular adenocarcinoma (*p* = 0.19) and STAD NOS adenocarcinoma (*p* = 0.4) (**Figure 1c**). We also used quantile and natural separation as cutoffs for patient grouping (**Supplementary Figure 2**). In the quantile grouping, we still observed that high levels of HER2 RNA expression were associated with better prognosis in GCA (*p* = 0.049) and EAC (*p* = 0.05), but not in ESCC or STAD, further indicating that HER2 could be a potential prognostic marker for GCA and EAC. Considering genetic background variations, we also calculated the HER2 RNA level ratio between tumors and the corresponding NATs and divide patients into high and low groups (**Methods, Figure 1d**), we still observed significantly better survival in the HER2 high group (*p* = 0.021, **Figure 1e**). Next, we expanded our cohort to 128 GCA patients for protein quantification of tumors and NATs (**Supplementary Figure 1**). 31 patients showed HER2 negativity in both tumors and NATs, while 97 patients (75.78%) were HER2 positive. We calculated the HER2 level ratio between tumor and corresponding NATs to quantify HER2 protein levels across patients. Patients were then categorized into HER2-high (n = 48), HER2-low (n = 49), and HER2- negative (n = 31) groups based on this ratio (**Figure 1f**, **Methods**). The HER2-high group showed the best survival rates among the three groups (*p* = 0.014, **Figure 1g**).

**Figure 1:**
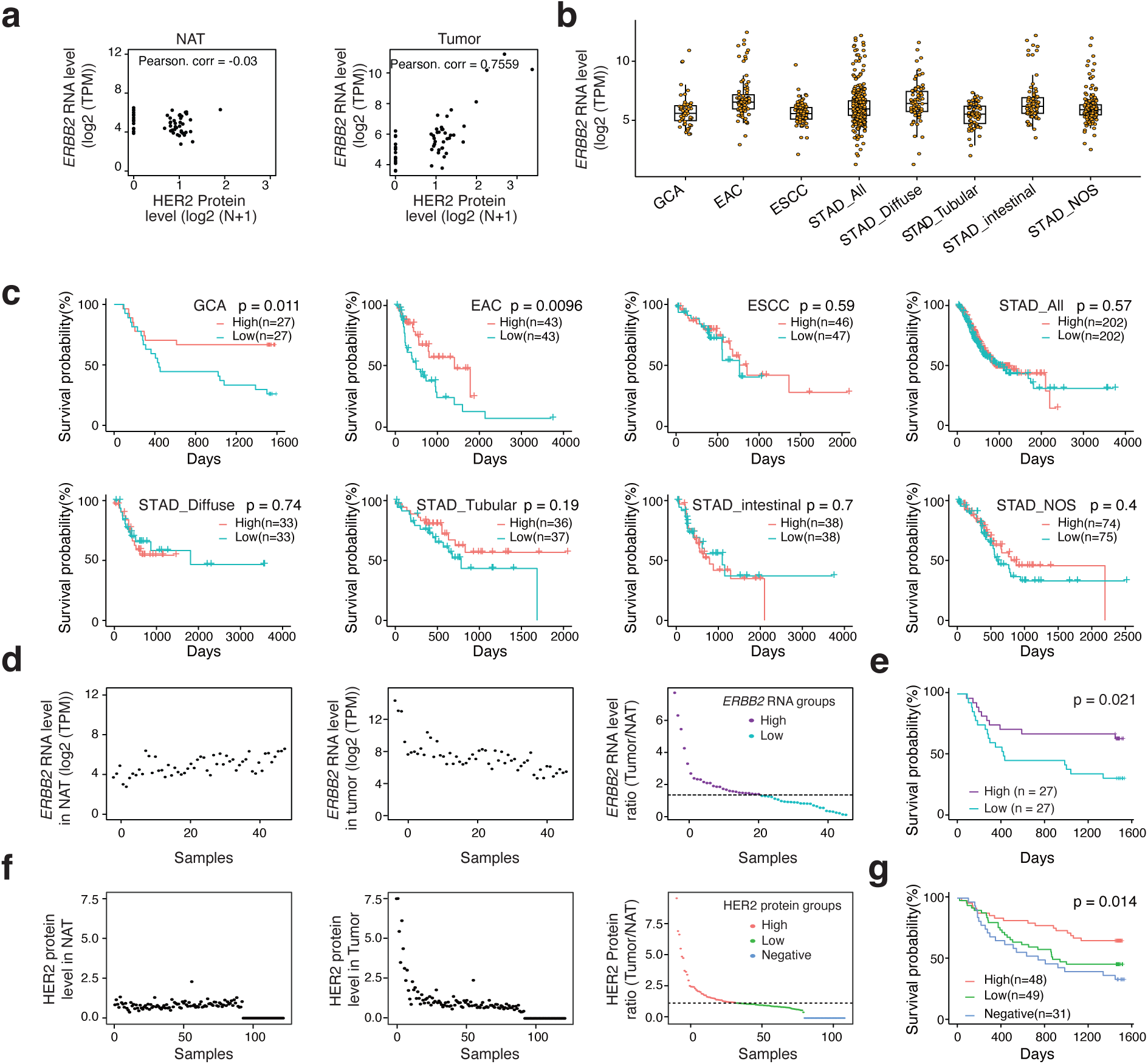
The expression of *ERBB2*/HER2 is a favorable prognosis marker for esophageal adenocarcinoma (EAC) and gastric cardia adenocarcinoma (GCA). **a.** The Pearson correlation between *ERBB2* RNA expression and HER2 protein expression in normal tissue adjacent to the tumor (NAT) (left panel) and tumor (right panel) from the GCA cohort. N = HER2 protein expression levels detected by mass spectrometry. **b.** The distribution of normalized *ERBB2* RNA expression in the GCA cohort, as well as in EAC and STAD from TCGA. ESCC = Esophageal squamous-cell carcinoma; STAD = Stomach adenocarcinoma; TPM = Transcripts Per Million. **c.** Kaplan-Meier survival curves of two patient groups based on the median *ERBB2* RNA expression from TCGA. High = upper half of patients; Low = lower half of patients. **d.** The strategy for dividing the GCA cohort into two groups based on the median ratio of *ERBB2* RNA expression levels between tumors and NATs (Left) and prognosis analysis for each group (Right). **e.** Kaplan-Meier survival curves of two patient groups based on ratio of ERBB2 RNA expression. **f.** The strategy for dividing the GCA cohort into three groups based on the ratio of HER2 protein expression levels between tumors and NATs. “Negative” refers to patients with HER2-negative expression; “High” refers to the upper half of HER2-positive patients; “Low” refers to the lower half of HER2-positive patients. **g.** Kaplan-Meier survival curves of three patient groups based on ratio of HER2 protein expression.

The grouping assignment of patients based on HER2 protein levels showed a significant association with the HER2 RNA-based grouping assignment (**Supplementary Figure 3a**). The HER2 scoring system was applied to the tumor samples in our GCA cohort (**Supplementary Figure 3b**), where 6.3% of patients were HER2 3+. The HER2-high group showed the largest proportion of HER2 3+ cases (14%), compared to 2% in the HER2-low group and none in the HER2-negative group. HER2 scoring significantly correlated with the MS HER2 grouping method (*p* = 0.034), suggesting that MS groups may accurately reflect HER2 expression (**Supplementary Figure 3c**). Post-surgery therapeutic strategies were unrelated to HER2 grouping (*p* = 0.320, **Supplementary Figure 3d**) or tumor stage (**Supplementary Figure 3e**) ansd did not affect prognosis of patients (*p* = 0.54, **Supplementary Figure 3f**). Correlation analysis showed that HER2 grouping was significantly associated with age (*p* = 0.005), family history (*p* = 0.049), alcohol consumption (*p* = 0.031), tumor differentiation (*p* = 0.029), and Lauren classification (*p* = 0.022) but was independent of other clinicopathological features (**Supplementary Figure 4a** and 4b**, Table S1**). We also analyzed the association between prognosis and Lauren classification and found that Lauren classification alone was not a prognostic marker (**Supplementary Figure 5**). However, the HER2-high group showed better prognosis across Lauren classifications, though this was not statistically significant. These findings suggest that while Lauren classification alone may not predict prognosis, it gains prognostic value when combined with HER2 grouping. In summary, integrating TCGA RNA-Seq data with our findings confirms that *ERBB2* RNA expression and HER2 protein levels are favorate prognostic markers for both GCA. This novel observation in GCA and EAC suggests that HER2-positive and high patient groups could be very different from those in other tumor types, such as breast cancer, where HER2-positive patients usually have a poor prognosis ^15^. Even though anti-HER2 therapies have started to be used for treatment in gastric and esophageal cancers^13,14,16^, following the strategy used in breast cancer, the specific features of HER2-positive patients in GCA may differ significantly from those in breast cancer. Detailed molecular characterization of GCA patients based on HER2 expression is crucial for better understanding GCA and identifying therapeutic targets for HER2-negative patients. Motivated by this, we conducted comprehensive analyses on our GCA cohort, including WES, RNA-Seq, scRNA-Seq, and spatial transcriptomics (**Figure 2a**).

**Figure 2:**
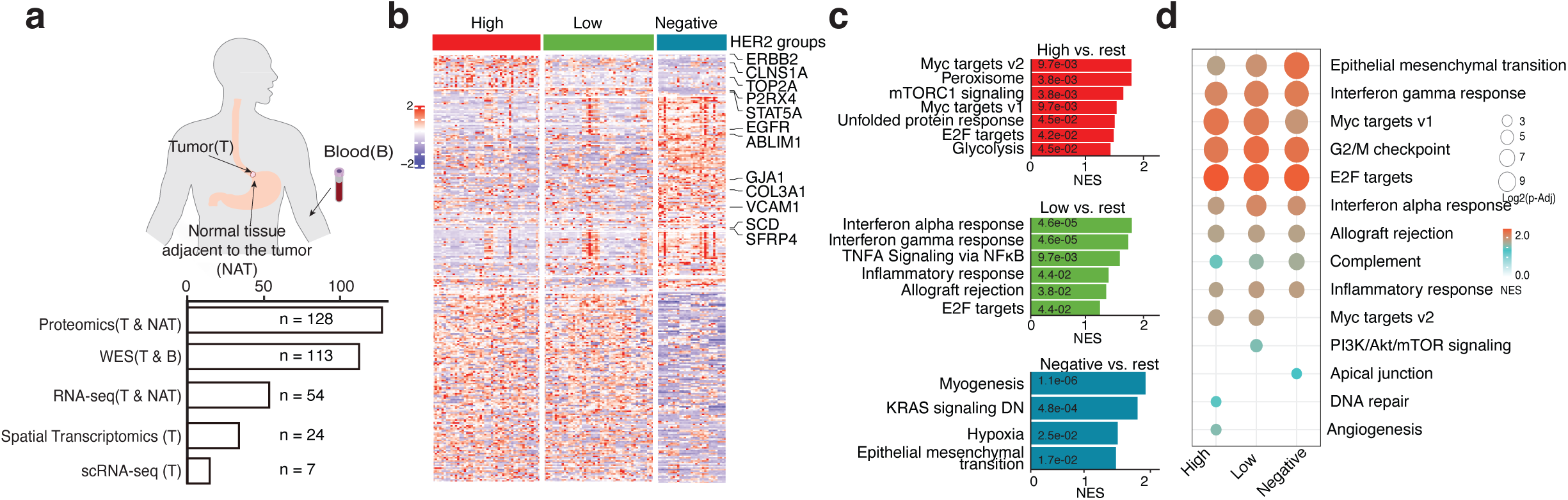
Protein expression landscape of different HER2 groups from GCA patients. **a.** The illustration of multiple omics data from GCA patients. **b.** Unique protein profiles of different HER2 GCA patient groups from tumor only. **c.** Gene set enrichment analysis (GSEA) with protein of different HER2 GCA patient groups from tumor only comparison. **d.** GSEA of different HER2 GCA patient groups by comparing tumors with corresponding normal tissue adjacent to the tumor (NAT).

### Enrichment of DNA repair pathways in HER2-High tumors and inflammatory signatures in HER2-Low/Negative tumors

A similar number of proteins were detected in both tumors (median = 5,088) and NATs (median = 5,089) (**Table S2**). To compare HER2 groups effectively, we used two strategies to identify signature proteins: comparing tumor protein expression across HER2 groups and evaluating protein levels by comparing tumors to NATs within each HER2 group. In the tumor-only comparison, we identified 23 proteins unique to the High group, 6 to the Low group, and 92 to the Negative group. Additionally, 31 proteins were shared by the Low and Negative groups, 11 by the High and Negative groups, and 132 by the High and Low groups (Fold Change > 1.5, *p* < 0.05) (**Supplementary Figure 6a**, **Figure 2b**, **Table S3**). The HER2- negative group had the most differentially expressed proteins. Gene set enrichment analysis (GSEA) of hallmark gene sets for each HER2 group revealed that the HER2-high group was enriched in proliferation-related sets, including Myc targets, mTORC1 signaling, and E2F targets (**Figure 2c**, **Supplementary Figure 6b**, **Table S4**). Similarly, E2F targets indicated proliferation features in the HER2-low group. Notably, the HER2-low group showed strong inflammatory features, with enrichment in Interferon alpha response, Interferon gamma response, and TNFA signaling via NFκB. Comparing tumors with their corresponding NATs (**Figure 2d**, **Table S5**) revealed that Myc targets v1 were more enriched in the HER2-high and low groups than in the HER2-negative group. Inflammatory hallmarks were present in all groups, with the interferon-alpha response notably enriched in the HER2-low group. Additionally, DNA repair and angiogenesis hallmarks were uniquely enriched in the HER2- high group. We analyzed differential protein expression between tumors and NATs for each HER2 group (Fold Change > 1.5, *p* < 0.05) (**Supplementary Figure 7a, 7b**, **Table S6**). In the HER2-high group, we identified 167 up-regulated and 299 down-regulated proteins; the HER2- low group had 99 up-regulated and 251 down-regulated proteins; and the HER2-negative group had 154 up-regulated and 305 down-regulated proteins. Gene ontology (GO) enrichment analysis of significant up-regulated proteins revealed distinct biological processes (**Supplementary Figure 7c**, **Table S7**). The HER2-high group showed enrichment in DNA replication terms, while the HER2-low group enriched immune response terms, and the HER2- negative group enriched terms related to viral response.

We extracted common and unique proteins among the three HER2 groups (**Supplementary Figure 8**). The HER2-high group had 60 specific proteins, including HER2, MCM2, MCM3, and MCM7, highlighting HER2’s role in cell cycle regulation and DNA replication. In contrast, the HER2-low group contained 15 specific proteins enriched in inflammatory-related proteins like TAP1, TAP2, IFT1, and IFIT3 ^25^, consistent with the inflammation features observed in GSEA and GO term enrichment analyses. The HER2-negative group included 47 specific proteins with HLA-DRA and HLA-A, crucial for immune response ^26^. Overall, our analysis indicated that HER2-high groups exhibited stronger proliferation and DNA repair features, while HER2-negative and low groups displayed more pronounced inflammatory features.

### Comparable mutation burden across groups with elevated genome gains in HER2-High patients

Next, we examined genome mutations and gains among the three HER2 groups (**Figure 3**, **Supplementary Figure 9**, **Table S8**). Although there was a slight reduction in SNV and InDel frequencies in the HER2-negative group, statistical analysis revealed no significant differences (**Supplementary Figure 10a**, **10b**). All three groups showed a high degree of similarity in mutation types (**Supplementary Figure 10c**). Additionally, there are no significant differences in non-synonymous or SNV frequencies across the groups (**Supplementary Figure 10d**). We compared the mutation frequencies of the top 20 mutated cancer genes across each group (**Figure 3a**, **Supplementary Figure 10e**, **Table S9**). Notably, ARID1A was the only gene with a significantly different mutation frequency among the three groups (*p* = 0.016), with higher mutations in the HER2-negative and HER2-low groups. Patients with ARID1A mutations had a worse prognosis (*p* = 0.00069) (**Figure 3b**). While prognosis varied among the three HER2 groups, those with ARID1A mutations generally fared worse, with significant differences in the HER2-high (*p* < 0.0001) and HER2-low groups (*p* = 0.036), but not in the HER2-negative group (*p* = 0.35) (**Figure 3b**). These findings suggest that HER2 expression and ARID1A mutation signatures together impact the prognosis of GCA patients. This conclusion is supported by examining patient prognosis differences across the three HER2 groups in both mutated and unmutated contexts (**Figure 3c**). Other top mutated genes, such as FAT4, may also affect patient prognosis in conjunction with HER2 grouping (**Supplementary Figure 11**). We assessed chromosomal instability (CIN) in the three groups using the genome integrity index (GII) ^27^ and weighted genome instability index (wGII) ^28^. The mean GII values were 0.102 for the HER2-high group, 0.078 for the HER2-low group, and 0.071 for the HER2- negative group. The HER2-high group’s GII was 1.44 times higher than that of the HER2- negative group (*p* = 0.015) (**Figure 3d**). Similarly, the mean wGII values were 0.060 for the HER2-high group, 0.027 for the HER2-low group, and 0.021 for the HER2-negative group, with the HER2-high group showing a 2.86-fold increase over the HER2-negative group (*p* = 0.0096) (**Figure 3d**).We also calculated chromosomal breakpoints, represented as the CIN Index (CINindex) and weighted CIN Index (wCINindex). The HER2-high group exhibited significantly higher CINindex values, 1.36 times greater than the HER2-negative group (*p* = 0.011), and wCINindex values, 1.37 times higher (*p* = 0.0064) (**Figure 3e**). While the HER2- high group showed more copy number alterations (CNA), the difference in CNA across the three HER2 groups was not statistically significant (*p* = 0.064) (**Figure 3f**).

**Figure 3:**
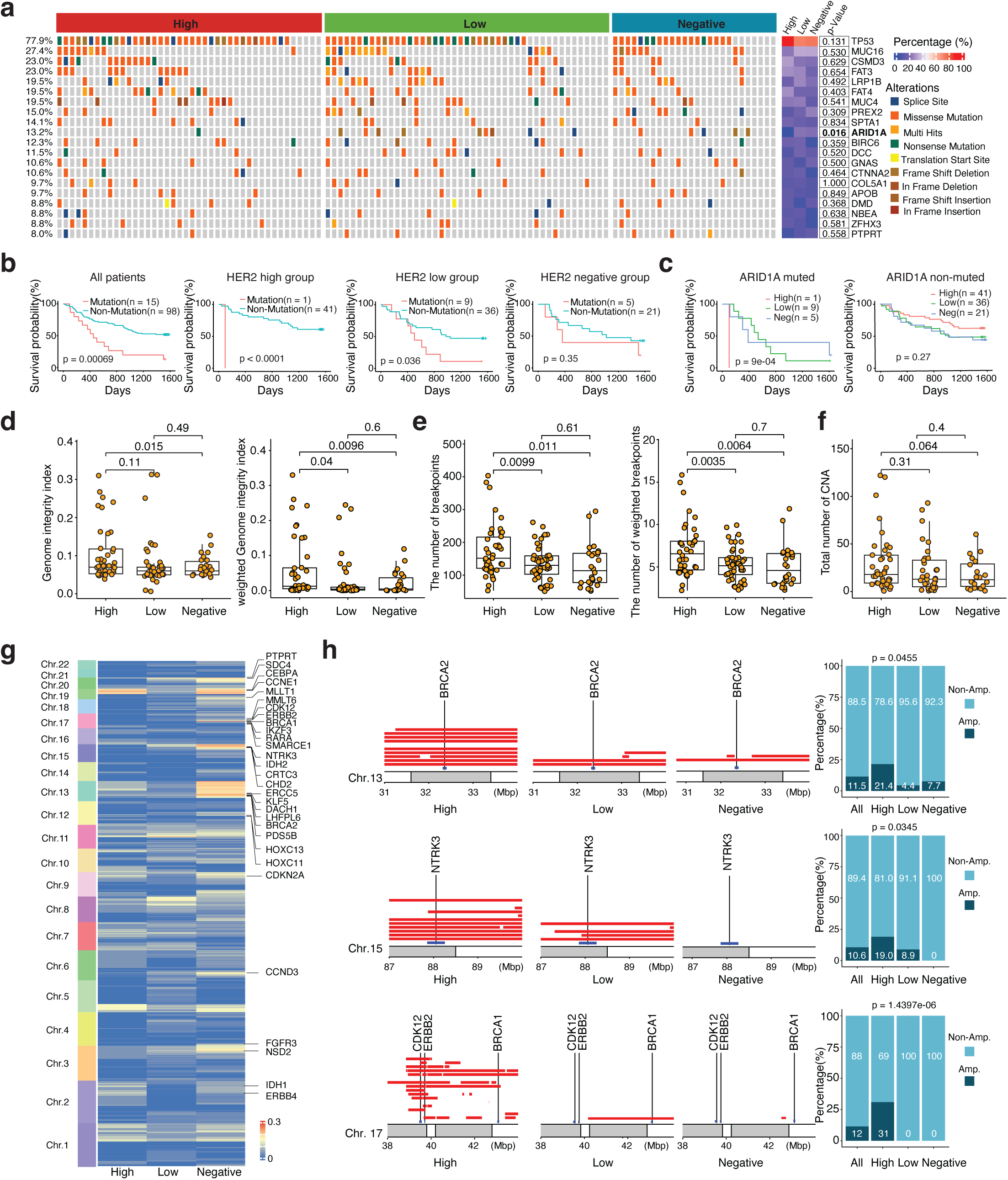
Somatic mutation and gain landscape of different HER2 groups from GCA patients. **a.** The top 20 mutated oncogenes and tumor suppressor genes captured in the GCA cohort, along with their mutation frequency in different HER2 groups. **b.** Comparison of prognosis in GCA patients with and without *ARID1A* gene mutations. **c.** Comparison of prognosis in GCA patients among the three HER2 groups and *ARID1A* gene mutations **d.** Comparison of chromosome instability (CIN) among the three different HER2 groups using the genome integrity index. **e.** Comparison of the number of breakpoints among the three different HER2 groups. **f.** Comparison of copy number alterations (CNAs) among the three different HER2 groups. **g.** The significantly different CNA regions in the three HER2 groups with a bin size of 1 Mbp; oncogenes and tumor suppressor genes are labeled on the side. **h.** Examples of significant CNA regions in the three HER2 groups on chromosomes 13, 15, and 17.

We calculated genome gain frequencies using 1 Mbp bins among the HER2 groups (**Figure 3g**, **Supplementary Figure 12**, **Table S10**). Notably, 142 regions exhibited higher gains in the HER2-high group, while 16 regions were more amplified in the HER2-negative group compared to both the high and low groups. Subregions of Chromosomes 13, 15, and 17 showed increased gains in the HER2-high group (**Figure 3g**, **3h**). Interestingly, Chromosome 19 subregions displayed high gains in both the HER2-high and negative groups but not in the low group. We also examined oncogenes and tumor suppressor genes (TSGs) in the gained regions (**Figure 3g**, **3h**). *CCNE1*, *MLLT1*, *CCND3*, and *FGFR3* were significantly gained in the HER2- high group. Notably, a subregion of Chromosome 13 containing the TSG *BRCA2* showed greater gain in the HER2-high group compared to the others. Additionally, ERBB2 gene gains were markedly more frequent in the HER2-high group (31%) compared to the low and negative groups (0%). Another TSG, *BRCA1*, located near ERBB2, also showed increased gain in the HER2-high group, though not statistically significant. In summary, our findings indicate that mutation is not a main distinguishing factor among these groups; however, the HER2-high group shows a significantly higher frequency of genome gains.

### Stronger inflammation features are observed in HER2-negative groups

Clear distinctions were observed between the RNA expression profiles of tumors and NATs (**Supplementary Figure 13a**). GSEA of tumor RNA-Seq data revealed significant enrichment in six cancer hallmark terms in the HER2-high group (**Figure 4a**, **Table S11**), particularly in DNA repair features. The HER2-low group enriched 20 cancer hallmarks, with inflammation-related terms like TNFA signaling via NFκB and Interferon gamma response. The HER2- negative group mainly exhibited enrichment in inflammatory response and allograft rejection. A differential gene expression analysis identified 572 significantly differentially expressed genes (Fold Change > 1.5, FDR < 0.05) across the HER2 groups (**Figure 4b**, **Supplementary Figure 13b**, **Table S12**). DNA repair-related genes, such as H2AFX ^29^, were enriched in both HER2-high and low groups (**Figure 4b**), while infection and immune response genes like *TNFSF4*, *TNFSF18*, and *TRIM50* were uniquely enriched in the HER2-negative group. We proceeded with GSEA analysis within each HER2 group comparing tumor samples with their corresponding NATs (**Figure 4c**, **Table S13**). Distinct enrichment patterns were evident, with unique terms like TNFA signaling via NFκB, Apical junction, IL-6/JAK2/STAT3 signaling, and Apoptosis enriched in HER2-low and negative groups. Notably, DNA repair hallmark was more pronounced in HER2-high and low groups. Subsequently, a differential gene expression analysis comparing tumors with NATs (Fold Change > 1.5, FDR < 0.05) revealed 5084, 5338, and 4697 differentially expressed genes in the HER2-high, -low, and -negative groups, respectively (**Supplementary Figure 14Aa-c**, **Table S14, Table S15**).

**Figure 4:**
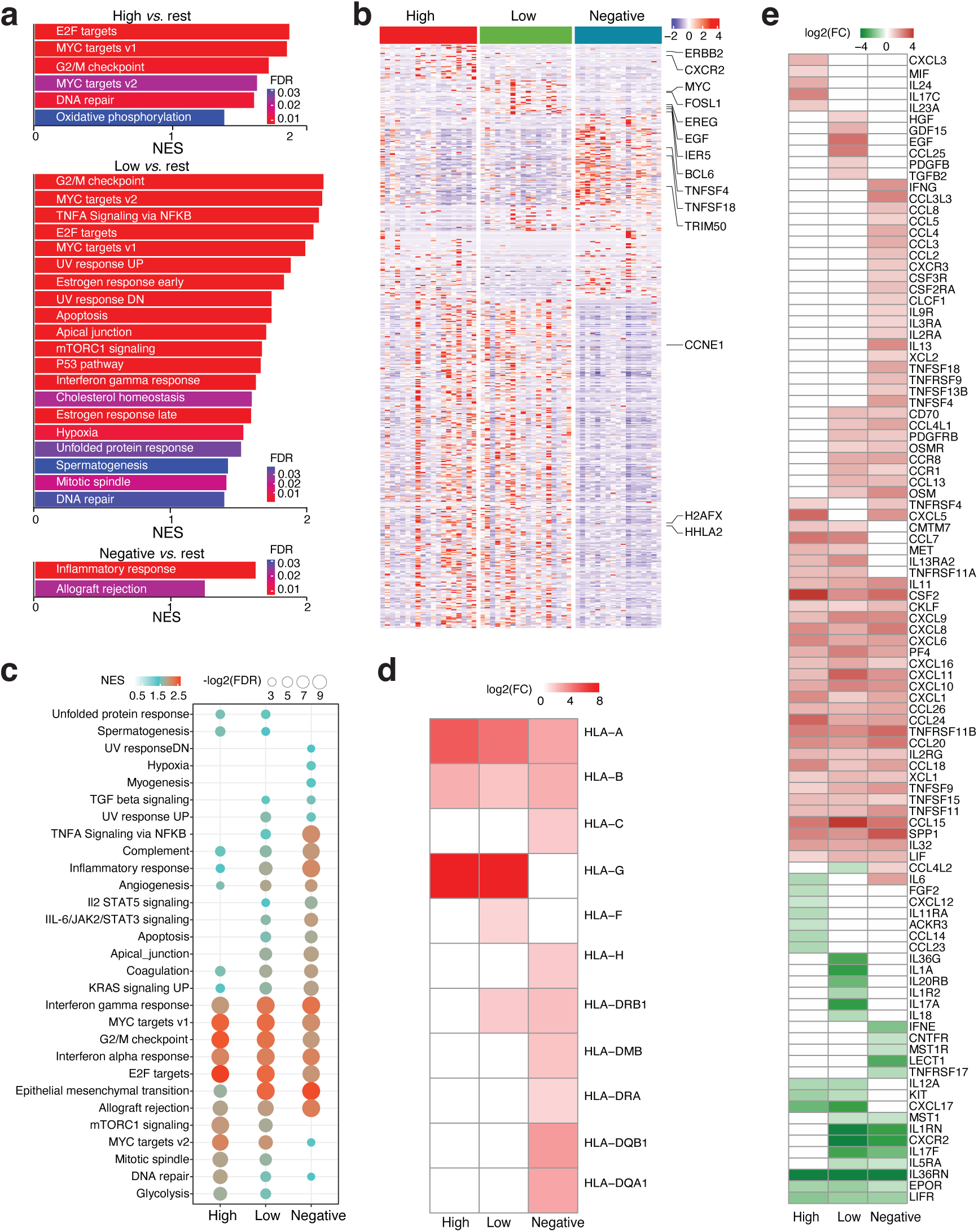
Gene expression signature of different HER2 groups from GCA patients. **a.** Gene set enrichment analysis (GSEA) using RNA expression from tumors of different HER2 GCA patient groups. **b.** Gene expression signatures from tumors of different HER2 GCA patient groups. **c.** GSEA with RNA expression of different HER2 GCA patient groups by comparing tumors with corresponding normal tissue adjacent to the tumor (NAT). **d.** Differential RNA expression from genes of major histocompatibility complex (MHC) class I and II in different HER2 groups by comparing tumors with corresponding NATs. **e.** Differential RNA expression from genes of cytokines from different HER2 groups by comparing tumors with corresponding NATs. **f.** FDR = false discovery rate.

Notably, MHC class II genes, including HLA-DMA, HLA-DMB, HLA-DRA, HLA-DQB1, and HLA-DQA1, were active only in the HER2-low and -negative groups (**Figure 4d**). GO term analysis using the list of up-regulated genes in tumors compared to NATs showed that cell cycle and DNA repair-related GO terms were strongly enriched in both the HER2-high and HER2-low groups, while inflammatory terms were prominently enriched in the HER2- negative group (**Supplementary Figure 15a-c, Table S16**). We also identified exclusively up- regulated genes in each HER2 group for these targeted GO terms (**Table S17**). Notably, genes like *H2AFX* and *BRCA2*, crucial for DNA repair, were significantly expressed in HER2-high tumors compared to NATs. *BRCA1* expression was significantly higher in HER2-high and low tumors compared to NATs. Additionally, MHC class II GO terms were exclusively enriched in the negative group (**Supplementary Figure 15d**). In the HER2-high group, all the top 15 GO terms are associated with the mitotic cycle and chromosome segregation (**Supplementary Figure 15e**). Similarly, in the HER2-low group, 9 of the top 15 GO terms focus on these processes, with an additional layer related to inflammation. In the HER2-negative group, 10 of the top 15 GO terms primarily revolve around inflammation. We analyzed the differential expression of cytokine genes between tumor and adjacent tissue in each HER2 group (**Figure 4e**). The HER2-negative group showed the highest activity, with 33 cytokine genes, including IL-6 and IFNG, exclusively active in HER2-negative tumors. IL-6 is linked to chronic inflammation ^30^, while IFN-γ promotes inflammation ^31^, underscoring distinct inflammatory responses in this group. We also evaluated 18 tumor inflammation genes ^32^ (**Supplementary Figure 16**); 50% were significantly elevated in HER2-negative tumors, 44.4% in HER2-low tumors, and only 16.6% in HER2-high tumors. These findings indicate distinct inflammatory tumor microenvironments across HER2 groups.

### Single-cell RNA-Seq reveals that lower HER2 expression is linked to stronger inflammatory features and higher M2 macrophage levels

To better understand the tumor microenvironment in different HER2 groups, we first used CIBERSORTx ^33^ to analyze immune cells in HER2 groups from bulk RNA-Seq data. We identified 22 immune cell types, with no significant differences in NATs (**Supplementary Figure 17a**, **17b**). In tumors, the HER2-negative group had significantly more M2 macrophages than the HER2-high group (*p* = 0.036), while the HER2-low group showed a higher but not statistically significant proportion (*p* = 0.21) (**Figure 5a**, **5b**, **Supplementary Figure 17c**). Staining for the M2 macrophage marker CD163 confirmed a higher presence in the HER2-low (*p* = 0.016) and negative groups (*p* = 0.18) compared to the HER2-high group (**Figure 5c**, **5d**, **Table S18**). Although the difference between the HER2-high and negative groups was not significant, there was a trend toward more CD163-positive cells in the HER2- negative. Our findings suggest distinct tumor microenvironments linked to HER2 expression levels in GCA patients.

**Figure 5:**
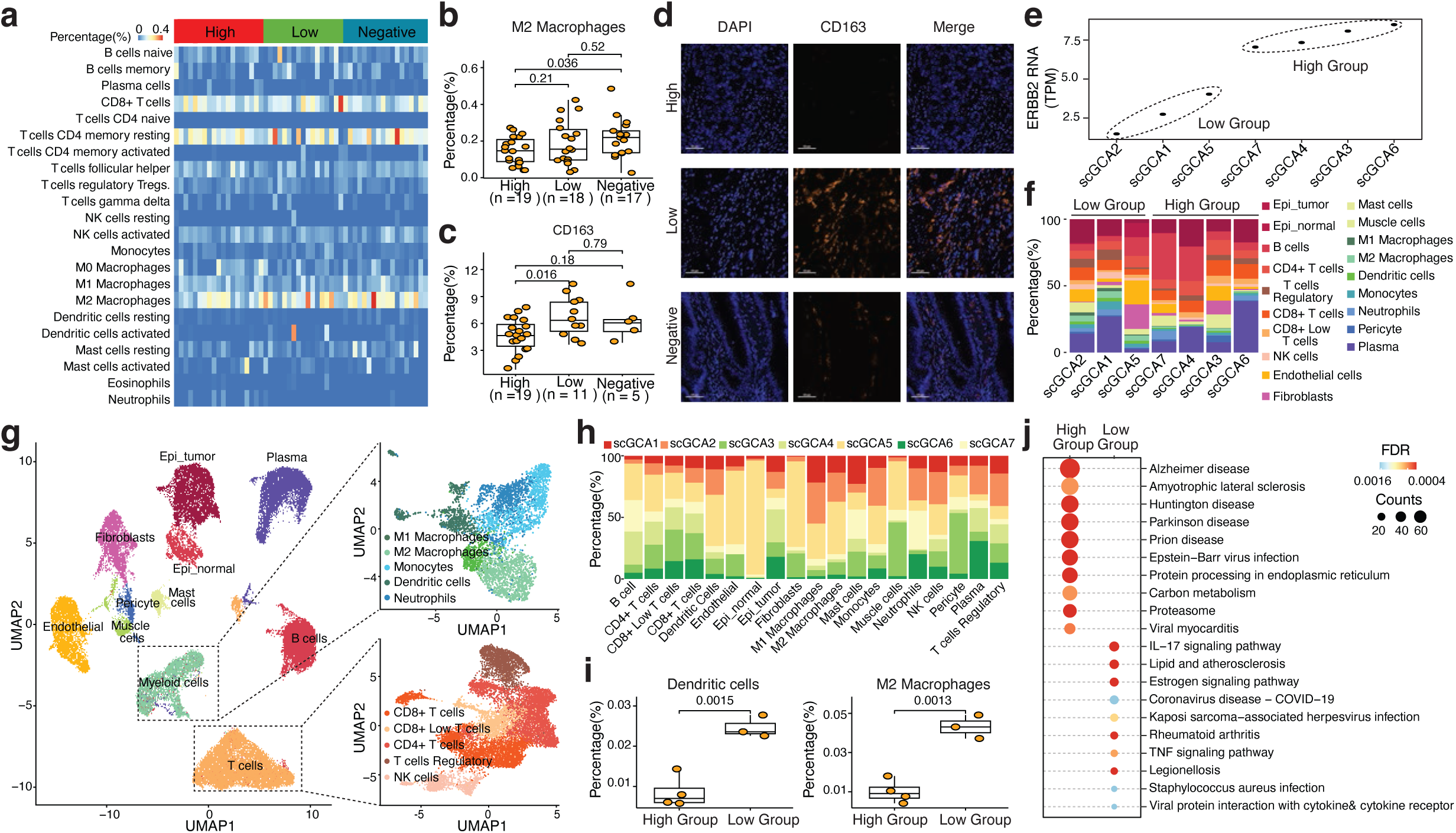
Deciphering tumor microenvironment of different HER2 groups from GCA patients. **a.** Comparison of cell components from tumor bulk RNA-seq predictions across different HER2 groups. **b.** Quantitative comparison of M2 macrophages in tumors only from different HER2 groups. **c.** Quantification of immunohistochemistry (IHC) staining of CD163 in tumors across different HER2 groups. **d.** Representative images of immunohistochemistry (IHC) staining of CD163 in tumors from different HER2 groups. **e.** Grouping of GCA patients into *ERBB2* high and low samples based on *ERBB2* gene expression levels. **f.** Proportional quantification of different cell types in seven GCA patients from scRNA-Seq. **g.** Cell type identification in seven GCA patients from scRNA-Seq. **h.** Distribution of different cell types in seven GCA patients. **i.** Quantitative comparison of dendritic cells (left panel) and M2 macrophages (right panel) in different HER2 groups from scRNA-Seq. **j.** The top 10 pathways enriched from KEGG (Kyoto Encyclopedia of Genes and Genomes) analysis in different HER2 groups from M2 macrophages.

To further confirm our findings, we performed scRNA-Seq on tumors from seven GCA patients, categorizing them into HER2 high (n = 4) and HER2 low groups (n = 3) based on HER2 RNA expression (**Supplementary Figure 18a-d**, **Figure 5e**). The analysis identified 46,912 cells, revealing 19 distinct cell types heterogeneously distributed among the patients (**Figure 5e-h**, **Supplementary Figure 18e**, **Figure S19**). Significant differences between the HER2 high and low groups were found only in M2 macrophages (*p* = 0.0013) and dendritic cells (*p* = 0.0015) (**Figure 5i**, **Supplementary Figure 20**), confirming higher M2 macrophage levels in lower HER2 expression patients. Differential gene expression analysis of M2 macrophages identified 1,591 highly expressed genes for the HER2-high group and 86 genes for the low group. Cytokine and chemokine genes like CXCL3 and S100A were specifically enriched in the low group (**Supplementary Figure 21**). KEGG analysis showed stronger inflammatory features, including TNF signaling, in the HER2-low group (**Figure 5j**, **Table S20**), which is further supported by GO term enrichment (**Supplementary Figure 21b**, **21c**, **Table S21**). Overall, lower HER2 expression is linked to stronger inflammatory features and higher M2 macrophage levels.

### Single cell analysis uncovered that low HER2 expression correlates with stronger inflammatory signaling pathways in epithelial tumor cells

Next, we focused on epithelial tumor cells from scRNA-Seq to understand the communication between tumor cells and their microenvironment. We identified seven subclusters of epithelial tumor cells and found no significant differences in their distribution in the two groups (**Figure 6a**). We defined cells with high CNV and aneuploidy in epithelial cells (**Figure 6a**, **6b**; **Supplementary Figure 22**). As expected, tumor cells exhibited significantly higher CNV scores than normal epithelial cells (**Figure 6b**, *p* < 2.2e-16). We found no significant differences in the distribution of aneuploid epithelial cells and high CNV cell numbers between groups (**Figure 6a**, **6b**). We performed pseudotime trajectory analysis of epithelial tumor cells, identifying three cell states (**Figure 6c**, **6d**, **Supplementary Figure 23**). Epithelial tumor cells in the HER2 high group were more enriched in state 3, while those in the low group were more prevalent in state 2 (*p* < 2.2e-16). Based on this analysis, we defined two cell fates (**Figure 6c**) and extracted gene features for each fate (**Figure 6e**, **Table S24**). Cell fate 1, enriched in the low group, exhibited high expression of cytokines and chemokines (e.g., *CCL11*, *CXCL5*) (**Figure 6e**). Cell fate 2, enriched in the high group, showed elevated expression of growth factors (e.g., *ERBB3*, *NRG1*). KEGG analysis revealed stronger inflammatory features in the low group, with pathways like IL-17 signaling, while the high group showed enrichment in growth factor pathways, including ERBB signaling (**Figure 6f**, **Table S23**).Our data suggest that low HER2 expression correlates with stronger inflammatory features in the epithelial tumor cells of GCA patients.

**Figure 6:**
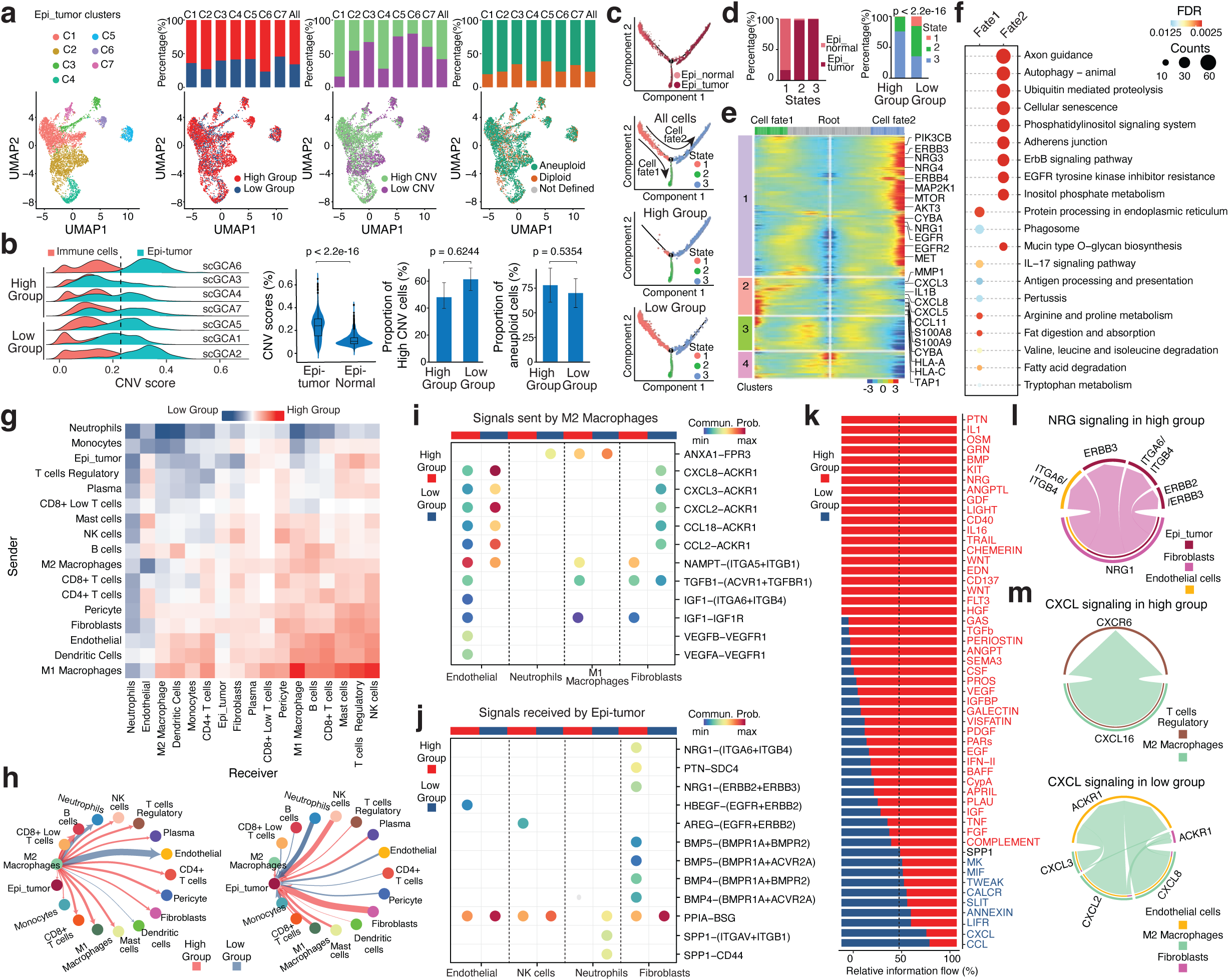
Unique signal pathways in epithelial cells of different HER2 groups from GCA patients. **a.** Cell type identification in epithelial tumor cells (first left), the distribution of cells in high and low groups (second left), the distribution of high and low CNV (copy number variants) in each cell cluster (second right), and the distribution of aneuploid and diploid cells in each cell cluster (first right). **b.** Identification of high CNV (copy number variants) cells (first left), comparison of CNV values between epithelial tumor cells and epithelial normal cells (second left), comparison of the proportion of high CNV cells in high and low groups (second right), and comparison of the proportion of aneuploid cells in high and low groups (first right). **c.** Pseudotime projection of epithelial tumor cells and the distribution of cells from high and low groups. **d.** Distribution of epithelial normal cells and tumor cells in different cell states (left) and different cell states in high and low groups (right). **e.** Extracted feature genes for different cell fates from epithelial tumor cells. **f.** The top 10 pathways enriched from KEGG (Kyoto Encyclopedia of Genes and Genomes) analysis for different cell fates from epithelial tumor cells. **g.** Comparison of signal communication among different cell types in high and low groups. **h.** Circle plots presenting signal communication sent by M2 macrophages (left) and received by epithelial cells (right). **i.** Representative cell signals sent by M2 macrophages from scRNA-Seq. **j.** Representative cell signals received by epithelial tumor cells from scRNA-Seq. **k.** Signal strength comparison from different signaling pathways in high and low GCA groups from scRNA-Seq. **l.** Circle plots presenting signal pathway communication of NRG in high group. **m.** Circle plots presenting CXCL pathway contributed by M2 macrophages in high and low groups.

We observed stronger signaling communications in the HER2 high group compared to the low group (**Figure 6g, Supplementary Figure 24a**, **24b**). In the high group, M1 macrophages exhibited stronger communication with other cell types, while CD4+ and CD8+ T cells interacted well with most. In the low group, M2 macrophages communicated with neutrophils, endothelial cells, and other M2 macrophages, whereas in the high group, they interacted with CD4+ T cells, M1 macrophages, and others (**Figure 6g**, **6h**, **Supplementary Figure 25**). M2 macrophages in the low group sent stronger cytokine and chemokine signals, while in the high group, they sent stronger growth factor signals (**Figure 6i**, **Supplementary Figure 24c**). Epithelial tumor cells received stronger growth factor signals in the high group and stronger inflammatory signals in the low group (**Figure 6j**, **Supplementary Figure 25d**). Overall, the high group exhibited stronger growth factor communication, whereas the low group displayed stronger inflammatory signaling (**Figure 6k**, **Supplementary Figure 26**). NRG signaling was exclusively contributed by epithelial tumor cells, endothelial cells, and fibroblasts in the high group (**Figure 6l**). Although CCL and CXCL signaling was stronger in the low group, communication was also present in the high group (**Supplementary Figure 27**). In the high group, CXCL signaling was driven by interactions between regulatory T cells and M2 macrophages, while in the low group, it involved M2 macrophages, endothelial cells, and fibroblasts (**Figure 6m**). Together, our data suggest distinct tumor microenvironments, with lower HER2 expression linked to stronger inflammation, based on HER2 expression levels in GCA patients. This conclusion is consistent with our findings from bulk RNA and protein analyses.

### The CD47-SIRPA axis as a promising therapeutic option for GCA patients with low HER2 expression

Since there is a distinct tumor microenvironment in different HER2 groups, with stronger inflammation in HER2-negative groups, we were motivated to identify potential therapeutic targets for HER2-negative patients. Immunotherapy has become a promising therapeutic treatment for esophageal and gastric cancers, often combined with anti-HER2 treatments. We focused on identifying specific immune checkpoints for different HER2 groups. To identify distinct immune inhibitory checkpoints among the three HER2 groups, we compared the RNA expression levels of key immune checkpoint genes, ligands, and receptors between tumors and adjacent normal tissue (NAT) in each group (**Figure 7a**). The HER2-negative group showed significantly higher levels of FGL1 and TNFSF18 in tumors compared to NAT. Both HER2- high and HER2-low groups had elevated HHLA2 expression, while the HER2-low group exhibited increased PVR and CD276 levels. Additionally, CD70, PDCD1LG2 (PD-L2), and SIRPA were higher in HER2-low and HER2-negative tumors. PD-L2 is a ligand for PD-1 and a key immunotherapy target, while the CD47/SIRPA axis is also emerging as a potential therapeutic target ^34^. We further investigated the relationship between HER2 expression and immune checkpoint levels, focusing on PD-L2 and CD47/SIRPA in different GCA patient groups. We performed staining of PD-L2 and SIRPA in tumors from different HER2 groups of GCA patients (**Figure 7b**). The proportion of PD-L2-positive cells was significantly higher in the HER2-negative group than in the HER2-positive group (*p* = 0.033), and SIRPA-positive cells were also more prevalent in HER2-negative patients (*p* = 0.039) (**Figure 7c**, **Table S24**). These findings indicate elevated PD-L2 and CD47/SIRPA axis expression in the HER2- negative group.

**Figure 7:**
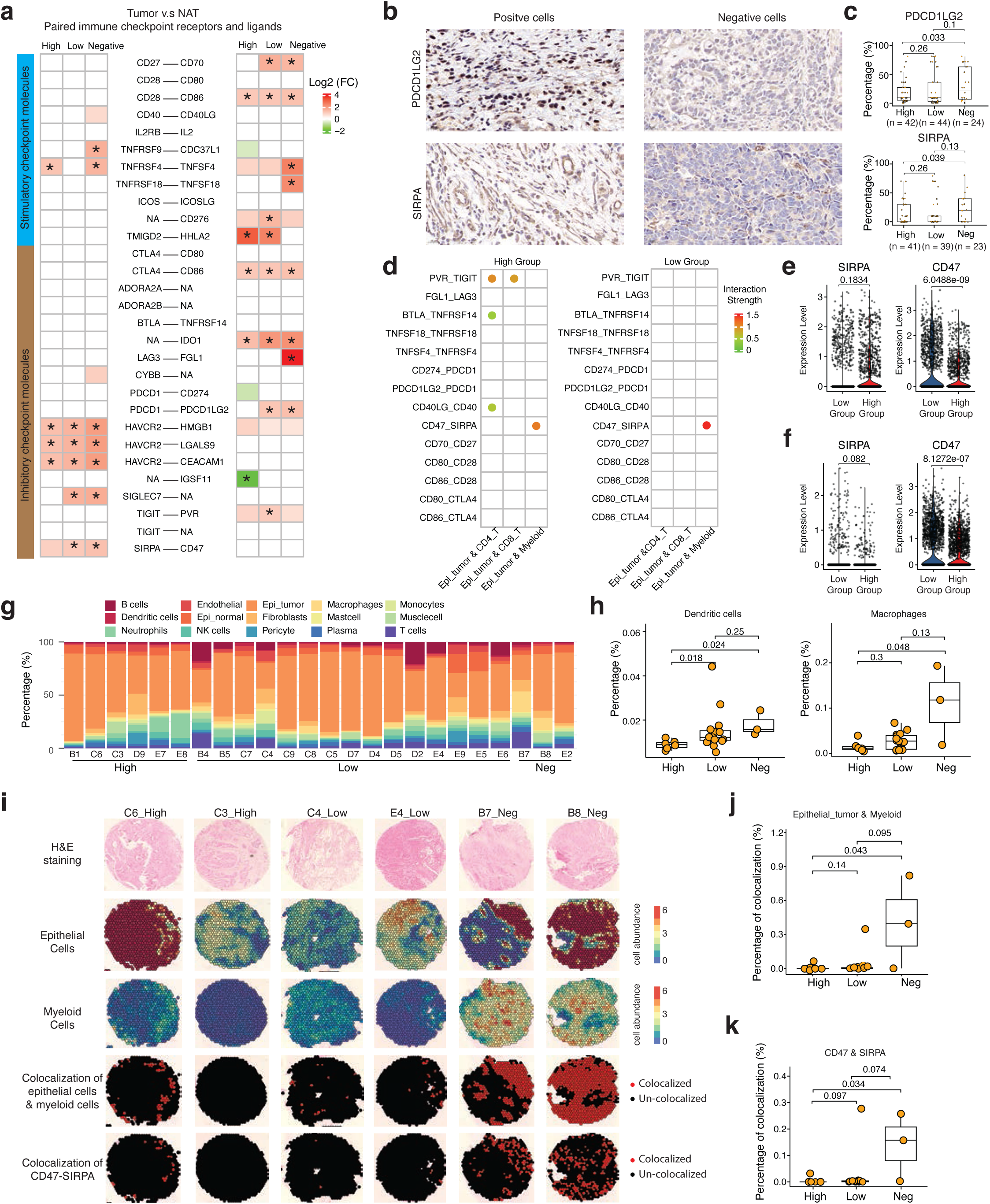
Spatial transcriptons uncovers distinct immune inhibitory checkpoints among three HER2 groups. **a.** Comparison of differential gene expression between tumors and corresponding normal tissue adjacent to the tumor (NAT) for paired immune checkpoint receptors and ligands in different HER2 groups; FC = Fold Change; white block = False Discovery Rate (FDR) greater than 0.05; an asterisk indicates |log₂(Fold Change)| ≥ 1 and FDR less than 0.05. **b.** Representative immunohistochemistry (IHC) staining of positive and negative cells for PD- L2 (PDCD1LG2) and SIRPA. **c.** Quantitative comparison of PD-L2 and SIRPA positive cells detected in the three HER2 groups. **d.** Interaction strength of paired immune checkpoint receptors and ligands in high and low groups detected in scRNA-Seq. **e.** Comparisons of *SIRPA* and *CD47* gene expression from M2 macrophages in high and low groups. **f.** Comparisons of *SIRPA* and *CD47* gene expression from epithelial tumor cells in high and low groups. **g.** Distribution of different cell types identified from spatial transcriptomics in the three GCA groups. **h.** Quantitative comparison of dendritic cells and macrophages in different HER2 groups from spatial transcriptomics. **i.** Representative images of H&E (Hematoxylin and Eosin) staining (top row), identified epithelial tumor cells (second row), myeloid cells (third row), co-localization of epithelial cells and myeloid cells (bottom second row), and co-localization of CD47-SIRPA in epithelial and myeloid cells (bottom row) from spatial transcriptomics in the three different HER2 groups. **j.** Quantitative comparison of the co-localization of epithelial cells and myeloid cells from spatial transcriptomics in the three different HER2 groups. **k.** Quantitative comparison of the co-localization of CD47-SIRPA in epithelial and myeloid cells from spatial transcriptomics in the three different HER2 groups.

Due to the lack of GCA cell lines and similarities between esophageal adenocarcinoma (EAC) and GCA, we employed the EAC cell line (OE19) to assess the relationship between ERBB2 expression and SIRPA and PD-L2 expression. Because CD47 and PD-L2 are not expressed in this cell line, we compared SIRPA gene expression levels in OE19 with and without treatment with the HER2 inhibitor lapatinib and after HER2 silencing via siRNA ^35^. The results showed a significant increase in SIRPA expression after HER2 inhibitor treatment and HER2 silencing (**Supplementary Figure 26**), confirming that decreased ERBB2 expression correlates with upregulated SIRPA.

We compared the strength of immune checkpoint interactions in different HER2 groups using scRNA-Seq data (**Figure 7d**). Although we did not observe an enriched interaction between PD-L2 and PD-1 due to low PD-L2 expression, we found a much stronger CD47-SIRPA interaction in the low group compared to the high group, specifically between epithelial and myeloid cells (**Figure 7d**, **Tables 25** and **26**). SIRPA expression was higher in the HER2-low group than in the high group among epithelial cells and M2 macrophages, though this difference was not statistically significant (**Figures 7e**, **7f**). Additionally, CD47 expression levels were significantly elevated in the low group compared to the high group in both epithelial cells (*p* = 6.0488e-09) and M2 macrophages (*p* = 8.1272e-07) (**Figures 7e**, **7f**). We also observed stronger immune checkpoint interactions, such as BTLA and TNFRSF14 between epithelial tumor cells and CD4+ T cells (**Figure 7d**). These findings indicate distinct immune checkpoints across HER2 groups in GCA patients, particularly in pathways involving SIRPA and CD47.

Next, we selected tumor-immune border regions from GCA patient samples to create tissue microarrays (**Supplementary Figure 29a**) and conducted spatial transcriptomics on these regions. High-quality data were obtained from 24 samples: 6 from the HER2-high group, 15 from the HER2-low group, and 3 from the HER2-negative group (**Supplementary Figure 29b- f**). We identified 15 cell types from the spatial transcriptomics data (**Figure 7g**, **Supplementary Figure 30**). Notably, the HER2-negative group had significantly higher levels of dendritic cells (*p* = 0.024) and macrophages (*p* = 0.048) compared to the HER2-high group (**Figure 7h**). Additionally, T cell levels were significantly higher in the HER2-low group compared to the HER2-high group (*p* = 0.014). However, no significant differences were observed in other cell types among the groups (**Supplementary Figure 31**). Since CD47 and SIRPA primarily interact between myeloid and tumor cells, we calculated the colocalization frequency of epithelial tumor cells and myeloid cells. This analysis revealed significantly higher colocalization in the HER2-negative group compared to the HER2-high group (*p* = 0.043) (**Figure 7i**, **7j**, **Supplementary Figure 32a**). We further conducted a molecular colocalization analysis of CD47-SIRPA at the interface of epithelial tumor and myeloid cells, revealing significantly higher colocalization in the HER2-negative group compared to the HER2-high group (*p* = 0.034) (**Figure 7i**, **7k**, **Supplementary Figure 32b**). Our findings strongly suggest that blocking the CD47/SIRPA axis could be a promising therapeutic option for GCA patients with low HER2 expression.

## Discussion

GCA exhibits distinct features compared to non-cardia gastric adenocarcinoma, yet specific therapeutic targets are lacking. To revela the gene signature of GCA patients and their therapeutic targets, we employed a highly sensitive molecular counting method using mass spectrometry to assess HER2 expression in 128 GCA patients. This assessment categorized patients into HER2-high, HER2-low, and HER2-negative groups, yielding novel insights into GCA and identifying new therapeutic targets, particularly within the HER2-negative subgroup. We discovered that the ratio of HER2 expression between tumors and their corresponding normal adjacent tissues (NAT) at both the RNA and protein levels serves as a valuable prognostic marker for GCA patients. We also found that the RNA expression of HER2 is a favorable prognostic marker for EAC, but not for gastric cancer. Although no significant differences in mutation burdens were observed among the three HER2 groups, we identified heightened genome instability in the HER2-high group. Notably, DNA repair pathways were significantly enriched in this group, likely due to the amplification and overexpression of tumor suppressor genes such as BRCA1 and BRCA2, which may contribute to the improved prognosis in this subgroup. Strong proliferation features in the HER2-high group could serve as potential therapeutic targets. Conversely, the HER2-low and HER2-negative groups exhibited significantly stronger inflammatory features. This suggests that anti-inflammatory drugs, in combination with other treatments, could be considered as part of the therapeutic regimen for these patients. Since anti-inflammatory cytokines can enhance anti-tumor immunity by modulating the inflammatory response ^36^, careful therapeutic design is essential. Additionally, we found that HER2 and ARID1A mutations together influence the prognosis of GCA patients. As the second most frequently mutated tumor suppressor gene ^37^, ARID1A presents therapeutic opportunities, particularly for HER2-low and HER2-negative patients.

Single-cell RNA-Seq and spatial transcriptomics revealed distinct tumor microenvironments and immune signaling pathways: the HER2-high group had stronger growth factor signaling, while the HER2-low and HER2-negative groups exhibited pronounced inflammation and higher M2 macrophage levels. Furthermore, our results suggest that the HER2-negative and HER2-low groups possess a tumor-suppressive immune microenvironment compared to the HER2-high group. Our work explores the response variability of GCA patients to specific immune inhibitory checkpoint treatments ^13,14^. By identifying distinct tumor microenvironments and variations in immune inhibitory checkpoints across HER2 expression groups, we propose the CD47/SIRPA axis as potential targets for the HER2-negative and HER2-low groups. These newly identified immune inhibitory checkpoints provide promising therapeutic avenues for individualized treatment in GCA patients. Overall, our study offers valuable insights and resources for GCA and paves the way for precision medicine in its treatment.

## Acknowledgments

This work is supported by the National Key R&D Program of China (2012AA02A503, 2016YFC0901403 to L.D.W.), the National Natural Science Foundation of China (81872032, U1804262, 30025016 to L.D.W.; 81972571,81472234 to S.G.G.), the State Key Laboratory of Esophageal Cancer Prevention & Treatment Key R&D Program of Henan province (202305425 to L.D.W.), Henan Medical Science and Technology Research Project (LHGJ20190001 to X.K.Z. and L.D.W.), Henan Province Key R&D and Promotion Special (Science and Technology) Project (232102311032 to X.S. and L.D.W.), the Wallenberg Academy Fellow in Medicine from Knut and Alice Wallenberg foundation (2023.0046 and 2024.0166 to X. C.), the Swedish Research Council (2022-00658 to X.C.), the Swedish Cancer Foundation (21 1449Pj, 22 0491 JIA, 24 3484 Pj to X.C.). The funders played no role in the study design, data acquisition and interpretation, or decision to publish.

## Author contributions

L.D.W. and X.C conceived and designed the study. P.X analyzed all omics data. M.X.W., X.K.Z and X.S analyzed the correlation of omics and clinic-pathological data and follow-up data. X.M.L., X.K.Z and L.L.L analyzed the immunohistochemistry data. All the other authors collected clinical samples and related clinical information, performed experiments, and interpreted data. P.X and M.X.W wrote the materials and methods. X.C wrote the manuscript by collecting information from all authors. L.D.W. made critical comments and suggestions to finalize the manuscript. All authors read the manuscript.

## Declaration of generative AI and AI-assisted technologies

Neither AI nor AI-assisted technologies were used in the writing of this manuscript.

## Supplemental information

Supplementary Figures 1–32, and Tables S1–S26 are in separate files.

## Materials and Methods

### Patient and Public Involvement

Patients or the public were not involved in the design, or conduct, or reporting, or dissemination plans of our research.

### Ethics Statement

The ethics permit in this study about human patients was approved by the Ethics Committee of Zhengzhou University (Approval No: 2021-0019).

### Study population, tissue and blood collection

All the 128 patients clinically diagnosed with GCA from January 2019 to February 2020 were enrolled in this study, including 102 males with an mean age ± SD of 64.6 ± 7.5, and 26 females with an mean age of 64.7 ± 6.3. All the patients were from Henan, Hebei, and Shanxi provinces, northern China, the highest incidence areas for both GCA and esophageal cancer. None of these patients had undergone preoperative radiation therapy or chemotherapy, and they had no history of other malignant tumors. All the patients were performed radical cardiectomy of GCA. The tumor tissues and normal tissue adjacent to the tumor (NAT) to the tumor from GCA patients were all collected in the operating room, and all the samples were processed within half an hour after removal, and each sample of tumors and NATs (taken from tissue 3cm beyond the edge of the cancer tissue) was certified through pathological examination by two well-trained pathologists separately. Hematoxylin and eosin (H&E) stainings of NATs and their corresponding tumors were further examined to valide by two pathologists. The clinical characteristics and survival data were summarized in the analysis (**Table S1**). Staging of GCA was determined by two pathologists based on the eighth edition of the AJCC (American Joint Committee on Cancer’s Staging System). The last follow-up was at the end of March 2024. Fasting blood samples were meticulously obtained from GCA patients before surgical interventions during their hospitalizations. Each patient contributed two whole blood samples, with distinct processing purposes. The sample was drawn into an anticoagulant tube containing ethylenediaminetetraacetic acid (EDTA) for Whole Exome Sequencing (WES). This study implemented an integrative approach, encompassing whole exome sequencing, RNA sequencing and proteomics. The investigation utilized formalin-fixed paraffin-embedded (FFPE) tumor tissue blocks for immunohistochemistry and immunofluorescence staining.

### DNA extraction, library preparation, and whole exome sequencing

Whole exome sequencing (WES) was performed on 113 pairs of fresh-frozen tissues and blood (n = 112) or NATs (n = 1) from GCA. Genomic DNA from tumor tissue and blood was isolated using the DNeasy Blood & Tissue Kit (69504, Qiagen), following the manufacturer’s instructions. Subsequently, precise quantification of DNA concentration was carried out using the Qubit 2.0. DNA samples with a concentration of ≥20 ng/µL. A total amount of 0.6 μg genomic DNA per sample was fragmented to an average size of 180–280 bp and subjected to DNA library preparation using Illumina TruSeq DNA sample preparation kit. The Agilent SureSelect Human All ExonV5 Kit (5190-6209, Agilent Technologies) was used for exome capture according to the manufacturer’s instruction. In brief, DNA libraries were hybridized with liquid phase with biotin labeled probes from the Agilent SureSelect Human All ExonV5 Kit, then magnetic streptavidin beads were used to capture the exons of genes. Captured DNA fragments were enriched in a PCR reaction with index barcodes for sequencing. Final libraries were purified using AMPure XP beads (A63880, Beckman Coulter) and quantified using the Agilent high sensitivity DNA kit (5067-4626, Agilent Technologies). WES libraries were sequenced on Illumina Novaseq 6000 (Illumina) with 150 bp paired end mode according to the manufacturer instruction.

### RNA isolation, library preparation, and sequencing

RNA sequencing involved 54 pairs of fresh-frozen tissues and corresponding normal adjacent tissues (NAT) from GCA patients. Total RNA extraction from each sample employed the RNA Easy Fast Cell Total RNA Extraction Kit (DP451, Tiangen), following the manufacturer’s protocol. RNA integrity and total amount were assessed using the Agilent 2100 Bioanalyzer (G2939A, Agilent). For mRNA library construction, the NEB Next® Ultra RNA Library Prep Kit for Illumina (E7530L, NEB) was used. Polyadenylated mRNA was enriched from total RNA using oligo(dT) magnetic beads, followed by random fragmentation. The first cDNA strand was synthesized using random oligonucleotides as primers in the M-MuLV reverse transcriptase system. Subsequently, RNaseH degraded the RNA strand, and the second cDNA strand was synthesized using dNTPs in the DNA polymerase I system. The double-stranded cDNA underwent end-repair, A-tailing, and ligation to sequencing adapters. AMPure XP beads were employed to select cDNA fragments of approximately 370∼420 bp, followed by PCR amplification and purification to obtain the final library. Following library construction, initial quantification used the Qubit 2.0 Fluorometer, with subsequent library dilution to 1.5 ng/µl. The Agilent 2100 Bioanalyzer was then utilized to confirm the insert size of the library. Upon meeting the specified criteria, qRT-PCR quantified the effective library concentration (ensuring it exceeded 2nM) for quality verification. The final library underwent sequencing on the Illumina NovaSeq6000, generating 150 bp paired-end reads.

### Protein extraction and proteomics

Qualitative and quantitative assessment of proteins were conducted on 128 pairs of fresh-frozen tissues and corresponding NATs from GCA. Initial processing involved the cryogenic grinding of GCA tissues into cell powder using liquid nitrogen, followed by centrifugation. Subsequently, the cell powder underwent sonication on ice, employing a high-intensity ultrasonic processor (Scientz), and the resultant lysate was treated with lysis buffer (8 M urea, 1% Protease Inhibitor Cocktail). Centrifugation was employed to eliminate residual debris, and the supernatant, containing proteins, was collected. Protein concentration was determined using a BCA kit following the manufacturer’s protocol. The subsequent steps encompassed trypsin digestion, TMT/iTraq labeling, high-performance liquid chromatography (HPLC) fractionation utilizing a Thermo Betasil C18 column (5 μm particles, 10 mm ID, 250 mm length), and liquid chromatography-tandem mass spectrometry (LC-MS/MS) analysis. The acquired data were processed using the MaxQuant search engine (v.1.5.2.8) against the Homo_sapiens_9606 database.

### Immunohistochemistry (IHC) of proteins

IHC of HER2, SIRPA, and PDCD1LG2 proteins was according to the same method as our previous study^23^. In brief, 5 µm thick formalin fixed paraffin-embedded GCA tissue sections were first deparaffined with xylene 15 min for three times, then were dehydrated through 100% alcohol, 85% alcohol and 75% alcohol for 5 min each, followed by distilled water rinsing for 5 min. The epitope retrieval is performed in the microware by putting the tissue into citrate buffer (pH 6.0). After the epitope retrieval, the tissue section is rinsed in Phosphate-Buffered Saline buffer (PBS, pH 7.4). After blocked with 3% bovine serum albumin (BSA) 30 min at room temperature, the tissues were incubated with primary antibodies, Rabbit anti-ERBB2 antibody (1:100 dilution, K00125, Roche), Rabbit anti-SIRPA (1:100, ab53721, Abcam), or Rabbit anti-PDCD1LG2 (1:100, ab200377, Abcam), overnight at 4 °C. In the next day, the washing is performed with PBS buffer for three times, 15 min each. The secondary antibody (1:1000 dilution, Horseradish Peroxidase, HRP marked, PV-9000, ZSGB-BIO) was incubated for 50 min at room temperature. After the secondary antibody incubation, the washing is performed with PBS buffer three times on shaker, 15 min each. The tissue is stained with the Harris Hematoxylin for 3 min. At last, the tissue section was mounted and imaged. The identification of HER2, SIRPA and PDCD1LG2 positive cells was validated by two senior pathologists.

### HER2 Immunohistochemical Scoring Criteria

The HER2 scoring criteria were established and validated by two senior pathologists. Tumor specimens were evaluated for membranous HER2 staining intensity using the following criteria: HER2 0+: No reaction or less than 10% of tumor cells exhibited membranous HER2 staining. HER2 1+: Faint or faintly visible membranous HER2 staining in at least 10% of tumor cells, with only partial cell membrane HER2 staining observed.

HER2 2+: Weak to moderate basolateral, lateral, or complete membranous HER2 staining observed in at least 10% of tumor cells. HER2 3+: Basolateral, lateral, or complete membranous HER2 staining present in at least 10% of tumor cells.

The HER2 scoring criteria were meticulously applied to assess membranous HER2 staining intensity in tumor cells, ensuring consistency through validation by two senior pathologists.

### Immunofluorescence staining of CD163

The immunofluorescence assay was conducted on paraffin-embedded (FFPE) sections obtained from GCA specimens. Initially, FFPE slices of GCA paraffin-embedded sections underwent baking in an oven for paraffin fixation. Immediate dewaxing of the paraffin slices followed, involving sequential treatments with an eco-friendly dewaxing solution, anhydrous ethanol, 90% alcohol, 70% alcohol, and distilled water. Dewaxed slices were then subjected to low-temperature fixation. The fixing solution was meticulously prepared in a dedicated fixing box and heated in a microwave until boiling, ensuring the absence of air bubbles. Following fixation, rapid cooling of the slices was achieved using ice cubes. Subsequent steps included the removal of the fixing solution with TBST, application of an appropriate sealing solution, removal of excess sealing solution, and sequential addition of primary and secondary antibodies targeting CD163 (ab182422, Abcam, UK). Incubation at room temperature allowed for antibody binding. Afterwards, residual primary and secondary antibody solutions were washed off using TBST. The cleaned slides were treated with corresponding fluorescent dyes, followed by incubation at room temperature. Subsequent TBST washes removed any excess dye. DAPI working solution was applied for nuclear staining, with thorough cleaning to eliminate residual DAPI, dewaxing solution, xylene, or debris. The slides were then sealed for further analysis.

The image acquisition was performed using the FUISON model full-spectrum scanner from the American company Akoya, capturing all image information of the entire tissue on the slides. Image analysis was conducted using VISIOPHARM software, initially focusing on tissue contour recognition. Experienced pathology experts further delineated tumor tissue regions by overlaying the mIHC-stained images onto the corresponding HE images. During this process, the software calculated the areas of the entire tissue and tumor regions. The VISIOPHARM software employed its inherent deep learning algorithms for cell segmentation, facilitating the determination of total cell counts within the tissue. The Phenplex Guided Workflow functionality in VISIOPHARM software was utilized for cell phenotype identification, enabling the quantification of individual positive cells for each channel and the determination of co-positive cells between channels. Subsequently, statistical calculations leveraging the obtained data on the entire tissue and tumor area, total cell count, positive cell counts for each channel, and co-positive cell counts between channels were conducted. These calculations included the determination of positivity rates, area density, as well as mean, standard deviation, and inter-batch P-values among samples.

### TCGA data processing

Transcriptome profiling data from The Cancer Genome Atlas Program (TCGA) were downloaded from the GDC Data Portal with the project IDs ESCA (Esophageal Carcinoma) and STAD (Stomach Adenocarcinoma). The expression data, represented by TPM (Transcripts Per Million) normalized values, were extracted from each sample. Subsequently, all samples were merged into a unified TPM matrix.

### Patient Stratification Based on ERBB2/HER2 Expression

#### ERBB2 RNA Expression Level

For TCGA patients (ESCC, EAC, STAD) and GCA patients (n = 54), individuals were stratified based on ERBB2 RNA levels (TPM) using three strategies: upper vs. lower median, top vs. bottom quantile, and natural separation (very high vs. rest) to distinguish between ERBB2 high- and low-expression groups.

#### Ratio of ERBB2 RNA Expression Between Tumor and normal tissue adjacent to the tumor (NAT)

For GCA patients (n = 54), individuals were divided into two groups—high and low—based on the median ratio of ERBB2 RNA expression levels between the tumor NAT.

#### Ratio of HER2 Protein Expression Between Tumor and normal tissue adjacent to the tumor (NAT)

For GCA patients (n = 128), individuals with NA values for HER2 expression were assigned to the HER2-negative group. The remaining patients were divided into two groups—high and low— based on the median value of the ratio between tumor and NAT. All omic data analyses were based on this stratification.

### Differential gene analysis in EAC cell line OE19

RNA-Seq data for the EAC cell line OE19, both with and without treatment with the HER2 inhibitor lapatinib, as well as following HER2 siRNA silencing, were obtained from ArrayExpress (E-MTAB-10304)^35^. The gene expression levels were further normalized using transcripts per million (TPM). Differential gene analysis was performed with DESeq2^38^. The significantly changes genes are with a cutoff of |log2(FoldChange)| >= 1 and FDR <= 0.

### Proteomics data processing and statistical analysis

The raw LC-MS datasets were initially searched against a database and converted into matrices containing the reporter intensity of peptides across samples. Subsequently, the relative quantitative value for each modified peptide was determined through the following procedures: First, the intensities of modified peptides (I) were centralized and transformed into relative quantitative values (U) for each modified peptide in each sample: Uij = Iij / Mean(Ij), where i represents the sample, and j denotes the modified peptide. Second, the relative quantitative values (Q) of modified peptides were corrected by the median value: Qij = Uij / Median(Ui). Proteins with 50% NA values across all samples were excluded. The protein expression ratio (R) between tumor samples (T) and normal adjacent samples (NAT) was calculated with Rmn = Tmn/NATmn, where m denotes each protein, and n denotes each sample. Differential analysis among the three groups with the “One v.s. Rest” strategy was conducted on the Rmn matrix using the wilcox.test function in R. Additionally, differential analysis between tumors and NATs was performed based on tumor and NATs using the wilcox.test function in R. Proteins were considered differentially expressed if their |FoldChange| was greater than or equal to 1.5, and the p-value was less than or equal to 0.05.

### Whole exome sequencing data processing

The pair-end sequencing data generated from Illumina NovaSeq 6000 were trimmed using fastp v0.23. Subsequently, the trimmed data were mapped to the hg38 reference genome using bwa mem v0.7.17^39^. The SAM files were converted into BAM format and sorted by samtools v1.16.1. Duplicate reads were marked using “GATK4.3 MarkDuplicates.” Base quality scores were recalibrated using “GATK4.3 BaseRecalibrator” and “GATK4.3 ApplyBQSR,” employing known variants from the dbSNP database (build 146) ^8^ and the 1000 Genomes project^40^.

### Somatic short variant calling

The somatic short variants, including SNVs and InDels, were called using GATK Mutect2^41^, leveraging matched normal samples as reference background and utilizing the “af-only-gnomad” file from the Gnomad datasets as germline resources. Subsequently, “GATK GetPileupSummaries” and “GATK CalculateContamination” were employed to compute cross-sample contamination. Finally, “GATK FilterMutectCalls” was used to filter short variants with the contamination table, retaining mutations with a “PASS” filter flag for subsequent analysis. Copy number alterations (CNAs) were identified using both CNVkit/0.9.8^42^ and FREEC-11.6^43^, utilizing tumor-matched normal samples as the reference background. Copy numbers equal to or less than 1 were categorized as CNA loss, while copy numbers equal to or greater than 3 were considered as CNA gain. CNAs identified by both software tools were retained for subsequent analysis.

### SNVs and InDels analysis

All SNVs and InDels identified by GATK were annotated using snpEff v5.1. Subsequently, the annotated SNVs and InDels were transformed into Mutation Annotation Format (MAF) using a custom script which is deposited in https://github.com/pengweixing/GCA_multi-omics, following the guidelines provided in https://docs.gdc.cancer.gov/Encyclopedia/pages/Mutation_Annotation_Format_TCGAv2/.

For each sample, variant types including “Missense_Mutation”, “Nonsense_Mutation”, “Splice_Site”, “Frame_Shift_Del”, “Frame_Shift_Ins”, “In_Frame_Del”, and “In_Frame_Ins” were selectively extracted. The identified SNVs and InDels from all samples were subsequently aggregated. The distribution of mutated genes within three groups was visualized using the R package “ComplexHeatmap” through a waterfall plot. Only mutated genes that align with entries in the Cancer Gene Database (COSMIC) were retained for further analysis. The tumor mutation burden (TMB) was quantified as the number of SNVs per million nucleotides in the coding region. Synonymous alterations were also counted separately for the purpose of reducing sampling noise.

### CNA analysis

Copy number alterations (CNAs) across the entire genome for each group were depicted using the R package Circlize. Additionally, CNAs in specific regions for each group were visualized utilizing the R package karyoploteR. The genome was divided into bins with a 1Mb window, and the CNAs were counted in each bin for three groups, with a subsequent application of the Chi-square test. CIN was estimated through genome integrity index (GII), the proportion of altered genome segments^44^. Weighted Genome Instability Index (wGII) was also caculated following a previous report^28^.

The formula is as follows (where ’n’ represents all autosomes):

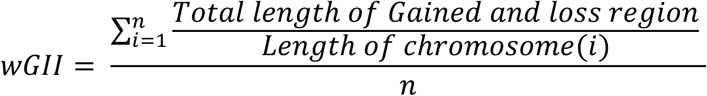

The number of breakpoints representing the Chromosomal Instability Index (CINindex) was calcuated with following formula:

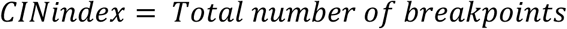

wCINindex (Weighted Chromosomal Instability Index) was calcuated by considering the number of breakpoints per million base pairs in each chromosome.

The formula is as follows (where ’n’ represents all autosomes):

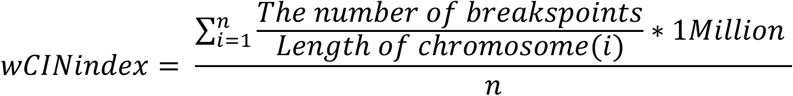

### RNA-seq data processing and differential analysis

The paired-end sequencing data were processed through the nf-core/rnaseq 3.10.1 pipeline. Briefly, fastq files were subjected to trimming using Trim Galore v0.6.10, followed by alignment to the hg38 genome using STAR v2.7.0. Sorting and indexing of the resulting BAM file were performed using samtools v1.16.1 as. Subsequently, Salmon was employed to quantify the expression levels of individual genes. The gene expression levels were further normalized using transcripts per million (TPM).

The categorization of groups in RNA-seq data aligned with the HER2-High, HER2-Low, and HER2-Negative groups identified in proteomics. Differential analysis was conducted within tumor samples with DESeq2^38^, comparing HER2-High versus Rest, HER2-Low versus Rest, and HER2-Negative versus Rest. A cutoff of |FoldChange| greater than or equal to 1.5 and a false discovery rate (FDR) less than or equal to 0.05 was applied to identify differentially expressed genes. Additionally, differential analysis was performed between HER2-High versus NATs, HER2-Low versus NATs, and HER2-Negative versus NATs, with a cutoff of |log2(FoldChange)| >= 1 and FDR <= 0.05 to filter differentially expressed genes.

### Cellular composition analysis

CIBERSORTx^33^ analysis was conducted in accordance with previously established procedures. The docker-based tool of CIBERSORTx was utilized to estimate the relative fractions of 22 immune cell sub-populations, employing the LM22 reference gene expression signatures. Comparisons were conducted among three groups for each cell type. Statistical differences were calculated with “t-test” function in R.

### Gene Ontology (GO) enrichment and Gene set enrichment analysis (GSEA)

Differentially expressed genes and proteins identified in transcriptomics and proteomics datasets were chosen for Gene Ontology (GO) enrichment analysis. The enrichGO function within the R package clusterProfiler v4.0 was employed for conducting gene ontology analysis, with FDR of 0.05 applied to filter significant GO terms. For Gene Set Enrichment Analysis (GSEA), 50 hallmark gene sets from the Human MSigDB collections were adopted. The genes for input were ranked based on their weights which calculated with “*Geneweight* = *sign*(*foldchange*) * -log10(*FDR*)”. “GSEA” function in clusterProfiler package was adopted for GSEA analysis. A significance threshold of FDR = 0.05 was applied to filter GSEA hallmarks.

### *ERBB2* grouping patients based on scRNA-seq data

Raw reads (R2) from the scRNA-seq data were treated as bulk RNA-seq, as the 10x barcodes and UMIs were located in R1. We used R2 as input for the bulk RNA-seq pipeline, employing nf-core rnaseq (version 3.14.0)^45^. Gene expression was normalized using TPM, and samples were divided into *ERBB2*-high and *ERBB2*-low groups based on expression levels.

### Tissue dissociation and preparation of single-cell RNA-Seq

The tumor samples obtained were washed 2-3 times in 4°C precooled 1× PBS on ice, transferred to a sterile RNase-free culture dish, and cut into small pieces (about 0.5 mm²) using 10 cm surgical scissors. The tissues were then washed with 1× PBS to remove as much non-target tissue as possible, such as blood stains, fatty layers, and connective tissue. The tissue samples were incubated in a constant-temperature water bath or shaker at 37°C with either a MACS Human Tumor Dissociation Kit (130-095-929, Miltenyi Biotec). Incubation was generally terminated once the digestion solution became turbid and the tissue mass had dissolved. Next, a 40 μm cell sieve was used to filter the suspension, or a 100 μm/70 μm cell filter if the tissue mass was larger. The cell suspension was typically centrifuged at 300 rpm at 4°C for 5 minutes. The supernatant was discarded, and 1 mL of pre-cooled 1× PBS was added to re-suspend the cells. Then, 3 mL of pre-cooled erythrocyte lysis buffer was added to the tube, mixed evenly, and incubated at 4°C for 5-10 minutes. After incubation, the suspension was centrifuged at 300 rpm at 4°C for 5 minutes, the supernatant was removed, and 1 mL of precooled 1× PBS was added to fully re-suspend the cells. The cell concentration was measured using a cell counter, and adjusted as needed to reach a target concentration of 700-1200 cells/μL, with cell viability above 90% and cell clumping below 15%. Once the final cell concentration and viability were achieved, the cells were placed on ice, and a 10X Genomics single-cell transcriptome chip experiment was initiated within 30 minutes.

### Single-cell RNA-seq library preparation and sequencing

We prepared single cell RNA-seq libraries with Chromium Next GEM Single Cell 3ʹ Reagent Kits v3.1 on the Chromium Controller(10× Genomics).We prepared single cell suspensions from cultured cell lines and single cells were suspended in PBS containing 0.04% BSA.The cell suspension was loaded onto the Chromium Next GEM Chip G and ran the Chromium Controller to generate single-cell gel beads in the emulsion(GEMs) according to the manufacturer’s recommendation.captured cells were lysed and the released RNA was barcoded through reverse transcription in individual GEMs. Barcoded, full-length cDNA was generated and libraries were constructed according to the performer’s protocol. The quality of libraries was assessed by Qubit 4.0 and the Agilent 2100. Sequencing was performed on the Illumina NovaSeq 6000 with a sequencing depth of at least 50,000 reads per cell and 150 bp (PE150) paired-end reads.

### Quality control, data normalization and intergration of scRNA-seq data

Raw reads from the 10x Genomics single-cell RNA-seq platform were demultiplexed and aligned to the human reference genome (GRCh38) using CellRanger (version 7.2.0). The filtered cell-to-gene matrix was processed in Seurat v5.0.3^46^. Cells were included if they expressed between 500 to 7,000 genes, had UMIs between 100 and 50,000, and less than 20% mitochondrial reads. Data normalization was performed with Seurat’s ‘NormalizeData’ function, followed by scaling using ‘ScaleData’ function. Dimensionality reduction was conducted via ‘RunPCA’, and samples were integrated to remove biological variations using the ‘IntegrateLayers function.

### Cell type identification from single cell RNA-Seq

We conducted cell clustering using Seurat’s ‘FindNeighbors’ and ‘FindClusters’ functions. To visualize the clusters, we applied the ‘RunUMAP’ function with “dims = 1:30”. For cell type annotation, we utilized esophageal adenocarcinoma (EAC) scRNA-seq data from the LUD2015-005 project (https://zenodo.org/records/8083316) as a reference^47^. This reference dataset was used to identify key cell compartments—Epithelium, Lymphocytes, Phagocytes, and Stroma—by applying the ‘FindTransferAnchors’ and ‘TransferData’ functions with “dims = 1:30” and normalization.method = ’SCT’. Additionally, plasma cells, B cells, T cells, endothelial cells, fibroblasts, mast cells, pericytes, muscle cells, and myeloid cells were annotated using the EAC reference. We extracted T cells and myeloid cells for further dimensionality reduction and clustering. Cell-type specific marker genes were identified using Seurat’s ‘FindAllMarkers’ function. By combining marker genes and leveraging GPT-4’s advanced annotation capabilities^48^, we refined the analysis of T cells and myeloid cells, identifying subtypes such as macrophages, monocytes, neutrophils, dendritic cells, CD8+ T cells, CD4+ T cells, regulatory T cells, and NK cells.

### Differential gene analysis in single cell RNA-seq

For specific cell types, such as CD4+ T cells, CD8+ T cells, myeloid cells, M2 macrophages, and epithelial tumor cells, we downsampled the cell numbers to balance the *ERBB2*-High and *ERBB2*-Low groups. Differential gene expression analysis was performed using Seurat’s^46^ ‘FindMarkers’ function. Genes with an adjusted p-value < 0.05 and an absolute log2(fold change) > 0.25 were considered significant. Gene ontology and KEGG pathway analyses were conducted using the ‘enrichGO’ and ‘enrichKEGG’ functions from the clusterProfiler v4.7.1 package. GSEA analysis was performed using ranked genes based on fold changes with the ‘GSEA’ function of clusterProfiler v4.7.1.

### Pseudotime analysis

Monocle2 was employed to infer epithelial cell developmental trajectories^49^. Seurat objects were converted to CellDataSet for input into Monocle2, and differential genes between normal and tumor epithelial cells were identified using ‘FindAllMarkers’ (p-adjusted < 0.05, log2(fold change) > 1). Dimensionality reduction was performed with ‘DDRTree’ function, and cells were ordered with the ‘orderCells’ function. Trajectories were visualized using ‘plot_cell_trajectory’, and cell fate was analyzed using the branched expression analysis modeling (BEAM) function with branch_point=1. Heatmaps of cell fates were generated with ‘plot_genes_branched_heatmap’ with parameters of ‘branch_point = 1, num_clusters = 4’. The genes associated with each cell fate corresponded to the clusters shown in the heatmap. These genes were extracted for Gene Ontology (GO) and KEGG pathway enrichment analysis using the ’enrichGO’ and ’enrichKEGG’ functions from the clusterProfiler package v4.7.16.

### CNV estimation in epithelial cells

We used the InferCNV package (version 1.14.2)(https://github.com/broadinstitute/inferCNV) with default parameters to infer copy number variations (CNVs) in epithelial cells, using lymphocytes and phagocytes as reference. CNVs were defined by copy number values greater than 1.05 or less than 0.95. A CNV score for each cell was calculated based on the proportion of genes affected by CNVs. To classify epithelial tumor cells, we applied the ’optbin’ function in Optbin v1.3 package to determine an optimal threshold that minimized the squared error, dividing the cells into CNV-high and CNV-low groups. Cells with CNVs above the threshold were classified as CNV-high, and those below as CNV-low. For each sample, we calculated the proportions of CNV-high and CNV-low cells. A two-sided Wilcoxon test was performed to compare the proportions of CNV-high cells between *ERBB2*-High and *ERBB2*-Low groups.

### Cell-cell communication analysis

Cell-cell communication analysis was conducted using CellChat v2.1.2^50^ with the ‘Secreted Signaling’ database. For both the *ERBB2*-High and *ERBB2*-Low groups, we first identified overexpressed genes using the ‘identifyOverExpressedGenes’ function, followed by identification of interactions using the ‘identifyOverExpressedInteractions’ function. We then computed communication probabilities and constructed the cellular communication network with the ‘computeCommunProb’ function. Low-cell-number communications were filtered out using the ‘filterCommunication’ function with a minimum cell threshold of 10. The ‘computeCommunProbPathway’ function was employed to calculate communication at the signaling pathway level. To compare the two groups, we merged the *ERBB2*-High and *ERBB2*- Low CellChat objects using the ‘mergeCellChat’ function. Differential interaction strengths between the groups were visualized using circle plots with ‘netVisual_diffInteraction’ function and heatmaps with ‘netVisual_heatmap’ function. We compared overall information flow between cell types using a bar plot generated by the ‘rankNet’ function. Dysfunctional signaling pathways were identified via differential expression analysis using the ‘identifyOverExpressedGenes’ function. Communication probabilities for ligand-receptor pairs between cell types were visualized using bubble plots with ‘netVisual_bubble’ function, and upregulated or downregulated ligand-receptor interactions were illustrated with a chord diagram with ‘netVisual_chord_gene’. Additionally, interactions of immune checkpoint ligands and receptors across cell types were analyzed using CellphoneDB v5.0.1^51^ with default parameters. P-values and means for immune checkpoint gene pairs were extracted and presented in a bubble plot.

### Spatial Transcriptomics

Formalin-fixed, paraffin-embedded (FFPE) samples from GCA tumors were cut to perform H&E staining (hematoxylin and eosin staining). The H&E staining of tumor sections was reviewed by two senior pathologists, and tumor-immune border regions were punched to create a tissue microarray for Spatial Transcriptomics. Nine tissue microarrays were included in each Spatial Transcriptomics analysis. The tissue microarrays were prepared according to the Visium CytAssist Spatial Gene Expression for FFPE-Tissue Preparation Guide (CG000518, 10× Genomics, Pleasanton, CA, USA). Tissue sections with a thickness of 5 μM were prepared for subsequent experiments, including deparaffinization, H&E staining, and decrosslinking. Library construction was performed using the Visium CytAssist Spatial Gene Expression Reagent Kits (PN-1000520, 10x Genomics), following the manufacturer’s protocol. Libraries were sequenced using the Illumina NovaSeq 6000 platform with the PE150 sequencing mode.

### Quality control and normalization of 10x spatial transcriptomics

Raw reads from the 10x Genomics Visium platform were demultiplexed and aligned to the human reference genome (GRCh38) using SpaceRanger v3.0.0 software. The filtered cell-to-gene matrix generated by SpaceRanger was then imported into Seurat for further analysis. High-quality spots were defined as those with nCount_Spatial > 2000, nFeature_Spatial > 500, and percent.mt < 20. Each 10x slide contained nine tissue sections arranged in a 3x3 grid. For each section, we calculated the proportion of high-quality spots and retained sections with over 90% high-quality spots. One lymph node metastasis section was removed, along with another section primarily composed of muscle cells and a section that was only partially covered with tissue. The remaining tissue sections were normalized using Seurat’s SCTransform function for downstream analysis.

### Cell type decomposition of 10x spatial transcriptomics

We performed cell type decomposition for each tissue section in the 10x spatial slide by integrating fine-annotated scRNA-seq data with spatial transcriptomics data using the cell2location v0.1.3 software^52^. Both datasets were normalized using Seurat’s SCTransform function, with the scRNA-seq data serving as the reference. Cell type signatures were estimated from the reference data using a Negative Binomial (NB) regression model with a maximum of 250 epochs. Spatial mapping was then conducted to estimate cell types in each spot based on the reference signatures, using a maximum of 10,000 epochs. The resulting cell abundance estimates were loaded into Seurat for visualization. Cell type composition in each tissue section was determined by normalizing the abundance of each cell type relative to the total cell abundance. Comparisons of cell type abundances between the *HER2*-High, *HER2*-Low, and *HER2*-Neg groups were conducted using a two-sided Wilcoxon test.

### Colocalization analysis

To investigate ligand-receptor gene interactions in the 10x spatial transcriptomics data, we first binarized the expression of each gene and the presence of each cell type. Macrophages, dendritic cells, and monocytes were grouped together as myeloid cells. A cell type was considered present in a spot if its cell2location-estimated abundance exceeded the median for that section. Similarly, a gene was considered expressed in a spot if its cell2location-estimated abundance was greater than one for that section. We counted the spots where both epithelial tumor cells and myeloid cells were either co-localized or not. Subsequently, in spots where these two cell types were co-localized, we identified spots where both ligand and receptor genes were co-expressed or not. Comparisons of co-localized spots among the HER2-High, HER2- Low, and HER2-Neg groups were made using a two-sided Wilcoxon test.

### Prognosis analysis

Survival analysis was conducted using the R package survival v.3.5-7 and survminer v.0.4.9. The Kaplan-Meier method was employed to estimate overall survival time, and statistical differences were assessed with log-rank test.

### Data processing and visualization

In addition to the previously mentioned R packages, the data processing and visualization also incorporated other tools, including dplyr v1.1.4, stringr v1.5.1, pheatmap v1.0.12, ggplot2 v3.4.4, ggpubr v0.6.0, and ggvenn v0.1.10.

## Code availability

The code to perform all analyses and regenerate all the figures in this study is provided at https://github.com/pengweixing/GCA_multi-omics.

## Data availability

All raw data from this study is deposited in the China National Center for Bioinformation with access number of HRA006640.

TCGA data that used in this study were downloaded from https://portal.gdc.cancer.gov/projects/TCGA-ESCA and https://portal.gdc.cancer.gov/projects/TCGA-STAD.

RNA-Seq data for the EAC cell line OE19, both with and without treatment with the HER2 inhibitor lapatinib, as well as following HER2 siRNA silencing, were obtained from ArrayExpress (E-MTAB-10304).

