## supplementary figures for "Multi-Omics Profiling Reveals Gene Signatures and Therapeutic Targets in HER2- Guided Gastric Cardia Adenocarcinoma Patients"

### Supplementary information:

#### Supplementary Figures 1-32

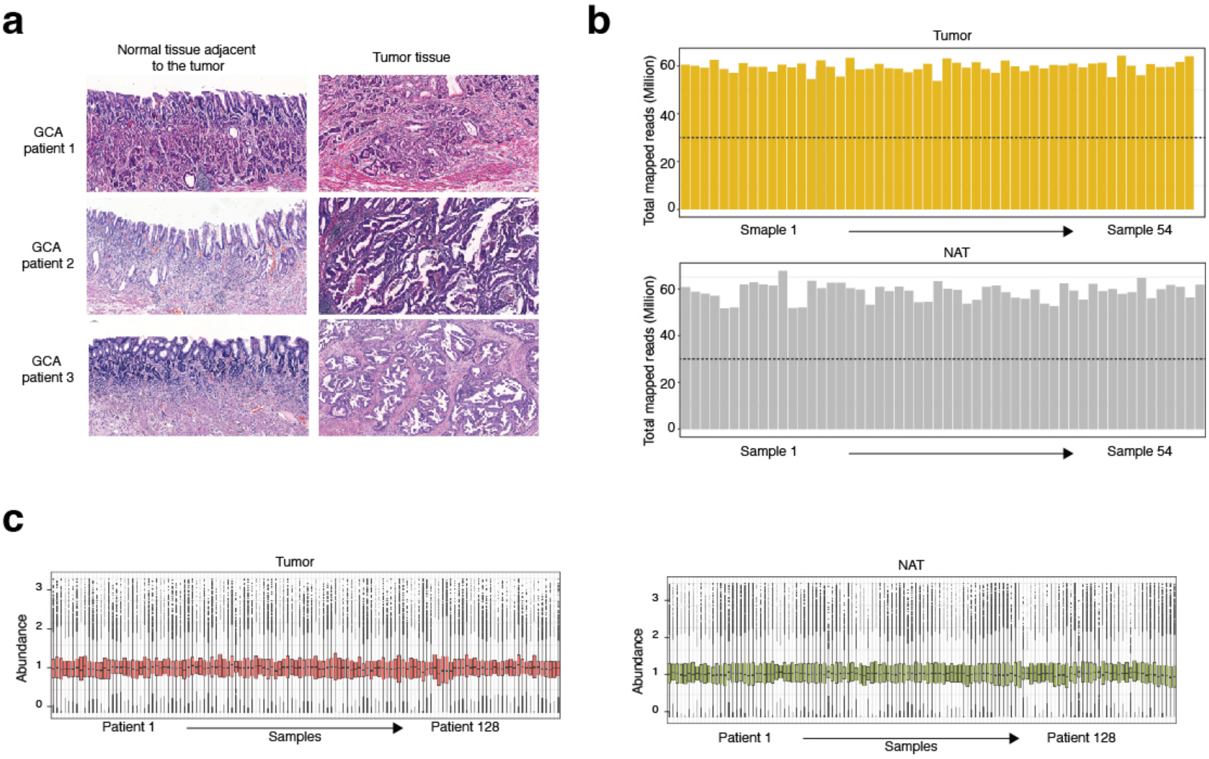


**Supplementary Figure 1: RNA and Protein measurements of tumor and normal adjacent tissue (NAT) in the GCA cohort.**

- **a.** Representative H&E (Hematoxylin and Eosin) staining of tumor and normal adjacent tissue (NAT).
- **b.** Quantification of mapped sequencing reads from tumor and NAT.
- **c.** Normalized protein levels in the tumor and NAT

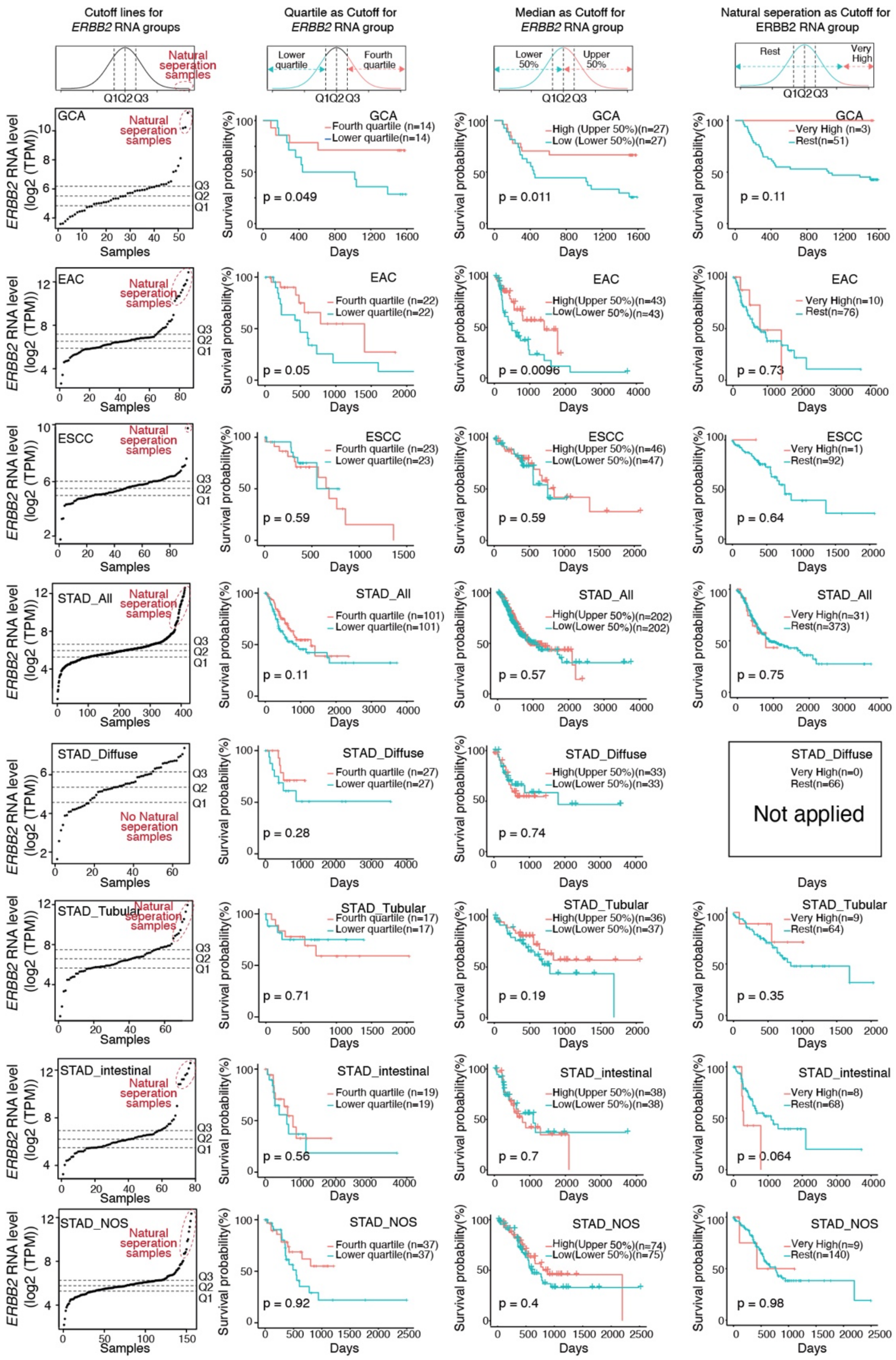

**Supplementary Figure 2: Prognostic analysis in EAC, GCA, ESCC, and STAD patients with different cutoffs for ERBB2 RNA expression levels.**

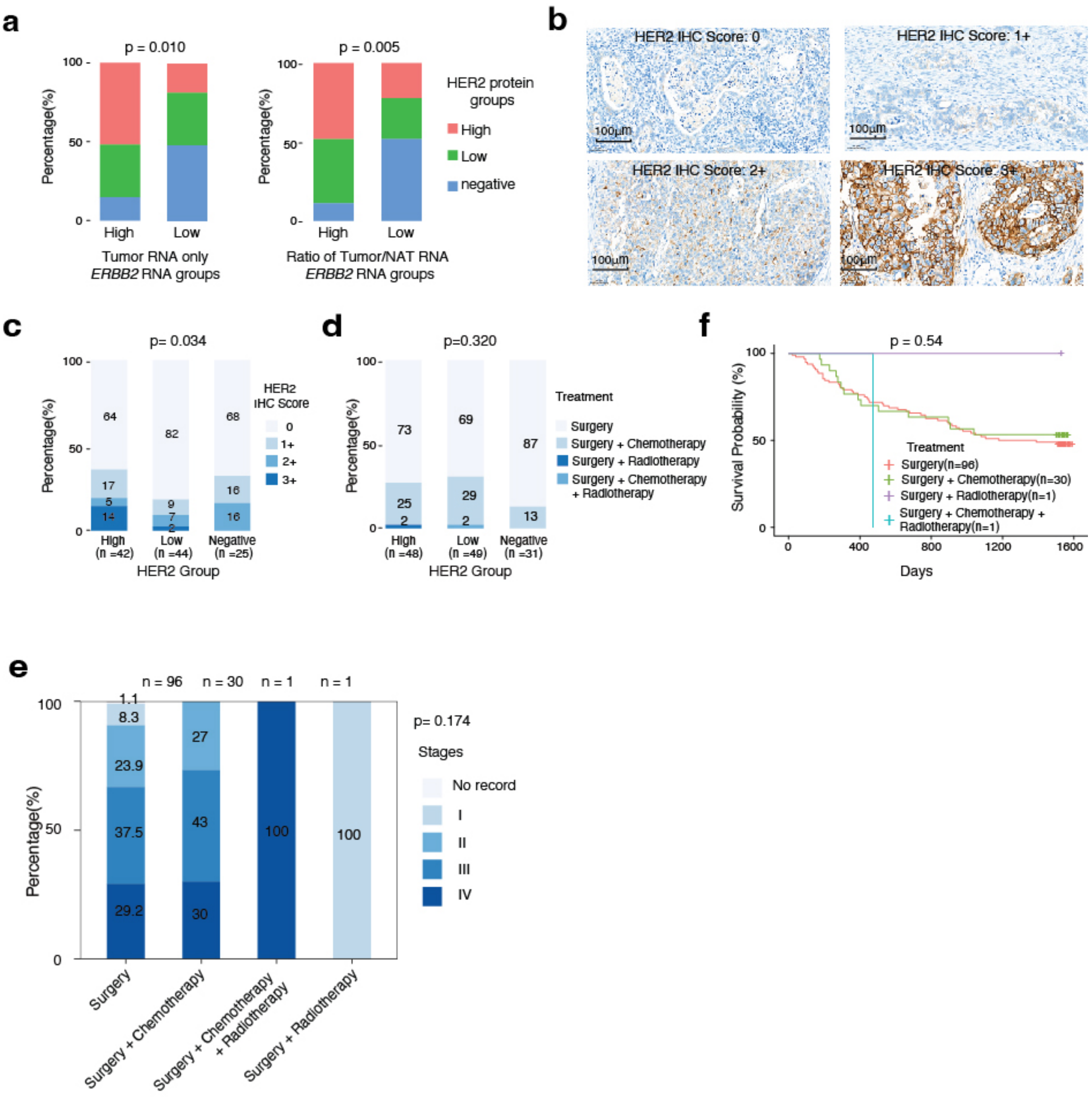


**Supplementary Figure 3: Association of GCA patients with grouping between HER2 RNA and HER2 protein.**

- **a.** Comparison of GCA patient stratification based on *ERBB2* RNA level, HER2 protein ratio between tumor and NAT, and *ERBB2* RNA ratio between tumors and NATs.
- **b.** Representative immunohistochemistry (IHC) staining of HER2 with scores of 3+, 2+, 1+, and 0+.
- **c.** The relationship between IHC staining HER2 scores and HER2 groups, along with the quantification of HER2 protein expression using mass spectrometry.
- **d.** The distribution of patients with different therapeutic treatments in three HER2 groups.
- **e.** Relationship between tumor stages and therapeutic strategies in the GCA cohort.
- **f.** Kaplan-Meier survival curves of different therapeutic treatments in our GCA cohort.

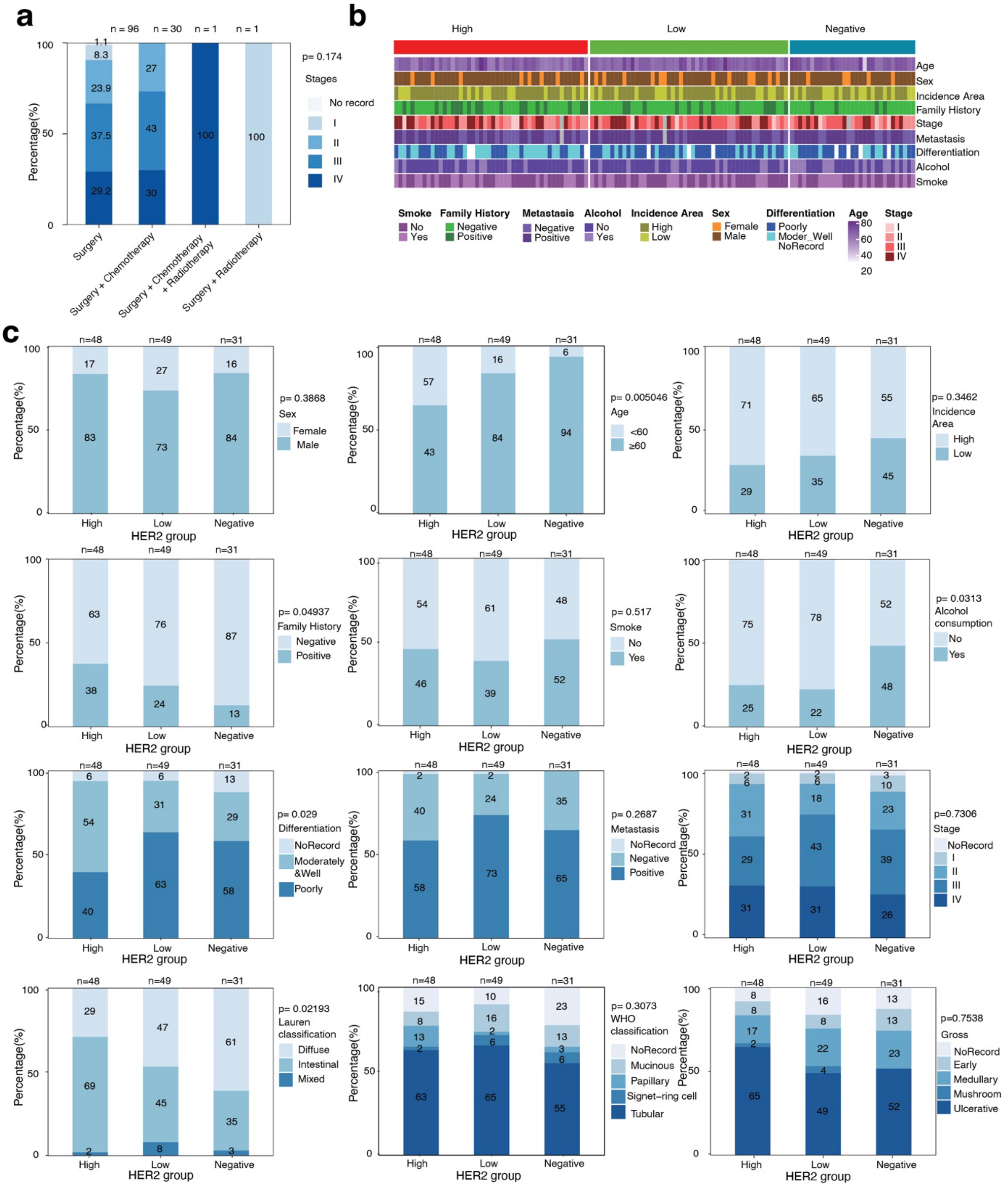


**Supplementary Figure 4: Clinical-pathological correlation with HER2 grouping in the GCA cohort.**

- **a.** Distribution of clinical pathological features in the GCA cohort.
- **b.** Relationship between HER2 grouping and clinical pathological features in the GCA cohort.

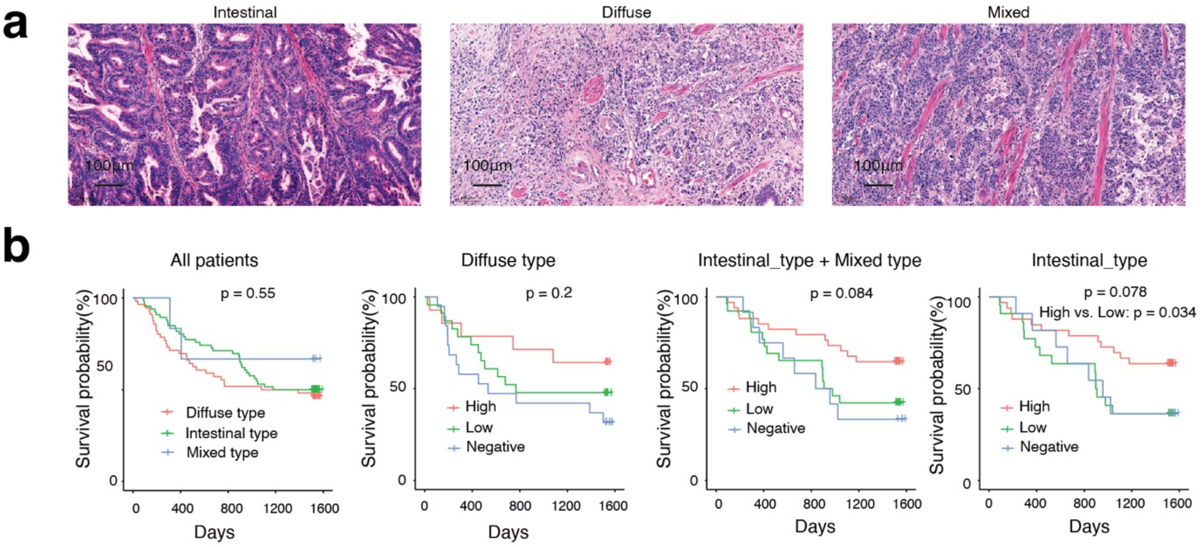


**Supplementary Figure 5: Lauren classification, HER2 grouping and prognosis of GCA patents.**

**a.** Representative H&E (Hematoxylin and Eosin) staining of different Lauren classifications of GCA patients.

**b.** Prognosis of GCA patients in different groups based on Lauren classification (left panel) and prognosis of GCA patients from different Lauren classifications within different HER2 groupings (right three panels).

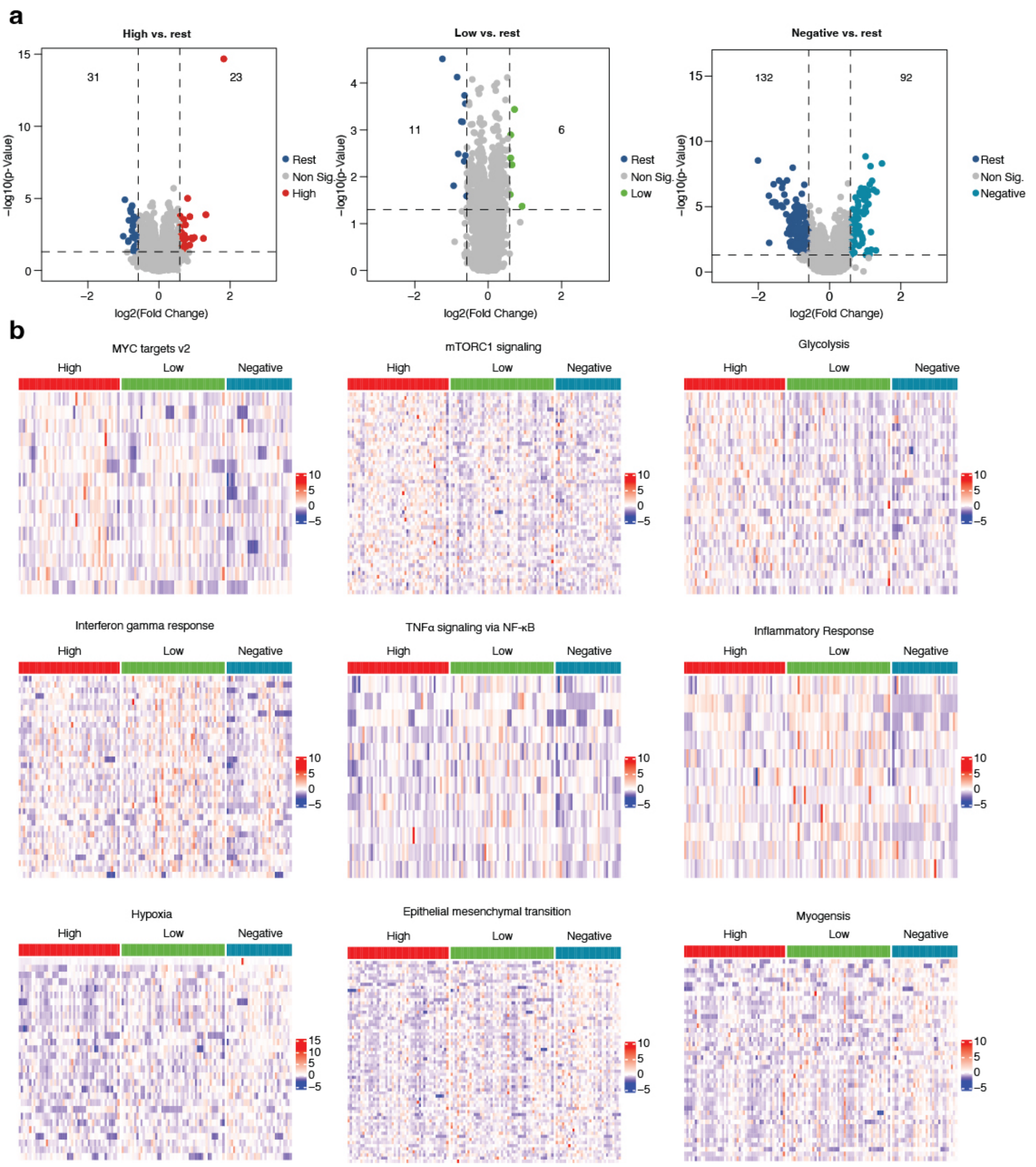


**Supplementary Figure 6: Identification of differential protein expression in three HER2 groups of GCA.**

- **a.** Volcano plots represent identified differential proteins expressions in each HER2 group from tumor only; number of proteins is indicated.
- **b.** Heatmaps show the enrichment of hallmark genes for each HER2 group.

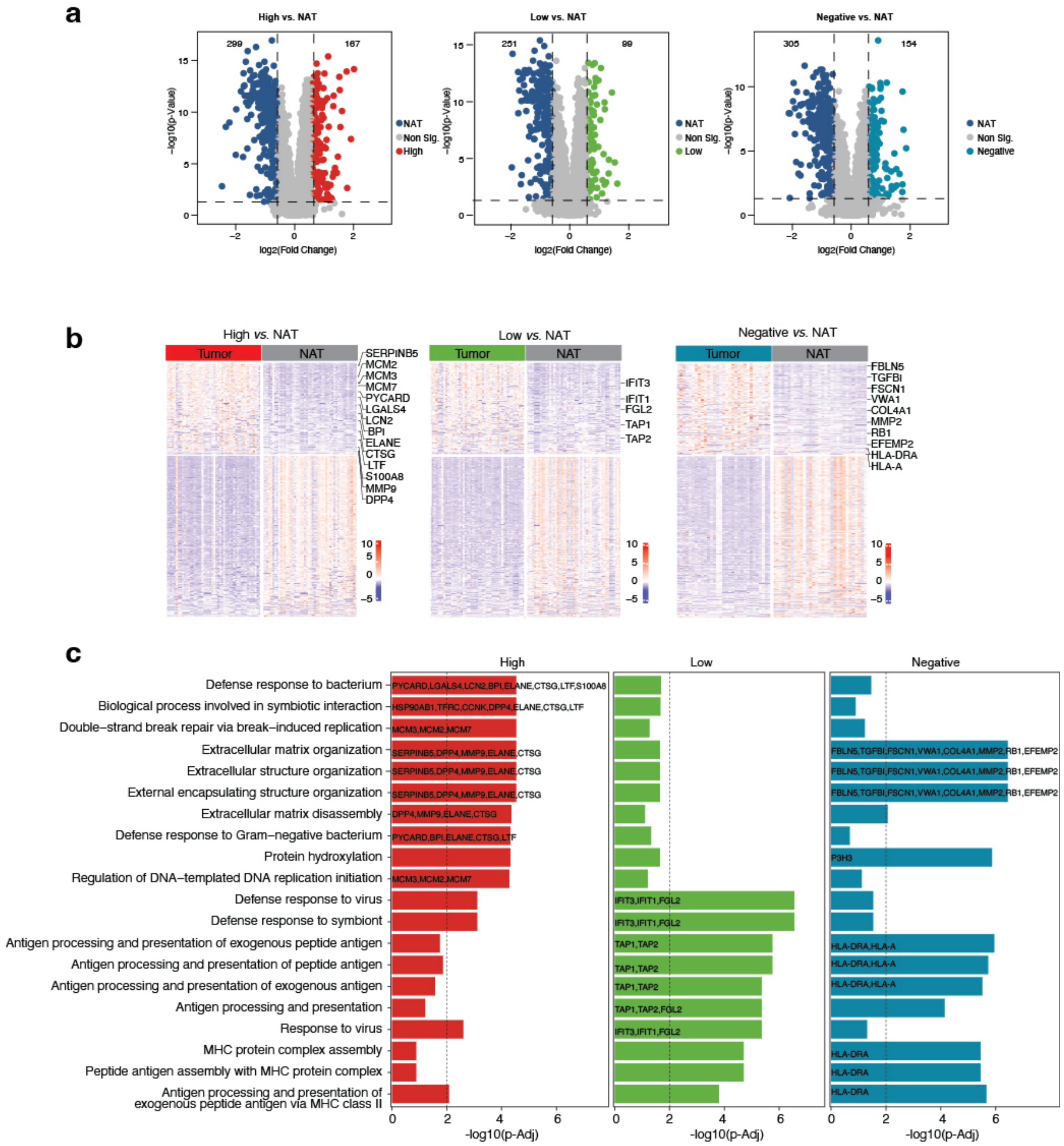


**Supplementary Figure 7: Identification of differential protein expression in three HER2 groups of GCA.**

**a.** Identification of differential protein expression by comparing tumors with normal tissue adjacent to the tumor (NAT) in each HER2 group.

**b.** Differential RNA expression from genes of major histocompatibility complex (MHC) class I and II in different HER2 groups by comparing tumors with corresponding NATs.

**c.** Top 15 enriched Gene oncology (GO) term of differential RNA expression comparing tumors with NATs in different HER2 groups.

**a**

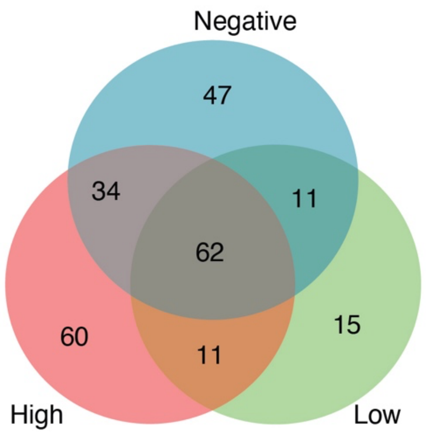


**b**

| High | | Low | Negative | | High & Low | High & Negative | Low & Negative | High & Low & Negative | |
|---|---|---|---|---|---|---|---|---|---|
| HSP90AB1 | LAMC2 | IFIT3 | ARPC1B | ARHGDIB | DDX21 | RCC2 | FEN1 | IKBIP | MCM4 |
| SSB | SMPDL3B | IFIT1 | CNN2 | MFGE8 | XPO5 | FKBP9 | ITGB2 | AGMAT | VCAN |
| GGCT | HSP90AB2P | TAP1 | FBLN5 | FILIP1L | FAP | PLOD3 | SERPINB9 | IGFBP7 | P3H1 |
| PON2 | KPNA2 | TAP2 | PLBD2 | FBN1 | CDK1 | PLOD1 | LXN | NNMT | ISG15 |
| MCM3 | ACSL5 | FGL2 | TGFBI | CBFB | MX1 | CRTAP | LIPA | STAT1 | PLBD1 |
| CAPRIN1 | CEACAM6 | FCGR2A | LDHA | MMP2 | NXT1 | NCL | NPL | EFEMP1 | IGF2BP3 |
| NOP2 | SERPINB8 | H1-5 | IL4I1 | POLD2 | TMEM164 | STMN1 | TAPBP | LTBP2 | FTL |
| RALY | DPP4 | DCK | FSCN1 | SAAL1 | RETN | CTSB | LCP1 | FHL3 | FN1 |
| PPA1 | RRM2 | SQSTM1 | ITGAM | HAT1 | MPO | SET | TNC | FKBP10 | THBS2 |
| MCM2 | NEU1 | NIP7 | CNN3 | CORO1A | AZU1 | CPNE1 | DOK2 | CTHRC1 | CDH17 |
| ERBB2 | ISLR | USP11 | GUSB | EMILIN2 | MMP8 | LPCAT1 | HLA-DPB1 | SERPINH1 | COL12A1 |
| HSPH1 | LCN2 | PRTN3 | KIF3B | C1QC | | CTSA | | GBP1 | CEACAM5 |
| DDX27 | MMP9 | SLCO1B7 | CTSC | NOP14 | | SLC16A3 | | ITGAX | PLSCR1 |
| PYCARD | PIGR | LPXN | VWA1 | C20orf27 | | MXRA5 | | TPM4 | PLOD2 |
| LACTB2 | BPI | FAM117A | MFAP2 | HTRA1 | | DCTPP1 | | EPCAM | SLC2A3 |
| CBX3 | ELANE | | SUMO1 | EVL | | MMP14 | | TYMP | TOP2A |
| GAR1 | CTSG | | AIF1 | FASTKD5 | | CTSS | | VIL1 | HLA-DMB |
| HSD17B11 | MUC5B | | HLA-DRA | RB1 | | GDA | | MUC13 | FCER1G |
| TFRC | PDAP1 | | NASP | EFEMP2 | | HK3 | | PCNA | SASH1 |
| S100A11 | LTF | | HLA-A | P3H3 | | PPIC | | LTBP1 | GPRC5A |
| RSL1D1 | SDCBP2 | | CACYBP | WDR3 | | ASS1 | | CYBB | CAMP |
| S100A6 | PLPP2 | | COL4A1 | SERPINE2 | | GRN | | CTSZ | THBS3 |
| ANXA4 | NCAPH | | APOE | MAP1LC3B | | PUS7 | | CPVL | OLFM4 |
| SMC2 | S100A8 | | DPYSL3 | | | L1RE1 | | SPARC | THBS4 |
| LGALS3BP | FUT6 | | | | | GPNMB | | ANP32E | STMN2 |
| SERPINB5 | MIEN1 | | | | | BRIX1 | | CD55 | SFRP4 |
| LGALS4 | SPARCL1 | | | | | MISP | | GGH | DMBT1 |
| CCNK | GRB7 | | | | | AOC1 | | MNDA | DEFA3 |
| EPS8L3 | ANXA13 | | | | | TMSB10 | | TIMP1 | S100A12 |
| MAGED2 | NEURL4 | | | | | ITPRID2 | | CYBA | SPINT2 |
| | | | | | | FABP1 | | MCM6 | CPS1 |
| | | | | | | ANPEP | | | |
| | | | | | | GCNT3 | | | |
| | | | | | | HLA-DRB3 | | | |

**Supplementary Figure 8: Number (a) and names (b) of proteins specific to HER2 groups and those common among them.**

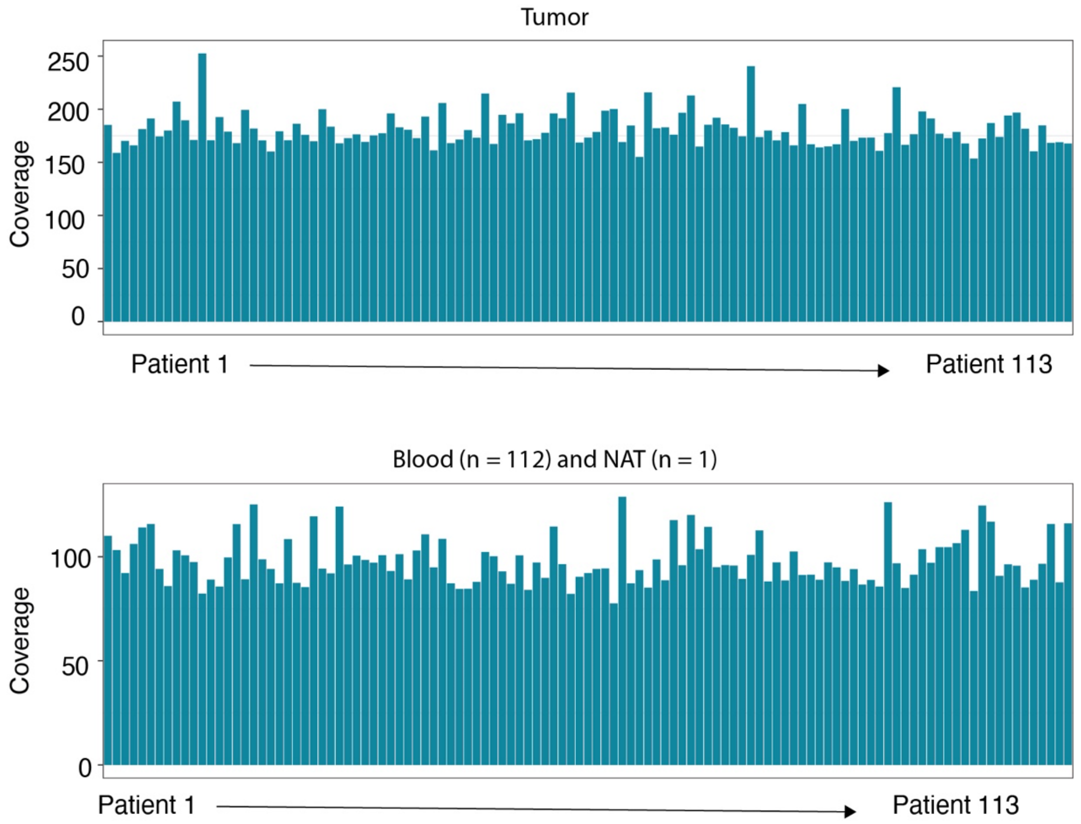


**Supplementary Figure 9: Quantification and comparison of sequencing coverage of whole exon sequencing from GCA cohort.**

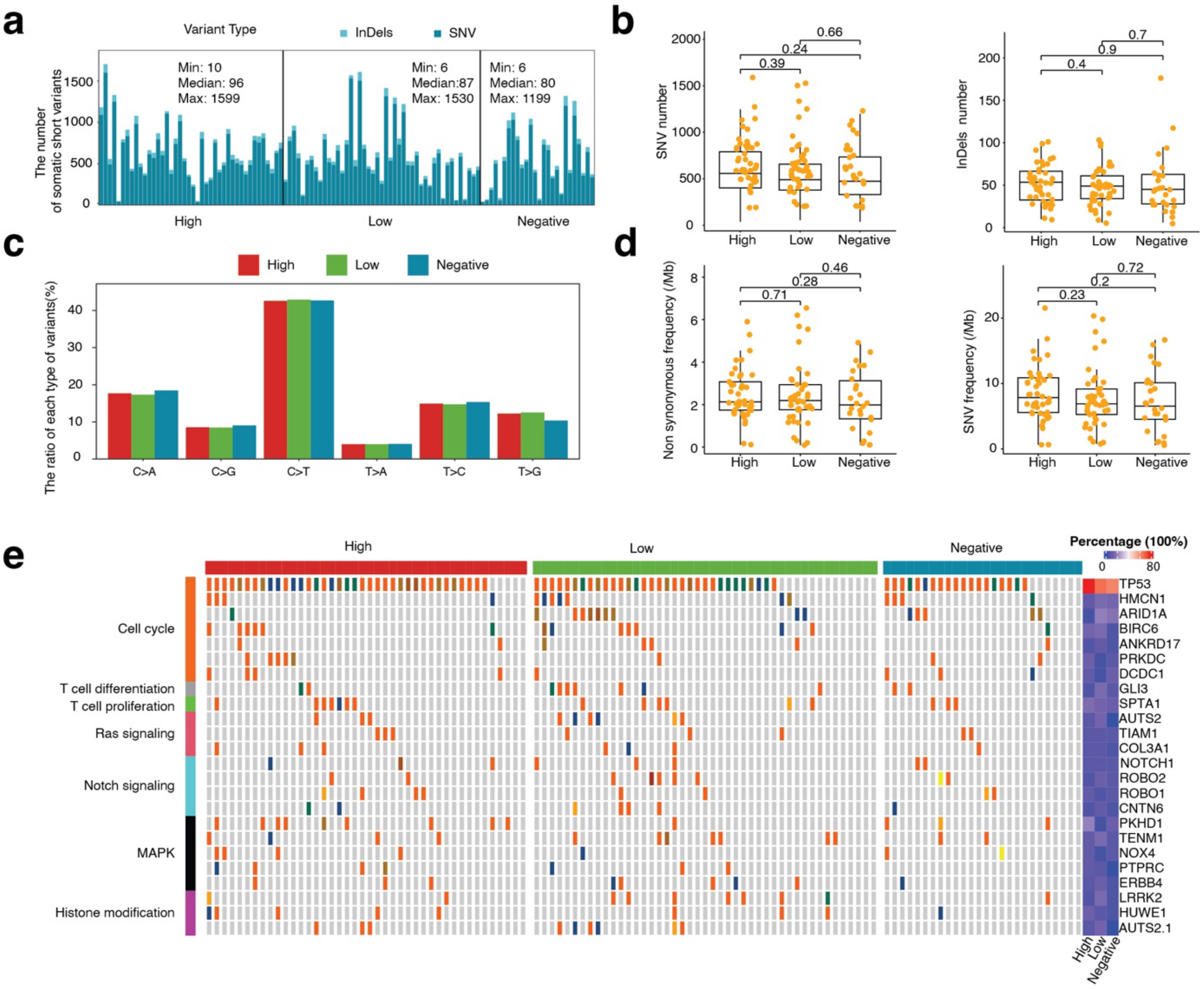


**Supplementary Figure 10: comparison of somatic mutations in different HER2 groups.**

- **a.** Comparison of somatic short variant types (single-nucleotide variant (SNV) and insertions and deletions (InDels)) in three HER2 groups.
- **b.** Quantification comparison of SNV number (left panel) and InDels numbers (right panel) in three HER2 groups.
- **c.** Comparison of mutation types in three HER2 groups.
- **d.** Frequency comparison of non-synonymous and SNV in three HER2 group.
- **e.** Ranking plots of the top mutated oncogenes and tumor suppressor genes involved in key signal pathways.

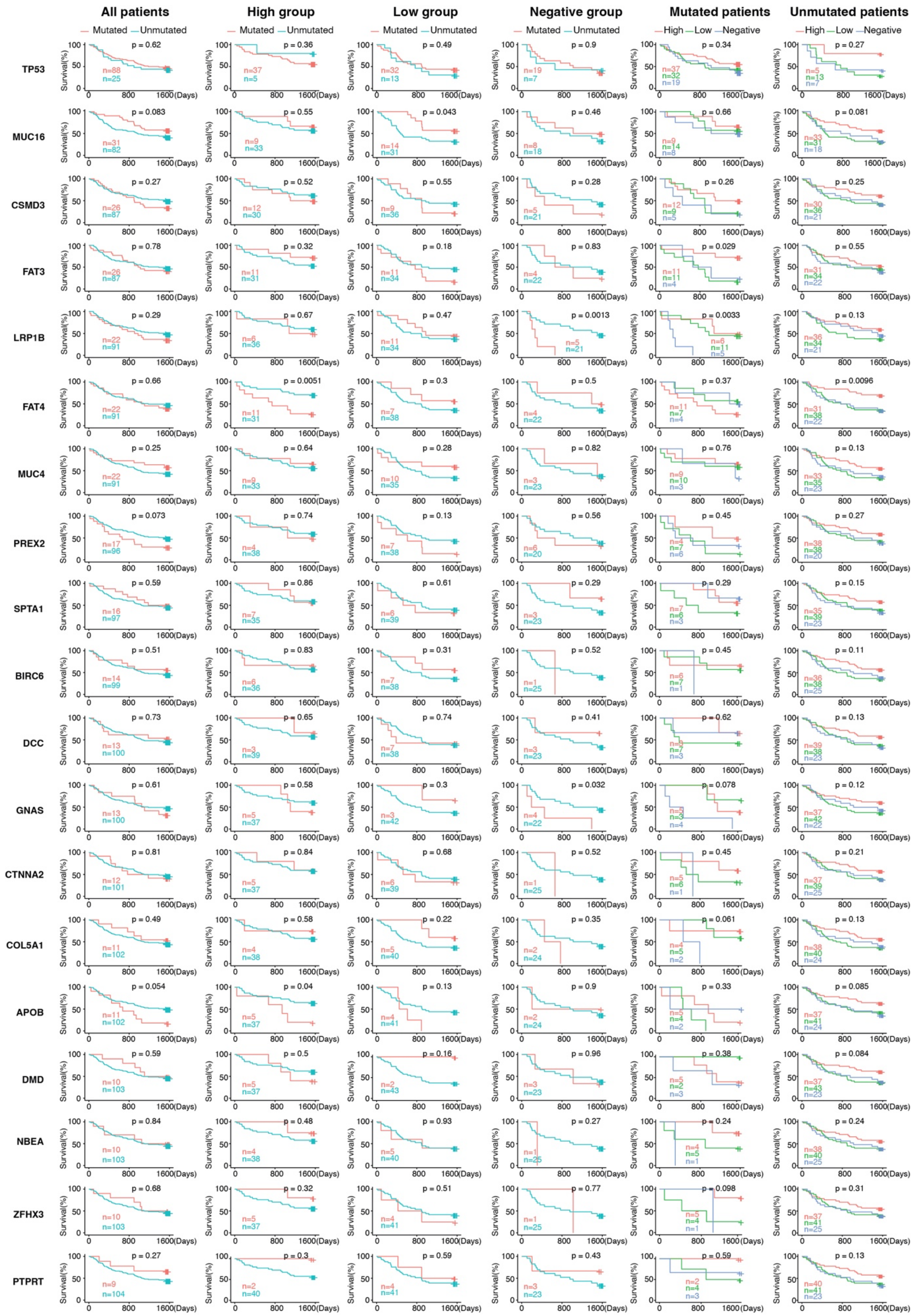


**Supplementary Figure 11: Relationship between the prognosis of GCA patients, top mutated genes, and HER2 grouping.**

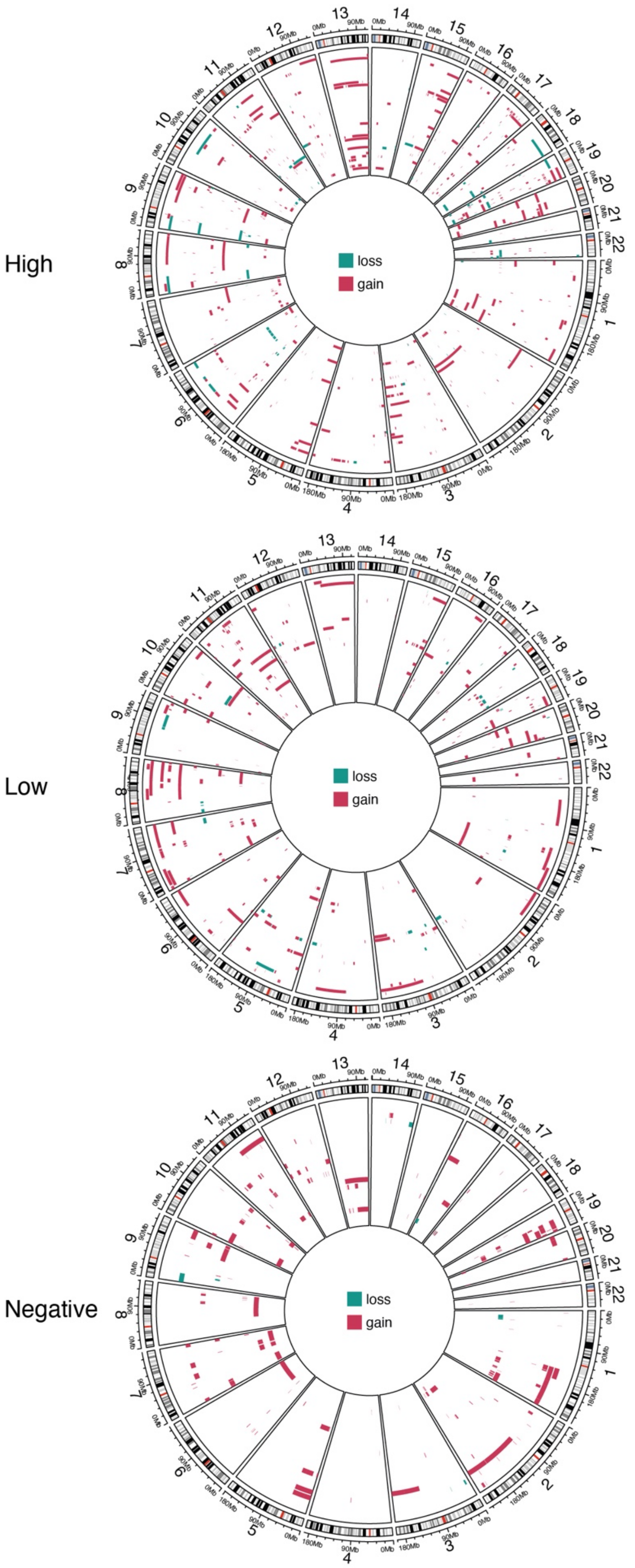


**Supplementary Figure 12: Copy number alterations in different HER2 groups.**

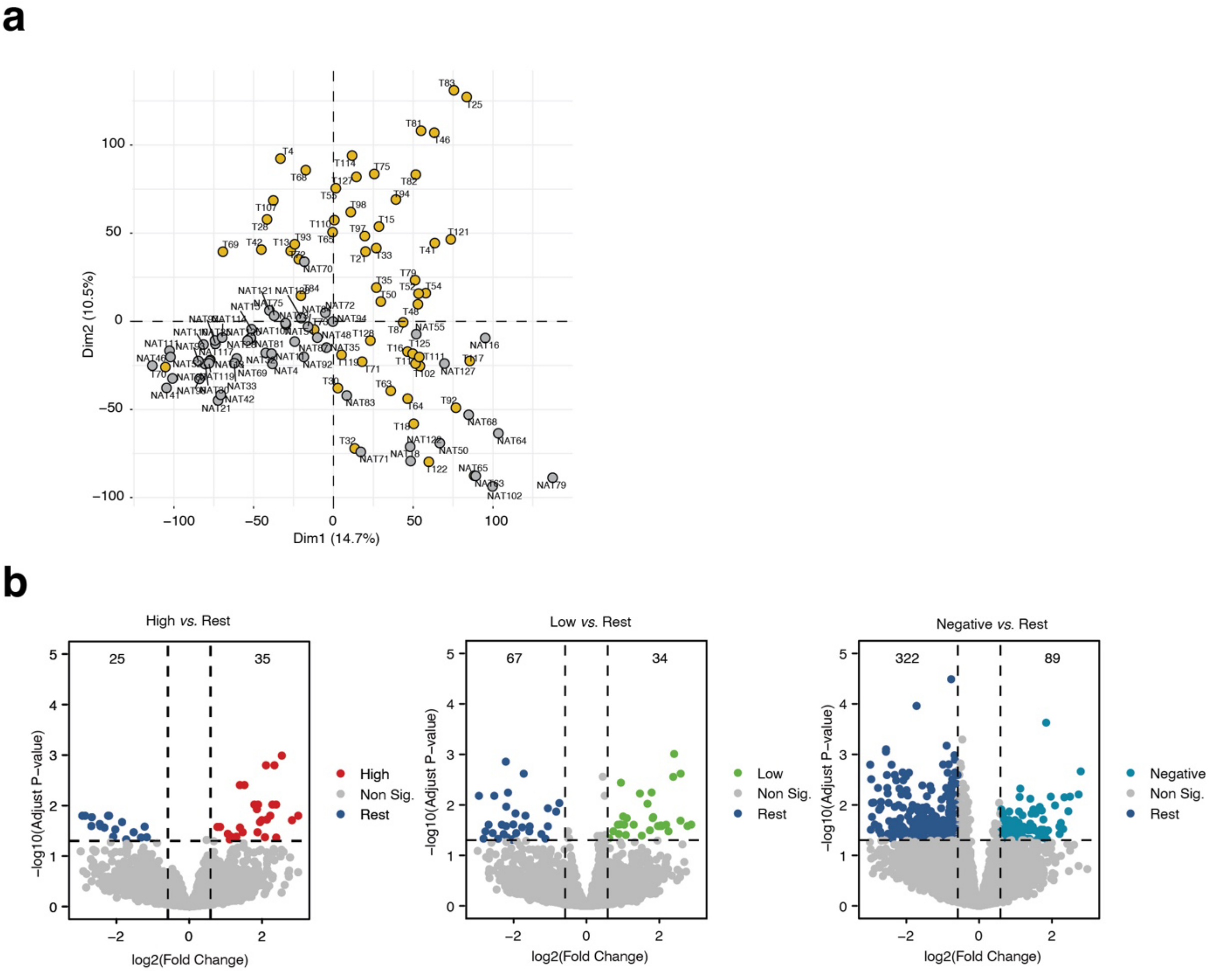


**Supplementary Figure 13: RNA expression signature in each HER2 groups.**

- **a.** Principal component analysis (PCA) of RNA expression in tumors and NAT.
- **b.** Volcano plots show the identification of differential RNA expression by comparing of each HER2 group with rest of groups.

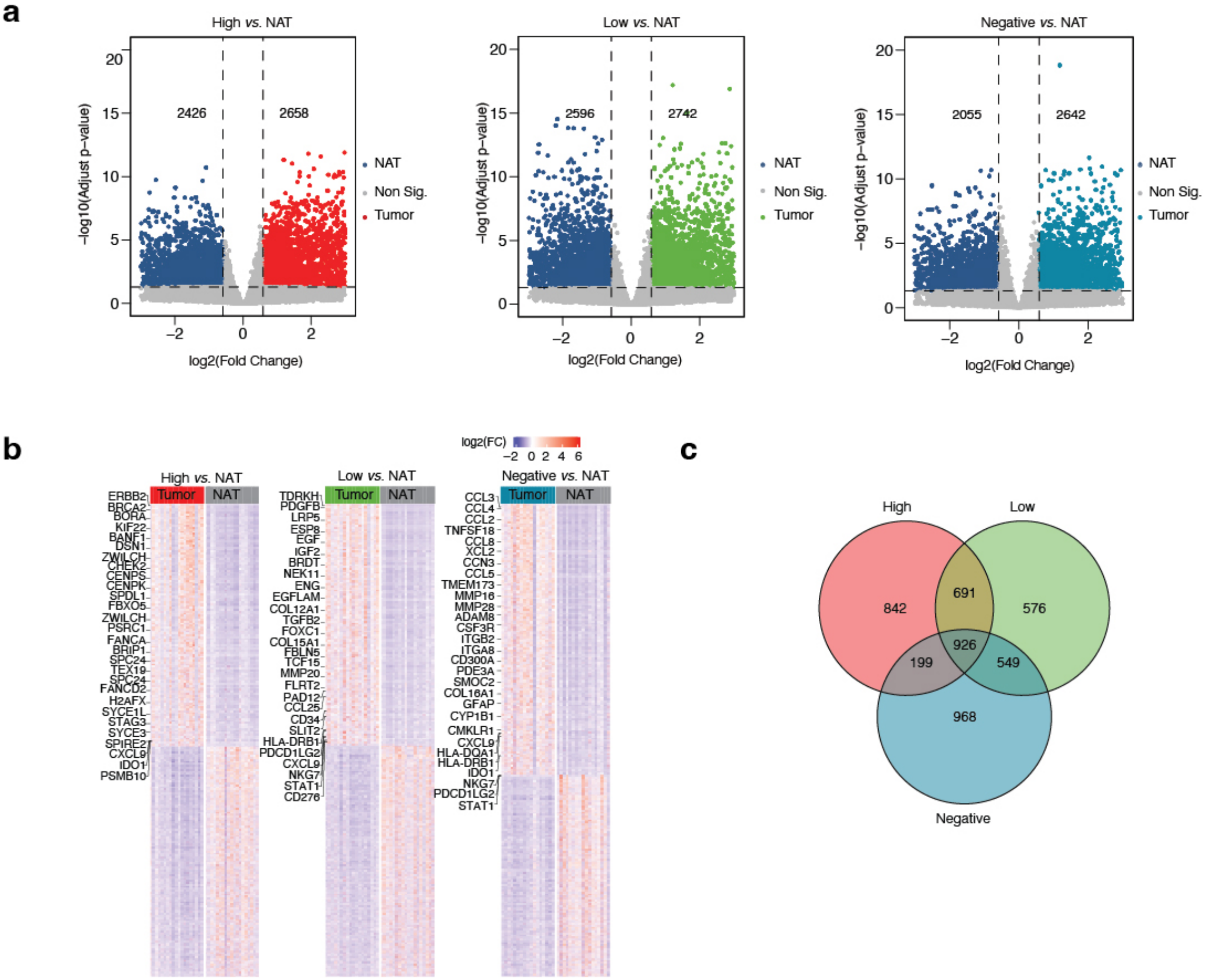


**Supplementary Figure 14: Identification of differential RNA expression by comparing tumor with its corresponding NAT in each HER2 group.**

- **a.** Volcano plots show the identification of differential RNA expression by comparing of tumor with its corresponding NAT.
- **b.** Left: Differential RNA expression in HER2-high group by comparing tumors with corresponding NATs.
- **c.** Middle: Differential RNA expression in HER2-low group by comparing tumors with corresponding NATs.
- **d.** Right: Differential RNA expression in HER2-negative group by comparing tumors with corresponding NATs.
- **e.** Venn diagram shows the unique and common differential RNA expression in each HER2 group.

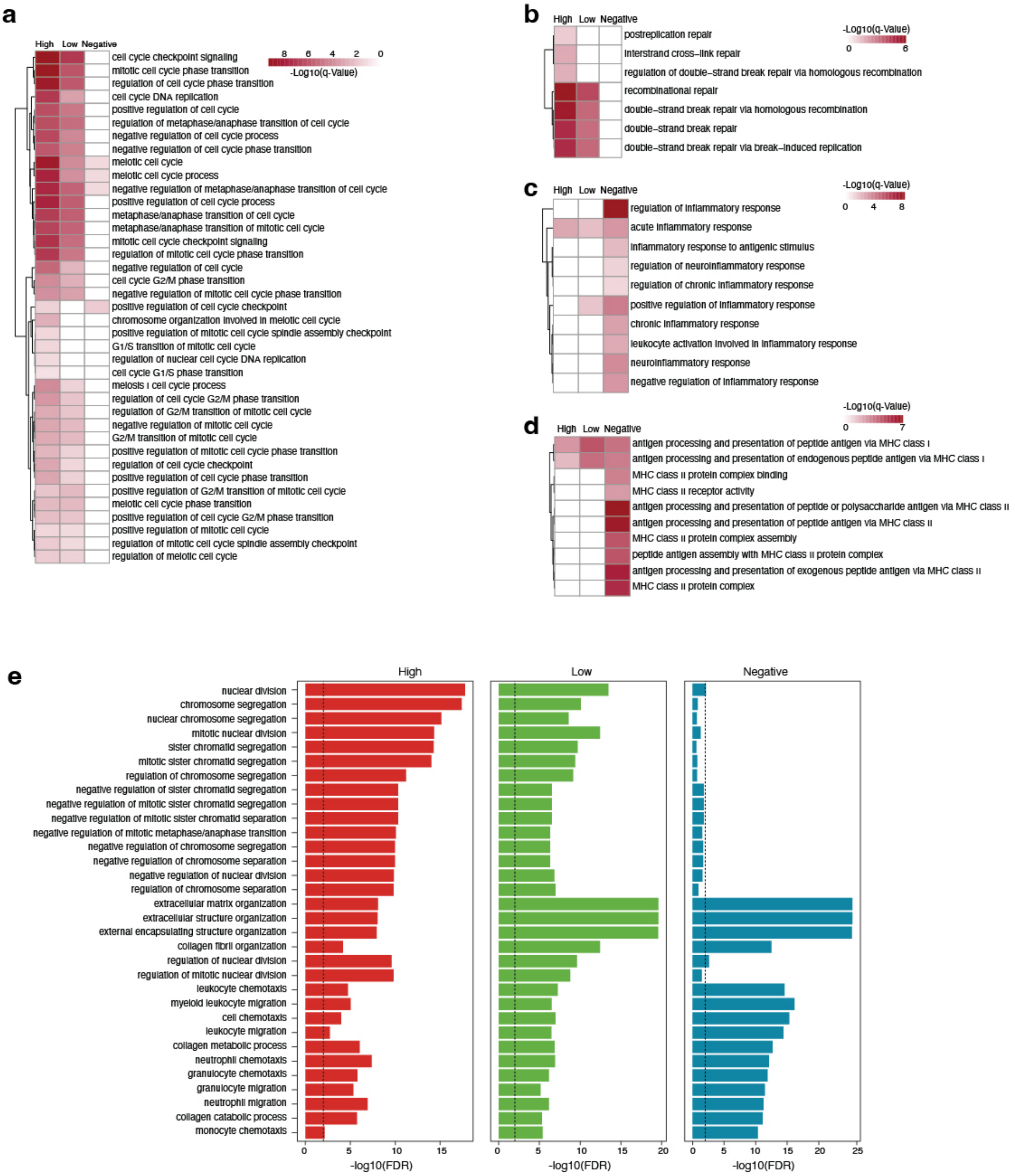


**Supplementary Figure 15: Enrichment gene ontology (GO) term comparison in different HER2 groups.**

- **a.** GO terms from cell cycle relevant terms.
- **b.** GO terms from DNA repair relevant terms.
- **c.** GO terms from inflammation relevant terms.
- **d.** GO terms from major histocompatibility complex (MHC) relevant terms.
- **e.** Top 15 enriched Gene oncology (GO) term of differential RNA expression comparing tumors with NATs in different HER2 groups.

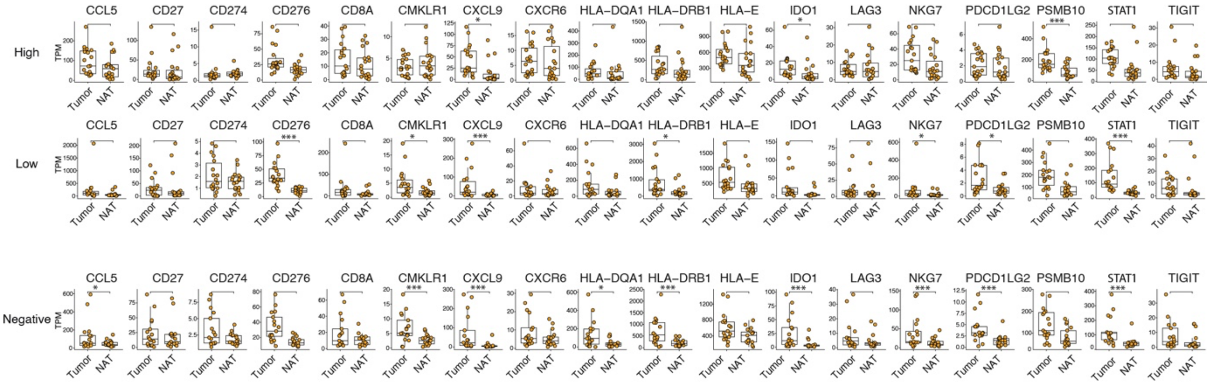


**Supplementary Figure 16: Comparison of RNA expression from 18 immune genes between tumors and NAT in different HER2 groups.**

* = False discovery rate (FDR) < 0.05; ** = FDR < 0.01; *** = FDR < 0.001; TPM = Transcripts Per Million.

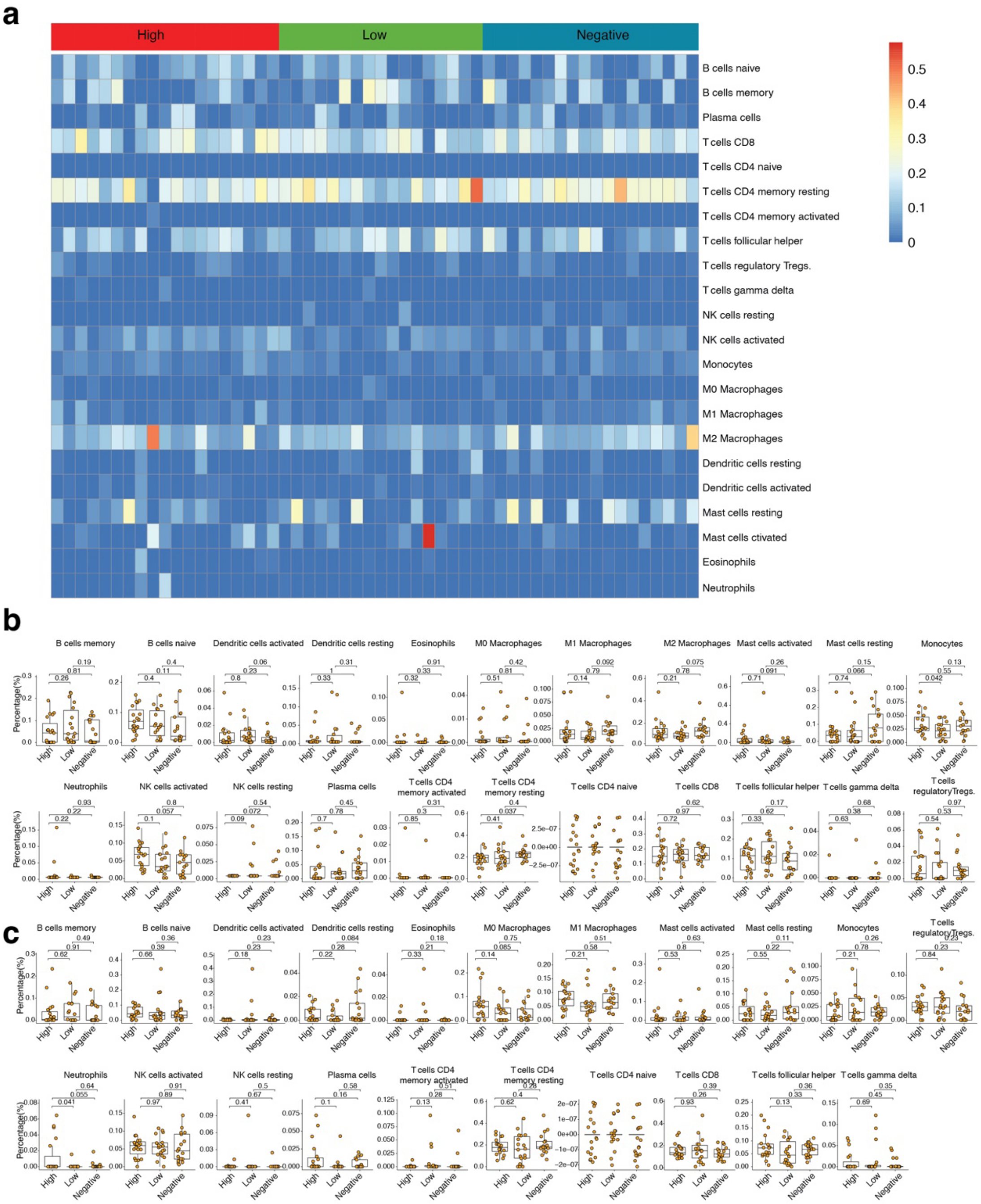


**Supplementary Figure 17: Deciphering tumor microenvironment in different HER2 group.**

- **a.** Quantitative comparison of different cell types in tumors only from different HER2 groups.
- **b.** Cell component comparison from bulk RNA-seq predictions of NAT in different HER2 groups.
- **c.** Quantitative comparison of different cell types in NAT from different HER2 groups.

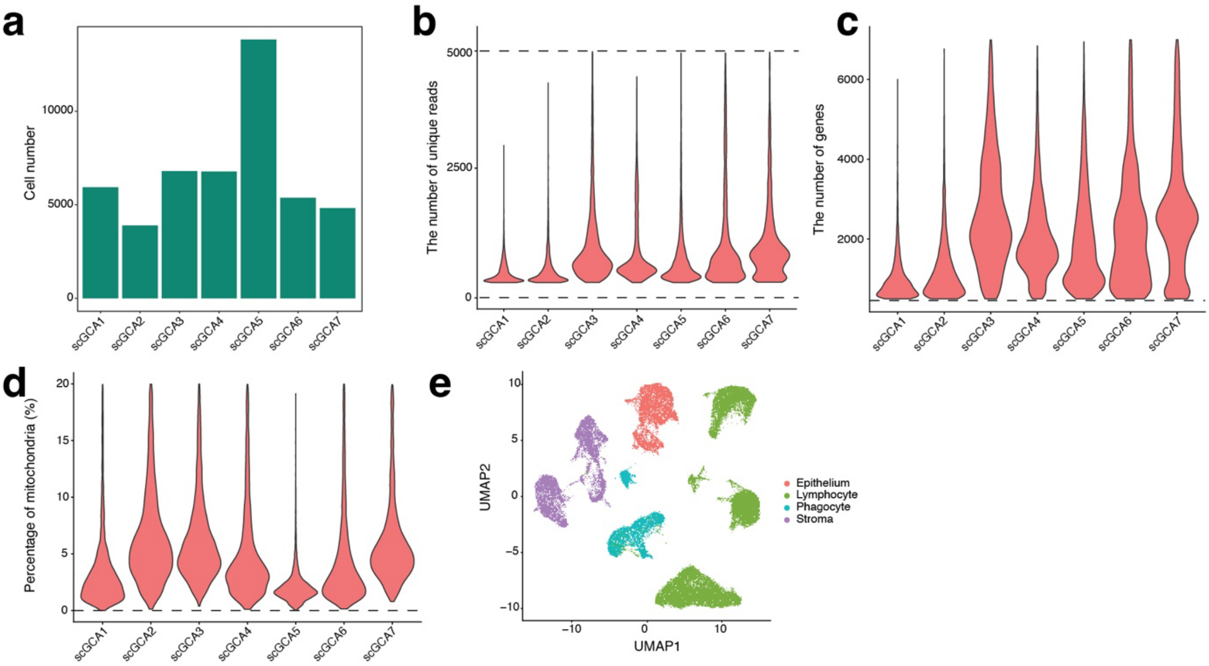


**Supplementary Figure 18: Single-cell RNA-Seq (scRNA-Seq) analysis from seven GCA patients.**

- **a.** Cell numbers identified in each GCA patient from scRNA-Seq.
- **b.** The number of unique reads identified in each GCA patient from scRNA-Seq, with dotted lines indicating the cutoff.
- **c.** The number of genes identified in each GCA patient from scRNA-Seq.
- **d.** The percentage of mitochondria identified in each GCA patient from scRNA-Seq.
- **e.** Cell components identified from scRNA-Seq in seven GCA patients.

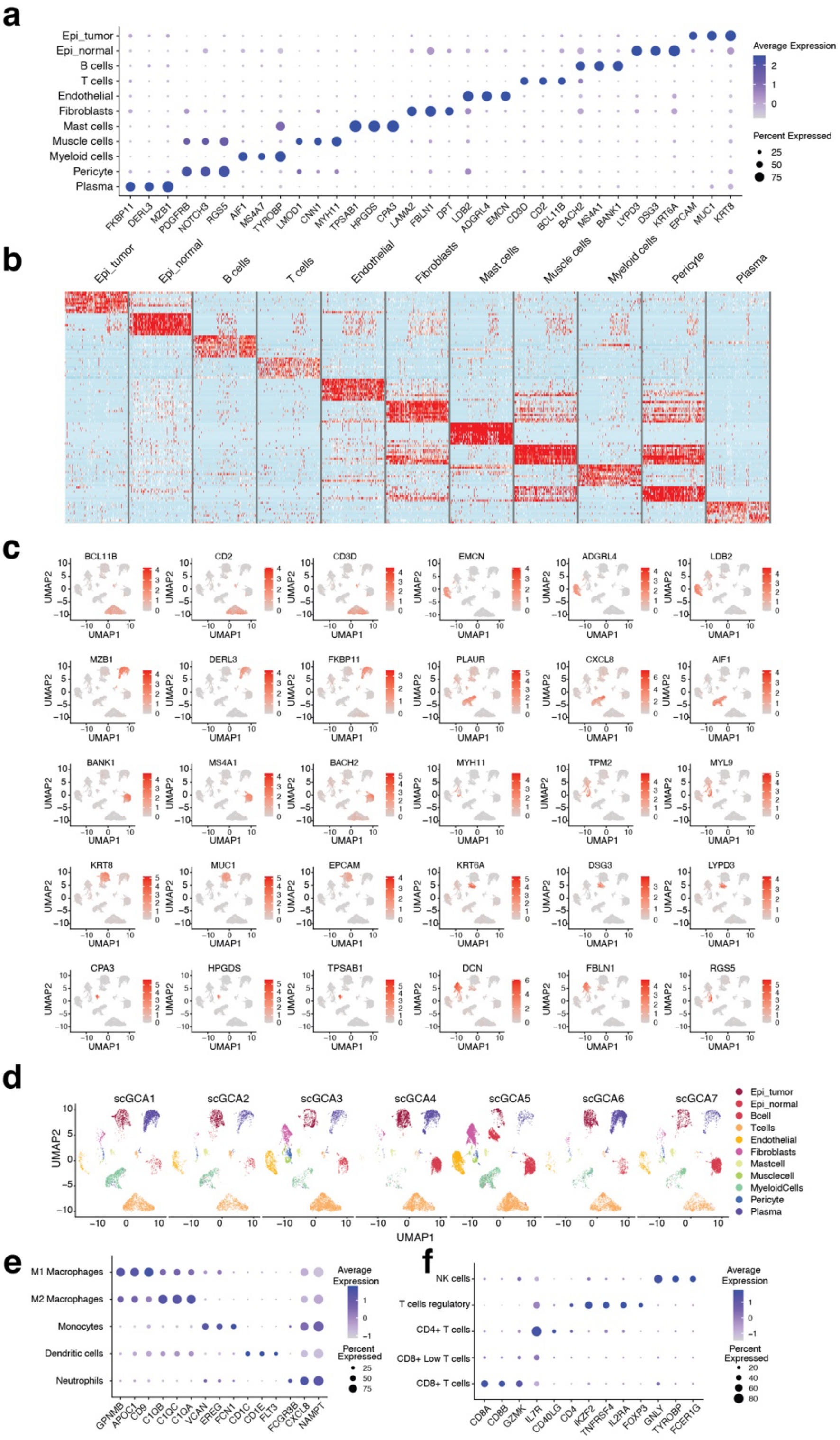


**Supplementary Figure 19: Single-cell RNA-Seq (scRNA-Seq) analysis of GCA patients.**

- **a.** Bubble plots present the top three marker genes for each cell type.
- **b.** Heatmaps present the marker genes extracted for each cell type from scRNA-Seq.
- **c.** The top marker genes for each cell type are projected on the UMAP.
- **d.** Distribution of each cell type in each GCA patient.
- **e.** Bubble plots present the top three marker genes for subtypes of myeloid cells.
- **f.** Bubble plots present the top three marker genes for subtypes of lymphoid cells.

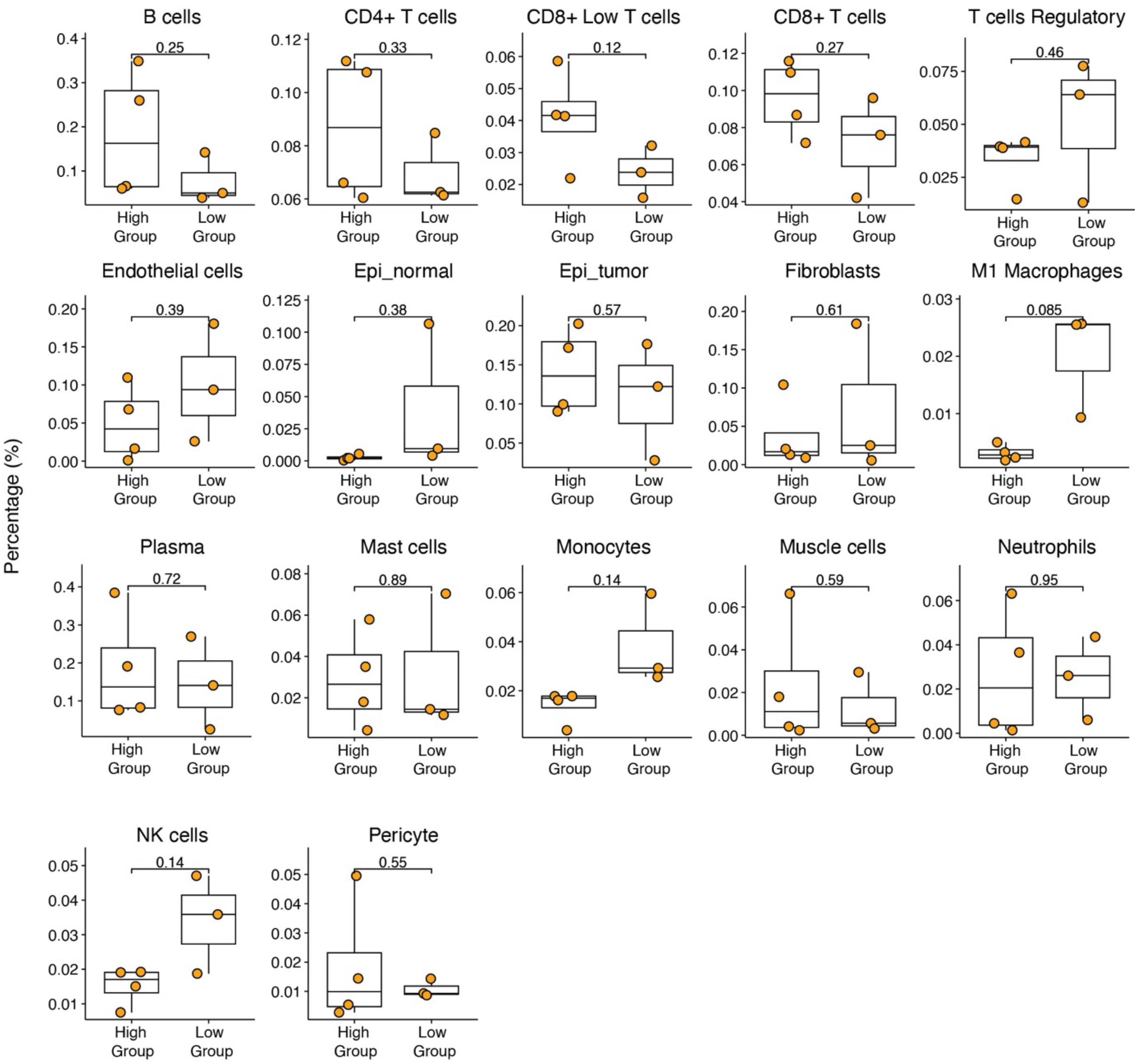


**Supplementary Figure 20: Comparison of cell components between the HER2 high and low groups in GCA patients.**

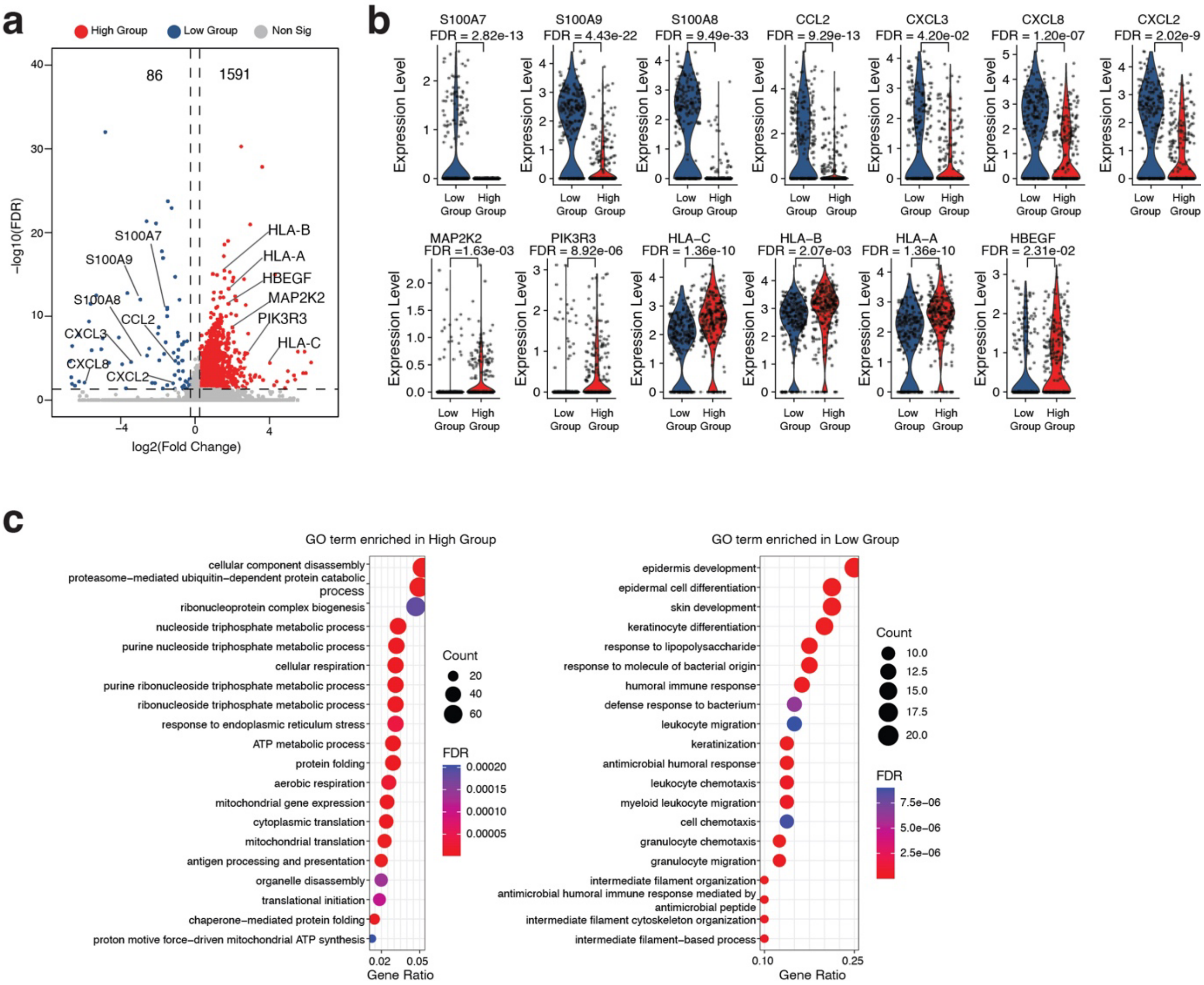


**Supplementary Figure 21: Comparison of gene signatures of M2 macrophages between the HER2 high and low groups.**

- **a.** Volcano plots present the differential genes identified from M2 macrophages in the HER2 high and low groups from scRNA-Seq, with some feature genes highlighted.
- **b.** Violin plots present the comparison of feature gene expression from M2 macrophages in the HER2 high and low groups from scRNA-Seq.
- **c.** Top 20 GO (Gene Ontology) terms enriched from the identified differential genes in M2 macrophages in the HER2 high and low groups from scRNA-Seq.

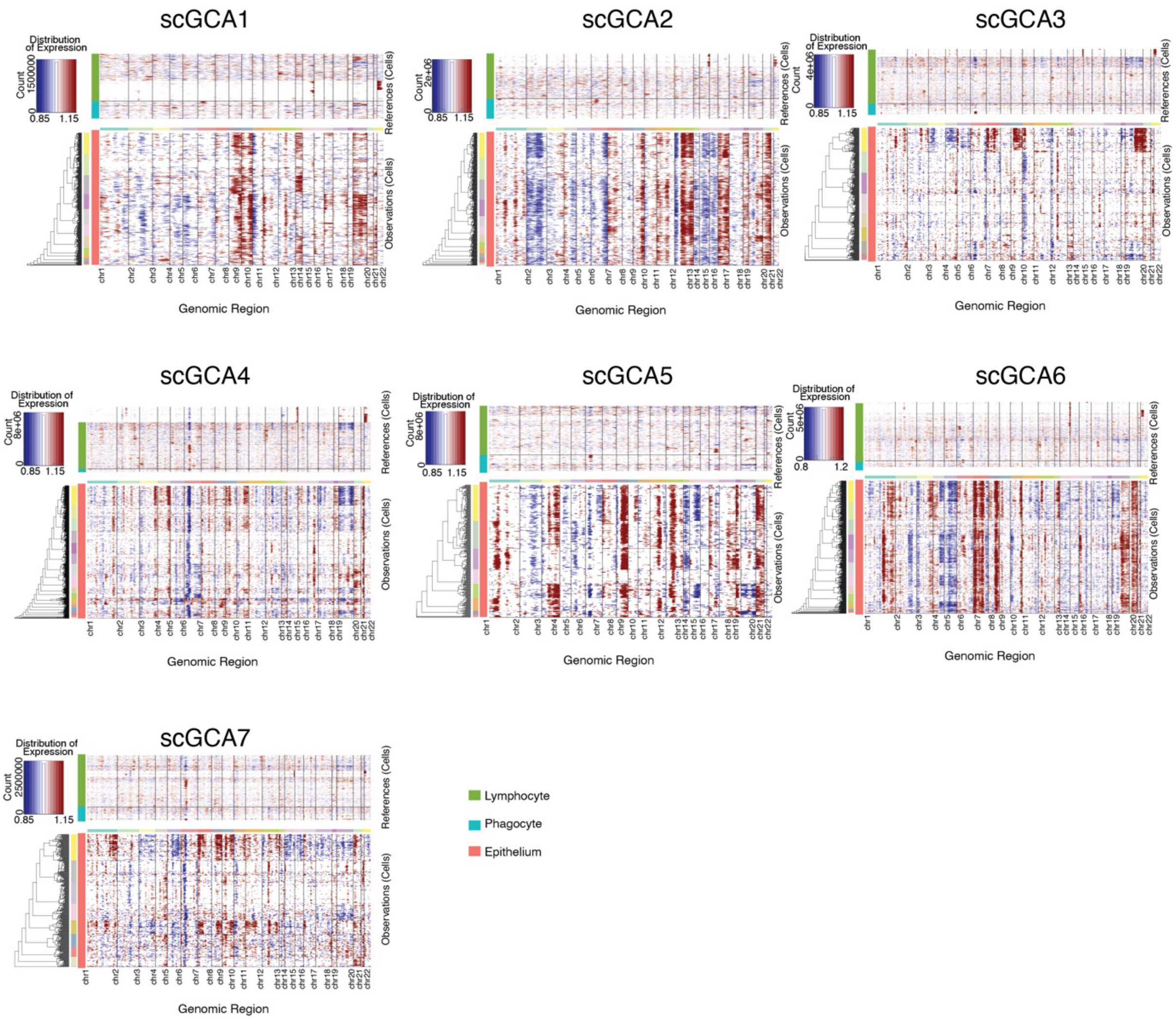


**Supplementary Figure 22: Heatmaps present the predicted copy number variations (CNV) for epithelial tumor cells from scRNA-Seq for each GCA patient.** The CNVs were calculated using lymphocytes and phagocytes as the background for each patient.

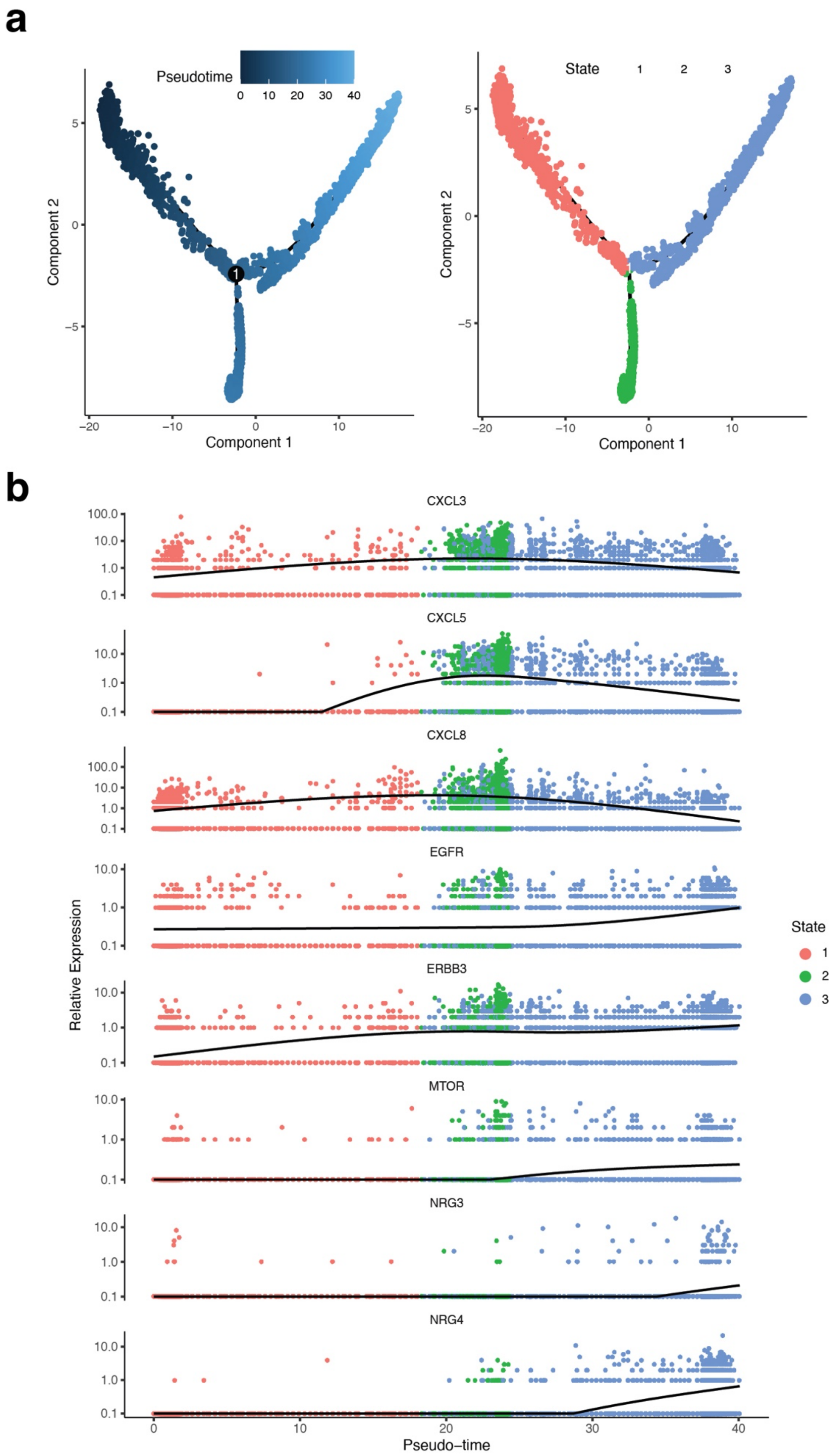


**Supplementary Figure 23: Pseudotime projection of epithelial cell fate from single-cell RNA-Seq (scRNA-Seq) data of GCA patients (a) and feature marker gene expression in different cell states (b).**

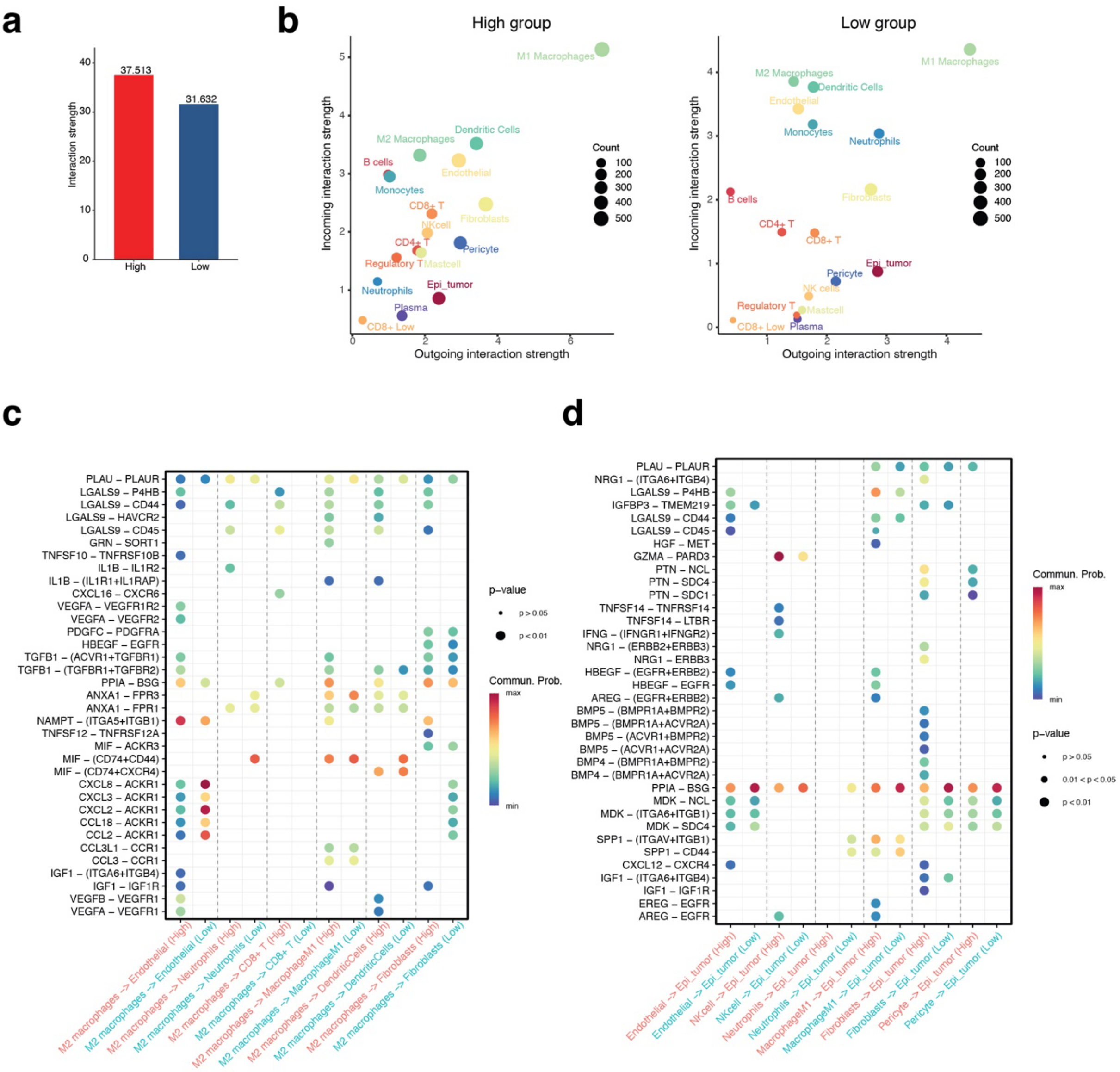


**Supplementary Figure 24: Cell-to-cell communication among different cell types from single-cell RNA-Seq in different groups of GCA patients.**

- **a.** Comparison of total interaction strength in HER2 high and low groups of GCA patients.
- **b.** Scatter plots present the incoming and outgoing interaction strength for each cell type identified from single-cell RNA-Seq in HER2 high and low groups of GCA patients.
- **c.** Bubble plots present the comparison of signals sent by M2 macrophages to endothelial cells, neutrophils, CD8+ T cells, dendritic cells, and fibroblasts in high and low groups.
- **d.** Bubble plots present the comparison of signals received by epithelial tumor cells from endothelial cells, NK cells, neutrophils, M1 macrophages, fibroblasts, and pericytes in high and low groups.

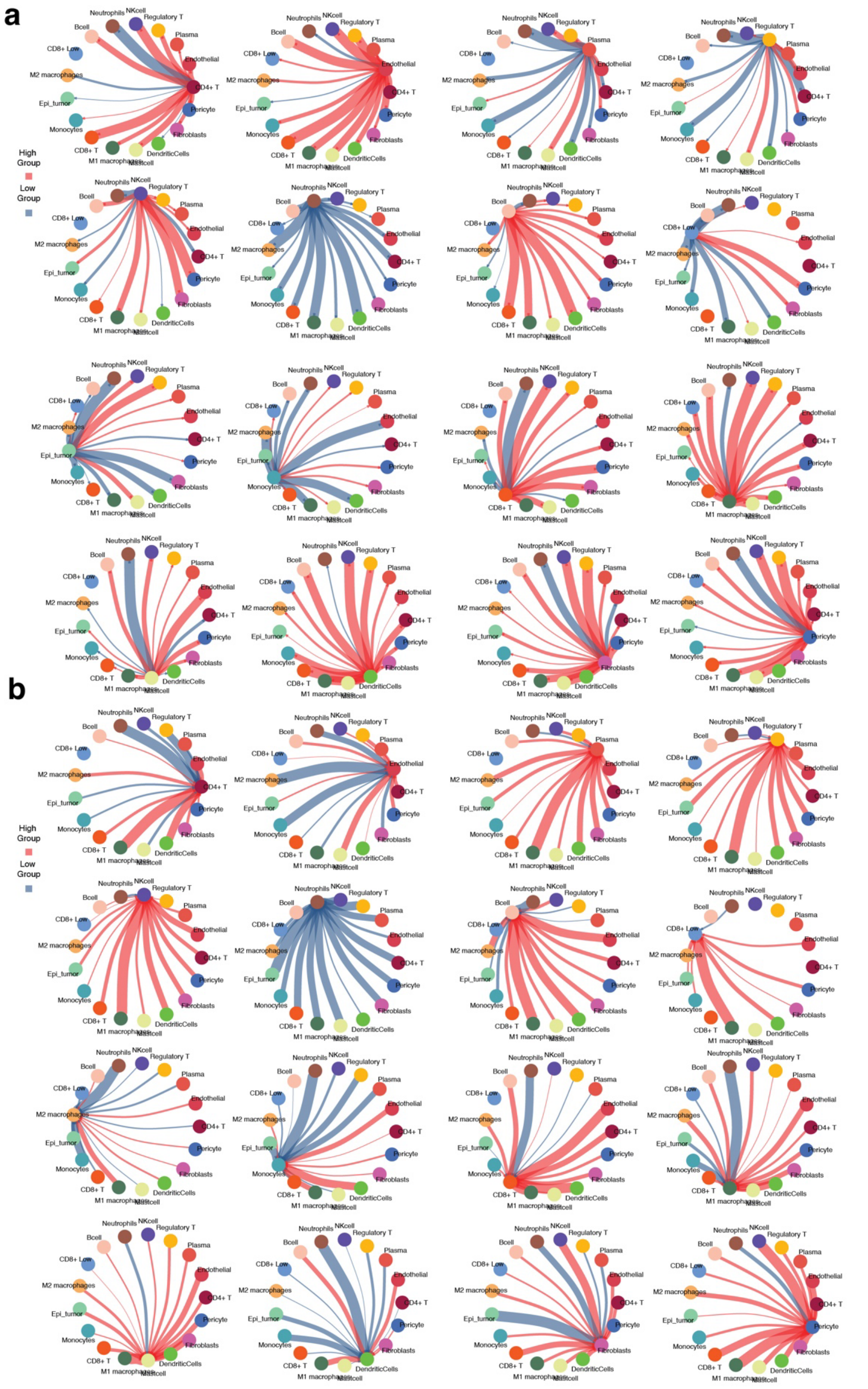


**Supplementary Figure 25: Signal communication among different cell types from single-cell RNA-Seq in GCA patients.**

- **a.** Circle plots present the signals sent by each cell type in high and low groups.
- **b.** Circle plots present the signals received by each cell type in high and low groups.

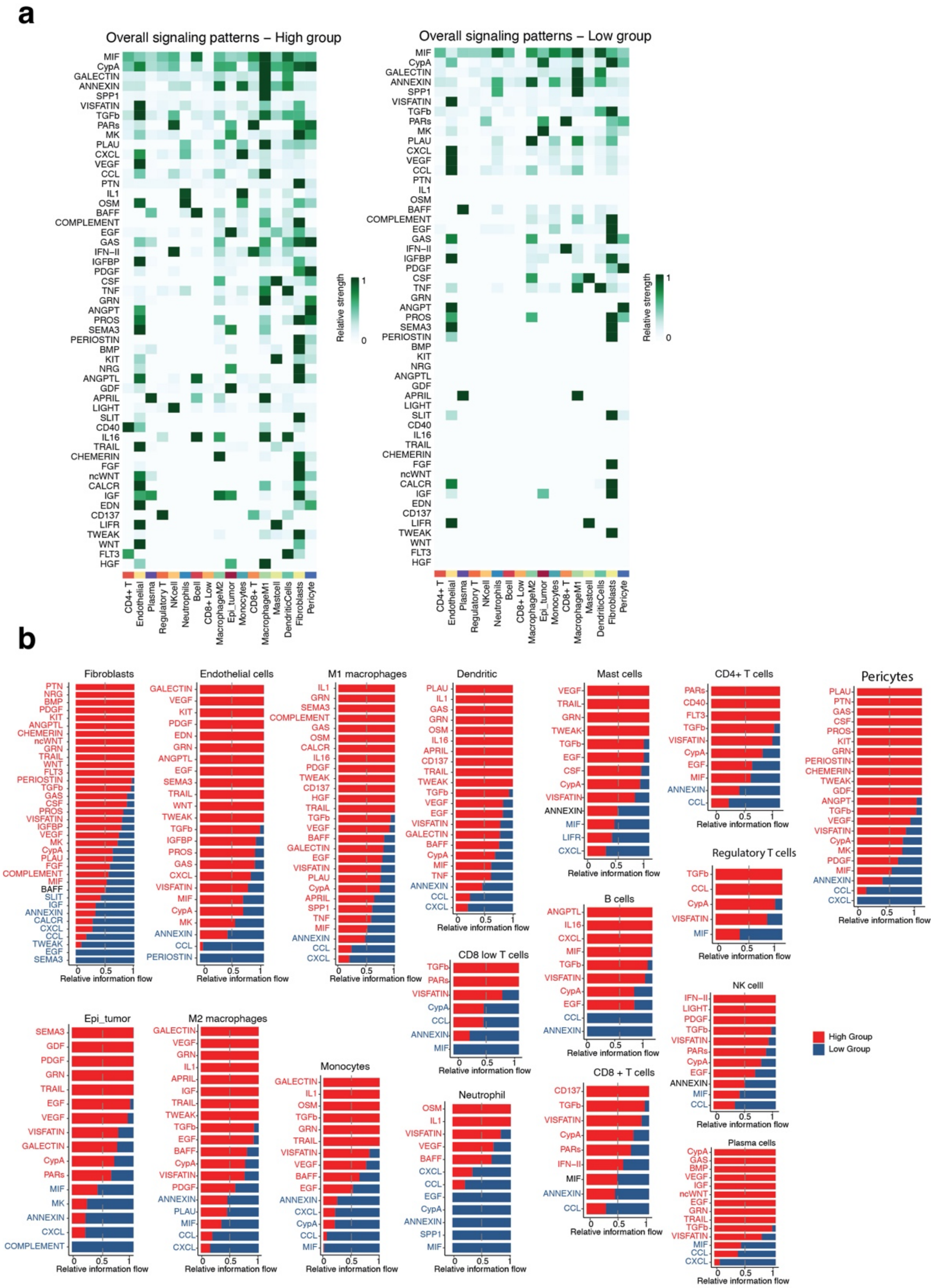


**Supplementary Figure 26: Comparisons of different signal pathway communications in high and low groups.**

**a.** Heatmaps present the total strength of each signal pathway from all cell types.

**b.** Bar plots present the strength of each signal pathway from each cell type by comparing the high and low groups.

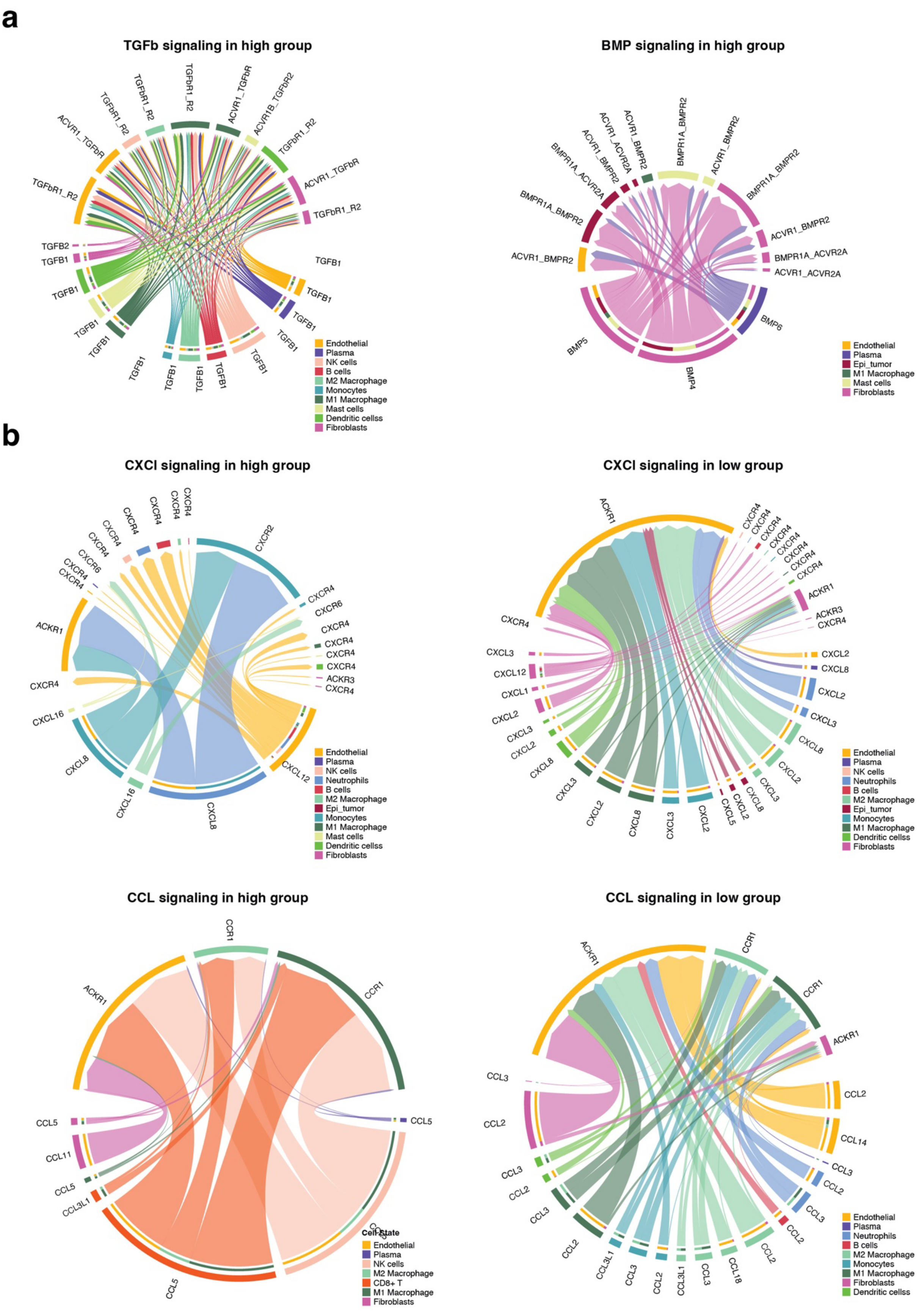


**Supplementary Figure 27: Signal pathway communications among different cell types.**

- **a.** TGF-beta and BMP pathways.
- **b.** CXCL and CCL pathways.

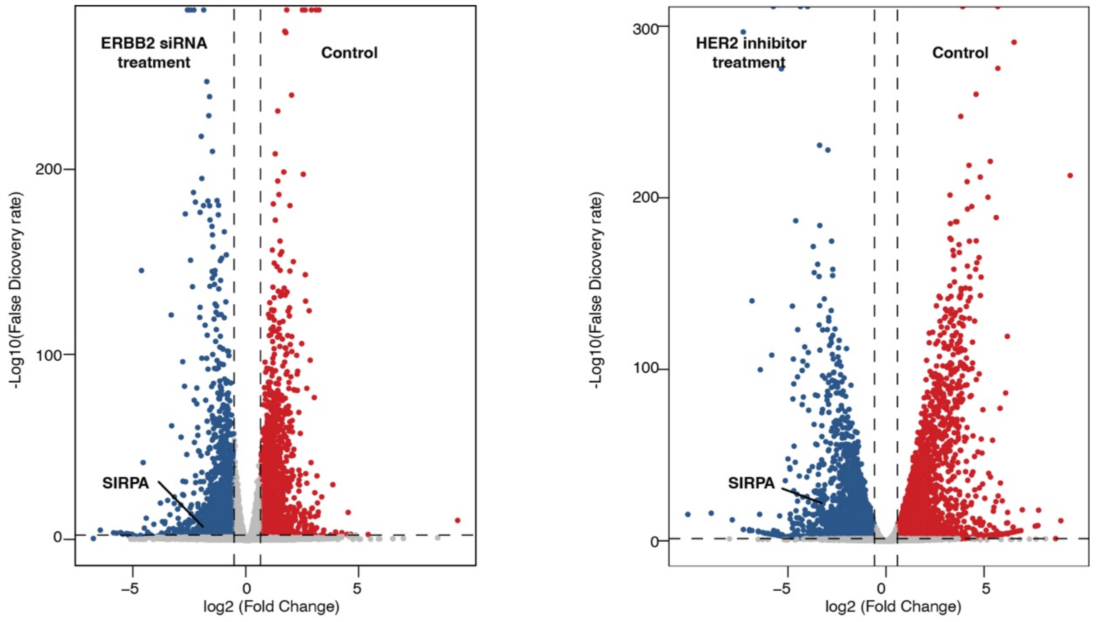


**Supplementary Figure 28: Differential gene expression analysis between control and ERBB2 siRNA treatment (Left), and between control and ERBB2 inhibitor treatment (Right) in OE19 cell line.**

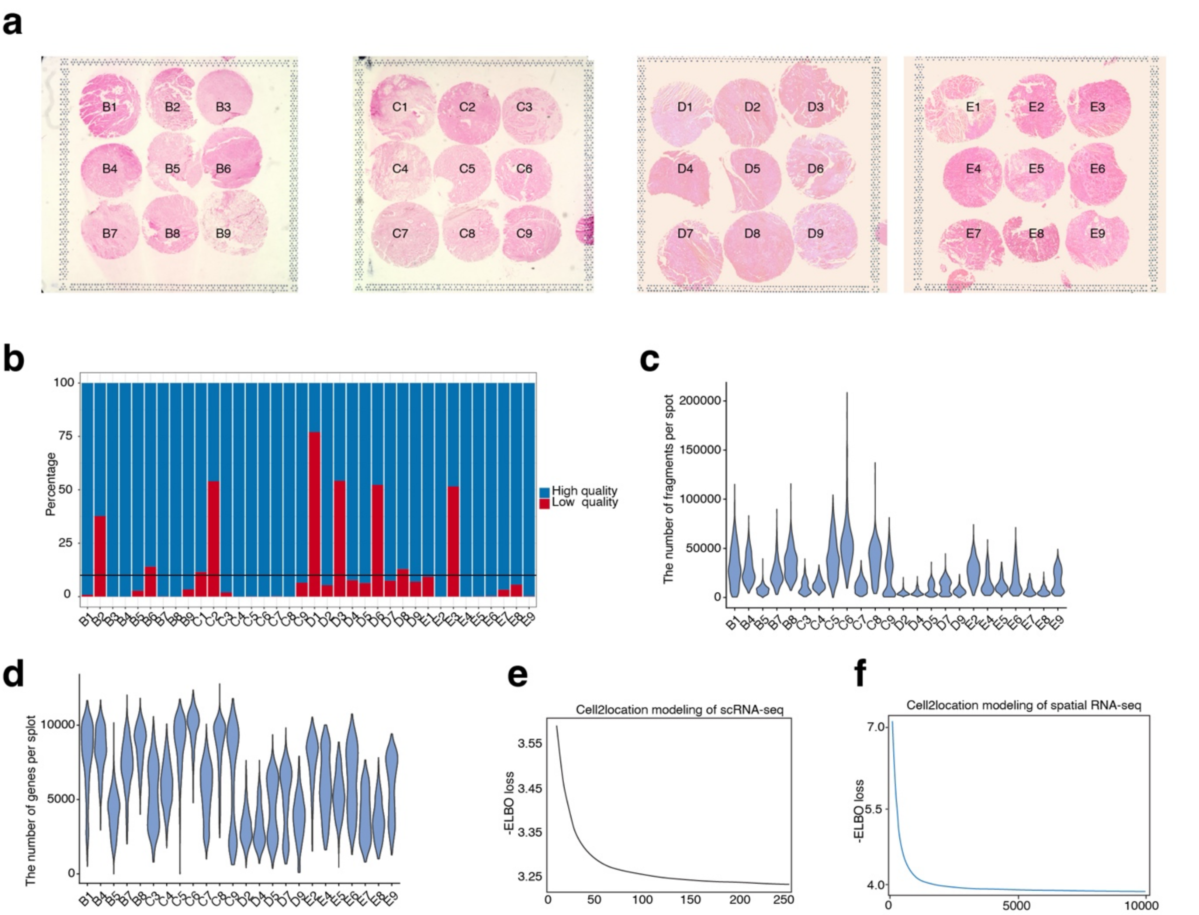


**Supplementary Figure 29: Spatial transcriptomics analysis of GCA patients.**

**a.** H&E staining of tissue microarray punched from boundary regions of tumor and immune cells in different HER2 groups of GCA patients. The HER2 high group includes the following regions: B1, C6, C3, D9, E7, and E8; the HER2 low group includes the following regions: B4, B5, C7, C4, C9, C8, C5, D7, D4, D5, D2, E4, E9, E5, and E6; and the HER2 negative group includes the following regions: B7, B8, and E2.

**b.** The threshold used to remove low-quality regions from spatial transcriptomics.

**c.** The number of fragments per spot in spatial transcriptomics for high-quality regions.

**d.** The number of genes per spot in spatial transcriptomics for high-quality regions.

**e.** Modeling curve of single-cell RNA-Seq using the Cell2location package.

**f.** Modeling curve of spatial transcriptomics using the Cell2location package.

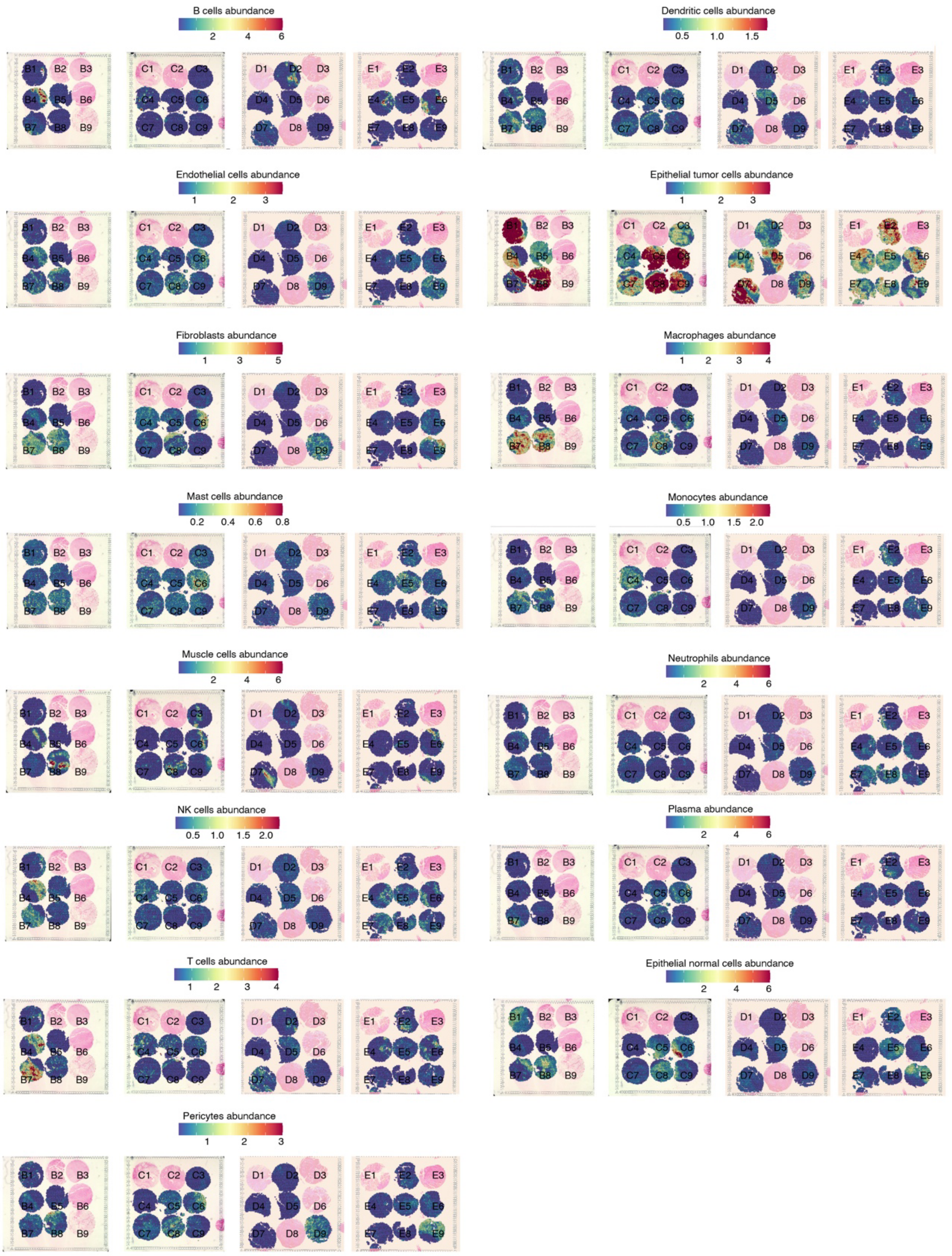


**Supplementary Figure 30: The abundance of different cell types identified from spatial transcriptomics in different HER2 groups.** The HER2 high group includes the following regions: B1, C6, C3, D9, E7, and E8; the HER2 low group includes the following regions: B4, B5, C7, C4, C9, C8, C5, D7, D4, D5, D2, E4, E9, E5, and E6; and the HER2 negative group includes the following regions: B7, B8, and E2.

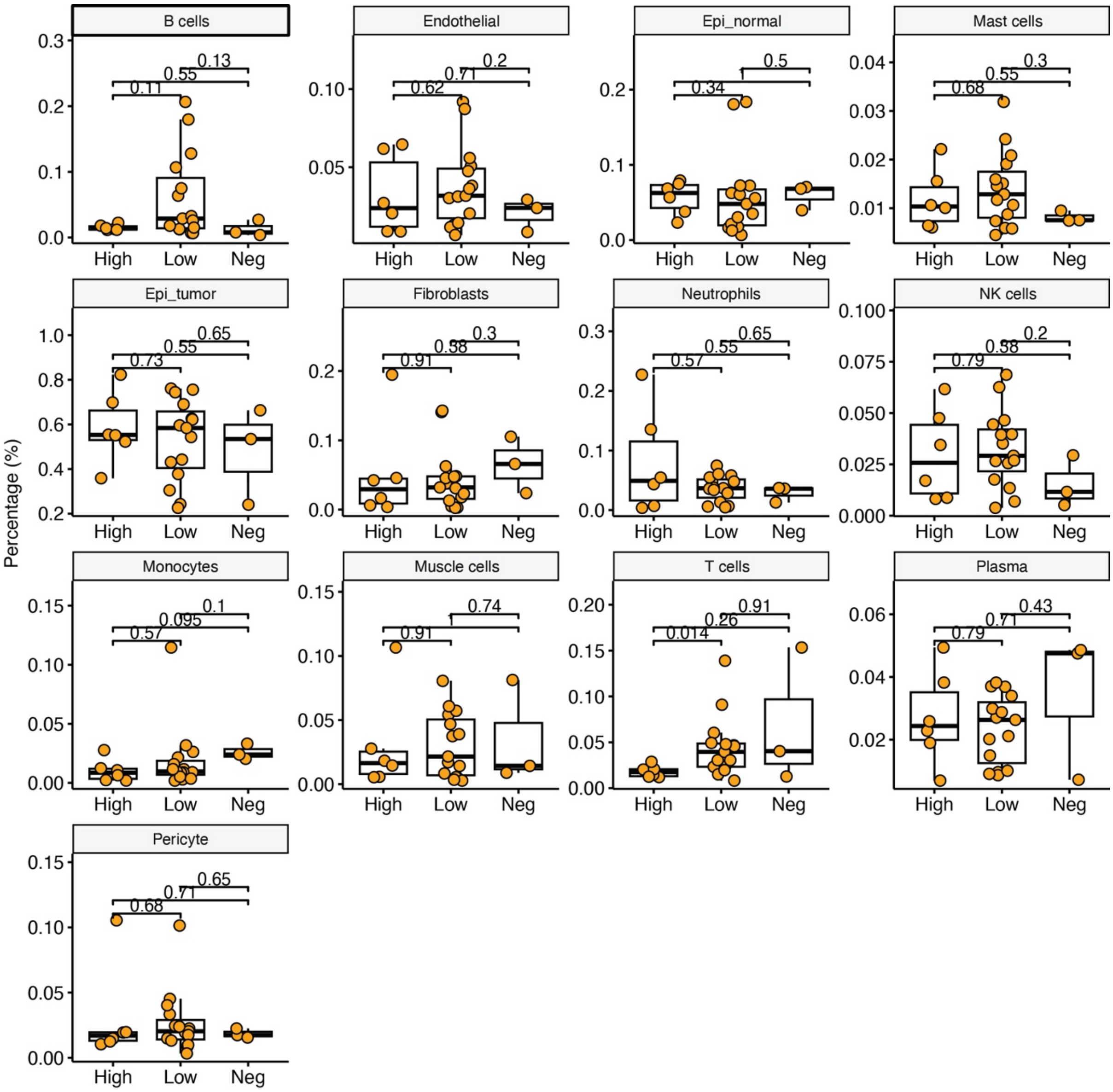


**Supplementary Figure 31: Comparison of different cell types identified from spatial transcriptomics across different HER2 groups.** Neg = Negative group.

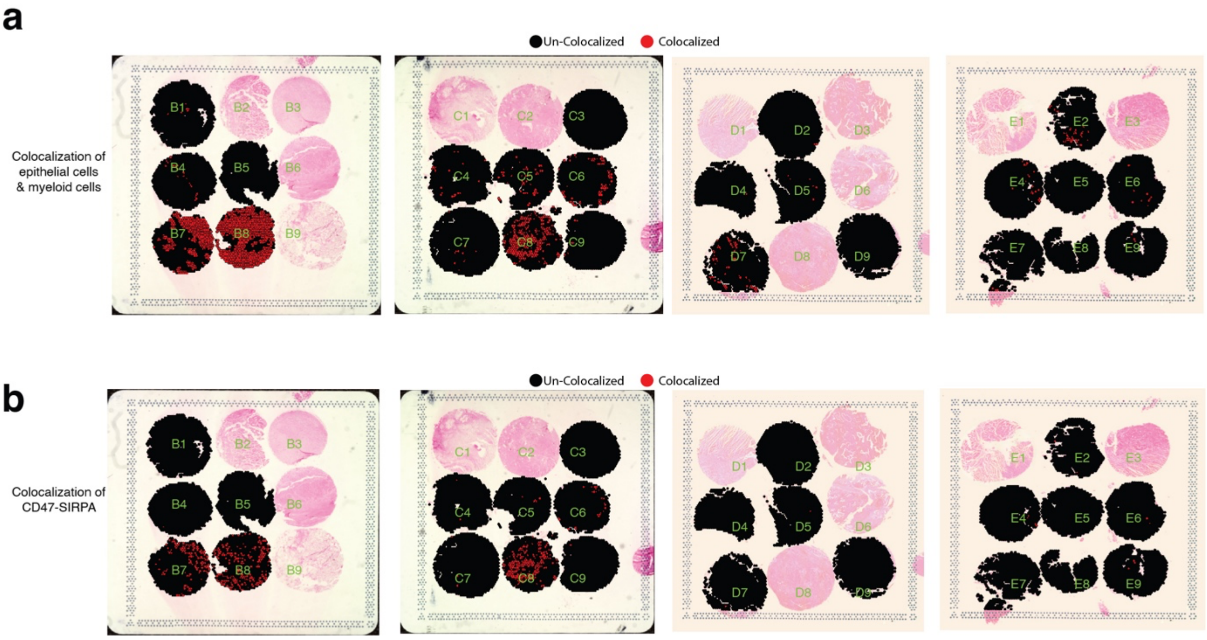


**Supplementary Figure 32: Imaging of co-localization of epithelial cells and myeloid cells (a), and co-localization of CD47-SIRPA in epithelial cells and myeloid cells (b) from spatial transcriptomics.**

**Tables S1-26**

**Table S1:** List of GCA cohort with 128 patients.

**Table S2:** Protein expression matrix of the GCA cohort.

**Table S3:** Uniquely identified proteins for each HER2 group.

**Table S4:** Gene set enrichment analysis of proteins by comparing each HER2 group with the other two groups.

**Table S5:** Hallmark gene sets analysis of protein expression for each HER2 group in GCA patients.

**Table S6:** Differential protein expression in each HER2 group by comparing tumors and their corresponding normal tissue adjacent to the tumor (NAT).

**Table S7:** Gene ontology (GO) terms enrichments from differential protein expression in each HER2 group by comparing tumors and their corresponding normal tissue adjacent to the tumor (NAT).

**Table S8:** Mapping summary of whole exome sequencing (WES) from tumors and their corresponding peripheral blood.

**Table S9:** Mutation frequency detected in genes in different HER2 groups.

**Table S10:** Copy number alteration (CNA) of whole genome with 1Mbp bin size from whole exome sequencing (WES).

**Table S11:** Hallmark gene sets analysis of RNA expression for each HER2 group in GCA patients from tumors only.

**Table S12:** Signature gene list for each HER2 group from the expression of tumors only.

**Table S13:** Hallmark gene sets analysis of RNA expression for each HER2 group by comparing tumors with their corresponding normal tissue adjacent to the tumor (NAT).

**Table S14:** List of differential genes for each group by comparing tumors with their corresponding normal tissue adjacent to the tumor (NAT).

**Table S15:** Group specific and common differentially expressed genes from the comparison of tumors and their corresponding normal tissue adjacent to the tumor (NAT) among the three HER2 groups.

**Table S16:** Gene ontology (GO) terms enrichments from differential RNA expression in each HER2 group by comparing tumors and their corresponding normal tissue adjacent to the tumor (NAT).

**Table S17:** list of exclusive genes in each HER2 group from GO terms of cell cycle, DNA repair, inflammation and major histocompatibility complex (MHC).

**Table S18:** Quantification of CD163 immunofluorescent staining in each HER2 group.

**Table S19:** Differentially expressed genes of M2 macrophages between the HER2 high and low groups.

**Table S20:** Enriched KEGG pathways from differentially expressed genes of M2 macrophages between the HER2 high and low groups.

**Table S21:** Enriched GO (Gene Ontology) pathways from differentially expressed genes of M2 macrophages between the HER2 high and low groups.

**Table S22:** Feature genes for cell fate 1 and cell fate 2 from epithelial tumor cells in GCA patients.

**Table S23:** Enriched KEGG pathways from feature genes for cell fate 1 and cell fate 2 in epithelial tumor cells from GCA patients.

**Table S24:** Quantification of positive cells from immunohistochemical staining of PD-L2 and SIRPA in tumors from different HER2 groups of GCA patients.

**Table S25:** Average strength of immune checkpoint interactions detected from scRNA-Seq in HER2 high and low groups.

**Table S26:** Significance measurement of immune checkpoint interactions detected from scRNA-Seq in HER2 high and low groups.
